# Potent and Selective SETDB1 Covalent Negative Allosteric Modulator Reduces Methyltransferase Activity in Cells

**DOI:** 10.1101/2024.09.27.615363

**Authors:** Mélanie Uguen, Devan J. Shell, Madhushika Silva, Yu Deng, Fengling Li, Magdalena M. Szewczyk, Ka Yang, Yani Zhao, Michael A. Stashko, Jacqueline L. Norris-Drouin, Jarod M. Waybright, Serap Beldar, Justin M. Rectenwald, Angie L. Mordant, Thomas S. Webb, Laura E. Herring, Cheryl H. Arrowsmith, Suzanne Ackloo, Steven P. Gygi, Robert K. McGinty, Dalia Barsyte-Lovejoy, Pengda Liu, Levon Halabelian, Lindsey I. James, Kenneth H. Pearce, Stephen V. Frye

## Abstract

A promising drug target, SETDB1, is a dual Kme reader and methyltransferase, which has been implicated in cancer and neurodegenerative disease progression. To help understand the role of the triple Tudor domain (3TD) of SETDB1, its Kme reader, we first identified a low micromolar small molecule ligand, UNC6535, which occupies simultaneously both the TD2 and TD3 reader binding sites. Further optimization led to the discovery of UNC10013, the first covalent 3TD ligand targeting Cys385 of SETDB1. UNC10013 is potent with a k_inact_/K_I_ of 1.0 x 10^6^ M^-1^s^-1^ and demonstrated proteome-wide selectivity. In cells, negative allosteric modulation of SETDB1-mediated Akt methylation was observed after treatment with UNC10013. Therefore, UNC10013 is a potent, selective and cell-active covalent ligand for the 3TD of SETDB1, demonstrating negative allosteric modulator properties and making it a promising tool to study the biological role of SETDB1 in disease progression.

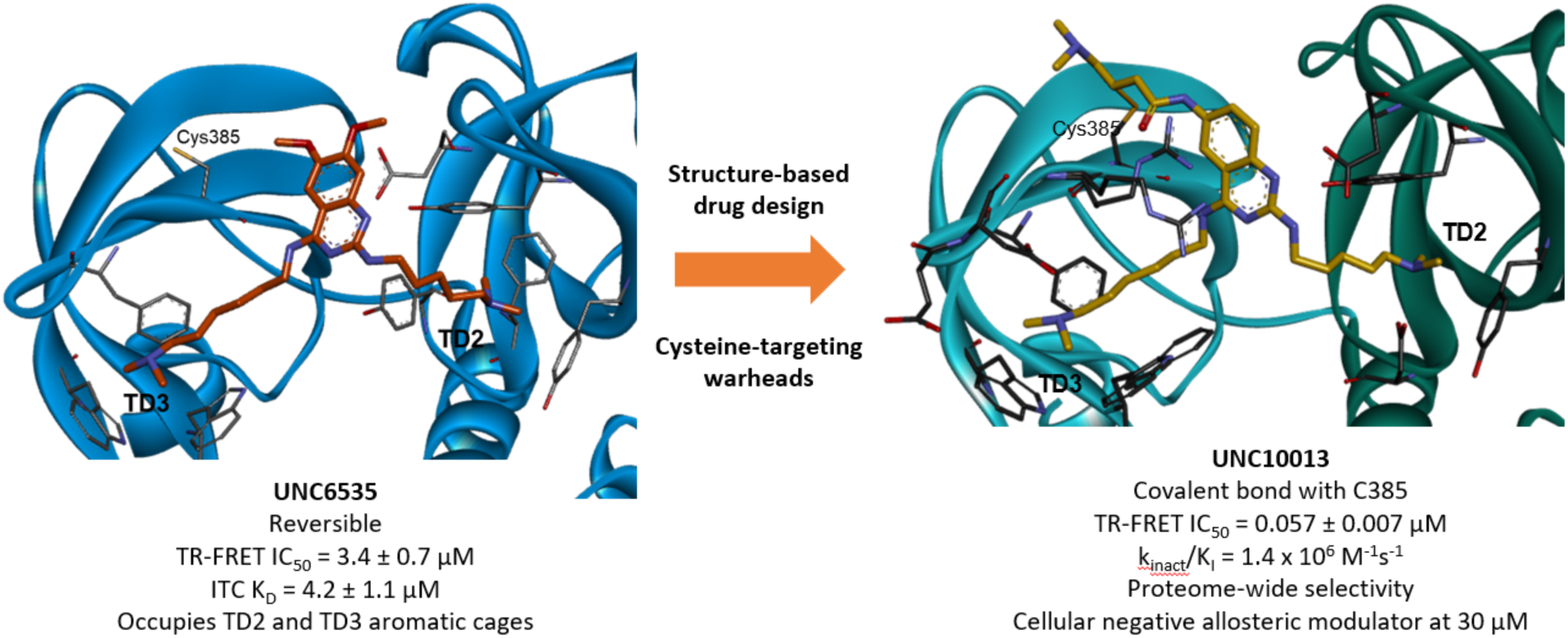

## INTRODUCTION

Histone lysine methylation is a key post-translational modification that regulates control differentiation, development, and gene transcription. Specific lysine methylation motifs on histones are installed by lysine methyltransferases, removed by lysine demethylases, and read by methyl lysine (Kme) readers.^1,2^ Kme reader proteins bind to histones when they exhibit the required methylation levels at a specific lysine residue.^3^ This binding event triggers downstream effects which influence gene expression or repression. Aberrant activity and overexpression of Kme readers can correlate with disease progression, notably in cancer, and although the structural basis for Kme reader to histones is generally well understood, the resulting signaling cascades are less well-studied and would benefit from the development of high-quality chemical probes.^4,5^

SET domain bifurcated 1 (SETDB1) belongs to the Kme reader protein family as it contains three different Tudor domains, forming the triple Tudor domain (3TD), a binding domain for the histone 3 tail when lysine 9 is di- or tri- methylated and lysine 14 is acetylated (H3K9Me2/3K14Ac). The dimethylated lysine of H3K9Me2K14Ac occupies the aromatic cage of the TD3, but when it is trimethylated it can also bind to the aromatic cage of the TD2.^6^ Importantly, the biological consequences of these two binding events are not fully understood.

In addition to its 3TD reader domain, SETDB1 also contains a methyltransferase domain which catalyzes the di- and tri-methylation of lysine 9 of the histone 3 tail as well as the methylation of non-histone proteins such as Akt1, p53 and MCT1.^7–11^ It is currently unclear how the 3TD and the SET domain of SETDB1 interact or if they cooperate in modulating the same signaling cascade.

Recently, a reversible small molecule ligand that binds the Tudor domain 2 (TD2) and the interface of TD2-TD3 of SETDB1, (R,R)-59, and can displace a H3K9Me2 peptide from the 3TD, was reported.^12^ Interestingly, we found that this compound promotes the SETDB1-mediated methylation of Akt1, making it the first positive allosteric modulator for SETDB1; suggestive of cross-talk between the 3TD and SET domains.^13^

Here, we disclose the identification of UNC6535, a novel ligand that reversibly binds both the TD2 and TD3 aromatic cages. Based on the crystal structure of UNC6535, a series of covalent analogs targeting Cys385 were designed and characterized. Among them, UNC10013 is a potent and selective covalent ligand for the 3TD, with a k_inact_/K_I_ of 1.0 x 10^6^ M^-1^s^-1^, demonstrating proteome-wide selectivity, and which showed no significant direct inhibition of the methyltransferase activity of SETDB1 *in vitro*. However, in cells, UNC10013 decreases the ability of SETDB1 to methylate the Akt kinase in a dose-dependent manner through an allosteric mechanism. Overall, this study successfully identified the first potent and selective negative allosteric modulator for SETDB1-mediated Akt methylation.

## RESULTS

### Identification of reversible ligand UNC6535

Screening of an in-house Kme reader-targeted library of small molecules by a time-resolved fluorescence resonance energy transfer (TR-FRET) detection of the displacement of an H3K9Me2K14Ac peptide^13^ revealed UNC6535 as a 3TD ligand with an IC_50_ of 3.4 ± 0.7 µM (Fig. 1a). An orthogonal isothermal titration calorimetry assay confirmed that UNC6535 binds SETDB1 3TD with a K_d_ of 4.2 ± 1.1 µM. UNC6535 also demonstrated the ability to weakly inhibit SETDB1 methyltransferase activity in an *in vitro* radiometric methyltransferase assay^13^ with an IC_50_ of 84 ± 7.2 µM for the full-length protein (SETDB1-FL, aa 1-1291) and 30 ± 2 µM on a SETDB1 construct (SETDB1-S, aa 570-1291) that does not contain the 3TD. A fluorescence polarization (FP) displacement assay showed an IC_50_ of 8.6 ± 0.5 µM for the displacement of FITC-H3 (1-25) from SETDB1-S by UNC6535. This shows that UNC6535 lacks selectivity for the 3TD and may also interact with the SET domain.

**Figure 1.**
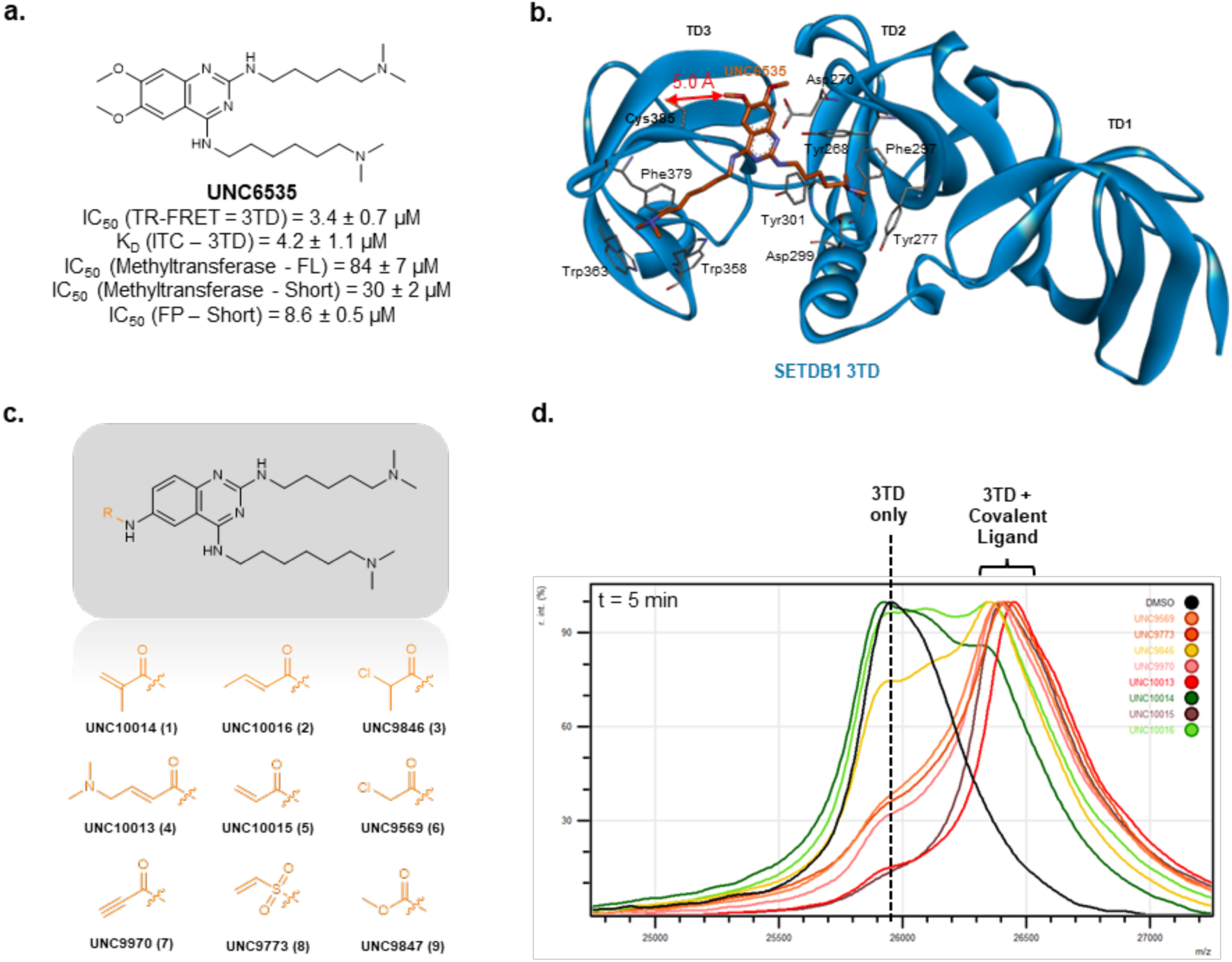
Structure-based design of UNC6535 analogs aimed at covalently binding to the 3TD. (a) Structure and characterization of UNC6535. Reported IC_50_ and K_d_ values are the average of at least two independent experiments ± s.d. (b) Crystal structure of UNC6535 bound to SETDB1 3TD (PDB: 8G5E). (c) Structure of nine covalent analogues of UNC6535 **1**-**9** that were synthesized and tested. (d) Representative MALDI-TOF qualitative evaluation of the ability of **1**-**9** to covalently modify SETDB1 3TD (1:1 ratio) after 5 min incubation on ice.

A crystal structure of UNC6535 bound to the 3TD of SETDB1 shows the formation of cation-pi interactions with the aromatic cages of TD2 and TD3 (Fig. 1b). The dimethylamine chain from position 2 of the quinazoline forms cation-pi interactions with Tyr268, Tyr277, and Phe297 and a salt bridge with Asp299 from the TD2 aromatic cage. The second dimethylamine chain forms cation-pi interactions with Trp358, Trp363, and Phe379 of the TD3 aromatic cage. Other favorable interactions include pi-pi stacking between the quinazoline core and Tyr301 and a hydrogen bond between the nitrogen atom at position 1 of the quinazoline and Asp270. Interestingly, UNC6535 is the first small molecule reported to occupy both TD aromatic cages simultaneously.

### Design and Synthesis of 3TD Covalent Ligands Targeting Cys385

The crystal structure of UNC6535 and the 3TD of SETDB1 revealed that the 6-methoxy group is 5.0 Å from Cys385 while the 7-methoxy moiety is solvent-exposed (Fig. 1b). Therefore, to improve the binding affinity of UNC6535 and its selectivity for the 3TD over the SET domain of SETDB1, a set of nine analogs containing cysteine-targeting electrophilic warheads at position 6 of the quinazoline (**1**-**9**) was designed and synthesized (Fig. 1c).

To evaluate the ability of compounds **1**-**9** to covalently modify SETDB1, the compounds were incubated with SETDB1 3TD protein (10:1 ratio, 2 h incubation at room temperature) and tested by matrix-assisted laser desorption/ionization – time-of-flight (MALDI-TOF) mass spectrometry. Encouragingly, eight out of the nine compounds were able to covalently modify the 3TD of SETDB1, only the methylcarbamate warhead-containing compound (**9**) did not show any covalent modification of SETDB1 (Extended Data Fig. 1). This might be explained by a longer distance between the electrophilic site of the methylcarbamate and the cysteine as compared to the eight other warheads. Thus, eight covalent analogs of UNC6535 were successfully synthesized and demonstrated the ability to covalently modify SETDB1 3TD by mass spectrometry.

### Comparison of Eight 3TD Covalent Hits Identified for SETDB1 3TD

Further characterization of the compounds **1**-**8** was performed to select the most promising hits for further evaluation. A time-dependent MALDI-TOF experiment was conducted to qualitatively evaluate the ability of these compounds to covalently react with the 3TD. After 5 minutes of incubation with 3TD on ice, all compounds show partial, if not full, covalent bond formation allowing for the qualitative ranking of the compounds for their reactivity towards the 3TD of SETDB1 (Fig. 1d). Warheads such as propiolamide (**7**) or vinyl sulfonamide (**8**) showed full covalent modification of the 3TD under these conditions, likely due to their higher intrinsic reactivity. When methylated at the α or β position, acrylamides (**1** and **2**), and chloroacetamides (**3**) showed slightly reduced reactivity towards the 3TD compared to the unmethylated (**5** and **6**) or dimethylamino-modified analogues (**4**). However, after 30 minutes of incubation on ice, all of the compounds completely covalently labelled the 3TD demonstrating a high reactivity of all compounds for the 3TD of SETDB1 (Extended Data Fig. 2).

The intrinsic reactivity of the covalent 3TD ligands was evaluated using glutathione (GSH) and was quantified by liquid chromatography–mass spectrometry (LC-MS). Molecules with high intrinsic reactivity are more likely to lack specificity for the target of interest and form covalent bonds with off-target proteins containing reactive cysteines.^14^ Results showed that the eight covalent SETDB1 ligands had a half-life between 9 minutes and over 10 days (Table 1). Compounds with t_1/2_ below 2 hours were considered to have risks of off-target reactions and were therefore deprioritized. This led to four promising compounds: UNC10014 (**1**), UNC10016 (**2**), UNC9846 (**3**), and UNC10013 (**4**).

**Table 1.**
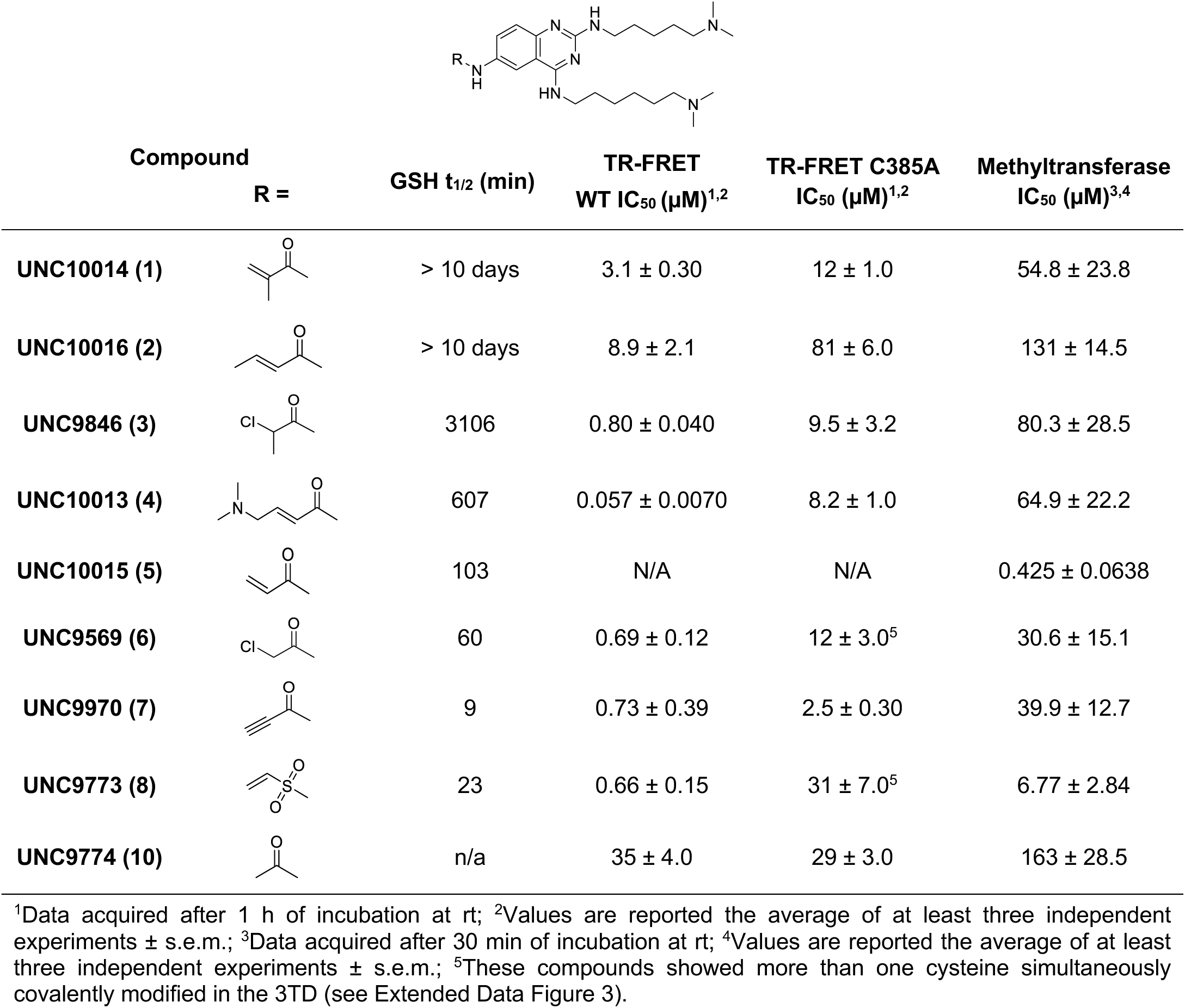
Evaluation of the intrinsic reactivity, binding affinity, and inhibition properties of compounds 1-10.

| Compound | R = | GSH $t_{1/2}$ (min) | TR-FRET<br>WT $IC_{50}$ (μM)<sup>1,2</sup> | TR-FRET C385A<br>$IC_{50}$ (μM)<sup>1,2</sup> | Methyltransferase<br>$IC_{50}$ (μM)<sup>3,4</sup> |
| --- | --- | --- | --- | --- | --- |
| UNC10014 (1) | | > 10 days | $3.1 \pm 0.30$ | $12 \pm 1.0$ | $54.8 \pm 23.8$ |
| UNC10016 (2) | | > 10 days | $8.9 \pm 2.1$ | $81 \pm 6.0$ | $131 \pm 14.5$ |
| UNC9846 (3) | | 3106 | $0.80 \pm 0.040$ | $9.5 \pm 3.2$ | $80.3 \pm 28.5$ |
| UNC10013 (4) | | 607 | $0.057 \pm 0.0070$ | $8.2 \pm 1.0$ | $64.9 \pm 22.2$ |
| UNC10015 (5) | | 103 | N/A | N/A | $0.425 \pm 0.0638$ |
| UNC9569 (6) | | 60 | $0.69 \pm 0.12$ | $12 \pm 3.0^5$ | $30.6 \pm 15.1$ |
| UNC9970 (7) | | 9 | $0.73 \pm 0.39$ | $2.5 \pm 0.30$ | $39.9 \pm 12.7$ |
| UNC9773 (8) | | 23 | $0.66 \pm 0.15$ | $31 \pm 7.0^5$ | $6.77 \pm 2.84$ |
| UNC9774 (10) | | n/a | $35 \pm 4.0$ | $29 \pm 3.0$ | $163 \pm 28.5$ |

<sup>1</sup>Data acquired after 1 h of incubation at rt; <sup>2</sup>Values are reported the average of at least three independent experiments ± s.e.m.; <sup>3</sup>Data acquired after 30 min of incubation at rt; <sup>4</sup>Values are reported the average of at least three independent experiments ± s.e.m.; <sup>5</sup>These compounds showed more than one cysteine simultaneously covalently modified in the 3TD (see Extended Data Figure 3).

In parallel, all compounds were tested in a TR-FRET assay to examine their ability to displace the endogenous ligand of the 3TD, H3K9Me2K14Ac (1-25) peptide. Compounds were incubated for 1 h at room temperature before quantification of the FRET signal. All compounds exhibited IC_50_s between 0.057 and 8.9 µM, except UNC10015 (**5**) which increased the TR-FRET signal over the control limit preventing the quantification of the H3K9Me2K14Ac displacement (Table 1). UNC10013 (**4**) stood out from the other compounds with a 10-fold potency increase compared to UNC9970 (**7**), the second most potent ligand. The different compounds were also tested in the same TR-FRET assay using a cysteine 385 to alanine (C385A) mutant SETDB1 3TD protein. The binding affinities of all covalent compounds decreased compared to the wild-type protein, confirming the role of the covalent bond formation with Cys385 in the binding mode of these compounds (Table 1).

After confirming that all eight covalent 3TD ligands are efficient at forming a covalent bond with SETDB1, and because the 3TD of SETDB1 contains three different cysteines, Cys314, Cys329, and Cys385, we wanted to evaluate the stoichiometry of the covalent adducts. All the compounds were incubated with the 3TD and submitted for high resolution intact protein mass spectrometry which is more sensitive and accurate than MALDI-TOF for larger proteins such as the 3TD of SETDB1. Results confirmed that compounds **1**-**8** were fully covalently modifying the 3TD of SETDB1, reinforcing the high reactivity of these small molecules for the 3TD (Extended Data Fig. 3). The main peak for each sample corresponded to the protein modified by one copy of ligand, although compounds UNC9569 (**6**) and UNC9773 (**8**) showed minor peaks for the mass corresponding to modification by two ligands. It should be noted that the relative intensity of these peaks was below 6% compared to the intensity of the main peak for the modification of the protein by one copy of ligand and that the protein/ligand concentration ratio was 1:10 which suggests that these 2:1 binding modes could be attributed to the high concentration of ligand (Extended Data Fig. 3). Overall, limited concern about unspecific reactivity for other cysteines of the 3TD was raised.

Compiling all of the data, all compounds were very efficient at forming covalent bonds with the 3TD of SETDB1 and displacing the H3K9Me2K14Ac peptide. However, only four compounds showed GSH t_1/2_ below 2 hours, UNC10014 (**1**), UNC10016 (**2**), UNC9846 (**3**) and UNC10013 (**4**), suggesting reduced potential for off-target covalent bond formation compared to the other four compounds. The substitution of acrylamide or chloroacetamide warheads with methyl or dimethylaminomethyl groups reduced the warhead reactivity but maintained the required geometry necessary for the formation of the covalent bond. Therefore, UNC10014 (**1**), UNC10016 (**2**), UNC9846 (**3**) and UNC10013 (**4**) were selected for further characterization.

### Characterization of Four Lead Covalent Ligands for SETDB1 3TD

As mentioned earlier, the 3TD of SETDB1 contains three different cysteines, Cys314, Cys329 and Cys385. We performed LC-MS/MS to identify the cysteines which were covalently modified by the ligands on purified 3TD. Results showed that all four ligands, UNC10014 (**1**), UNC10016 (**2**), UNC9846 (**3**) and UNC10013 (**4**), modified Cys385 as expected. UNC9846 (**3**) also showed the ability to covalently modify Cys329 but to a smaller extent than Cys385.

The four best compounds were also evaluated in the SETDB1 methyltransferase activity assay, following three different pre-incubation times of 0, 1, and 2 hours. Surprisingly, UNC10014 (**1**) was the only compound with an IC_50_ dependent on the pre-incubation time (Extended Data Fig. 4). One hypothesis is that UNC10014 (**1**) is forming a covalent bond with another cysteine among the 36 cysteines of SETDB1 leading to inhibition of the methyltransferase activity. Nevertheless, the three other compounds also exhibited inhibitory properties for the methyltransferase activity of SETDB1 with IC_50_ between 21 and 42 µM but did not display a dependence on the pre-incubation time, consistent with a reversible mechanism.

To quantitate the targeted covalent reactivity of our best ligands, k_inact_, the irreversible reaction rate, and K_I_, the reversible inhibition constant of the ligand, were assessed for UNC10016 (**2**), UNC9846 (**3**), and UNC10013 (**4**), adapting the previously described TR-FRET system. We examined the time and concentration dependence of our ligands competition with the endogenous ligand, H3K9Me2K14Ac, and hypothesized that the prevention of binding of the peptide by the tested ligand was a consequence of the formation of the covalent bond with Cys385, as previous mass spectrometry data confirmed that the compounds were efficient at covalently modifying the Cys385 of the 3TD, and the assay was performed with dilution to concentrations below the apparent IC_50_ of the compounds to minimize the reversible binding contribution. For all three compounds, k_inact_s were comparable (k_inact_ = 2.61-6.22 x 10^-2^ s^-1^), suggesting similar abilities to form covalent bonds for the three compounds. The differences in K_I_ were surprising with UNC10013 (**4**) exhibiting a K_I_ of 60 ± 1.0 nM while UNC9846 (**3**) and UNC10016 (**2**) K_I_s were respectively 0.75 ± 0.35 µM and 7.9 ± 3.4 µM (Fig. 2a). This low K_I_ made UNC10013 (**4**) a promising lead compound with a k_inact_/K_I_ of 1.04 x 10^6^ M^-1^s^-1^, in the same range as FDA-approved covalent drugs.^15^

**Figure 2.**
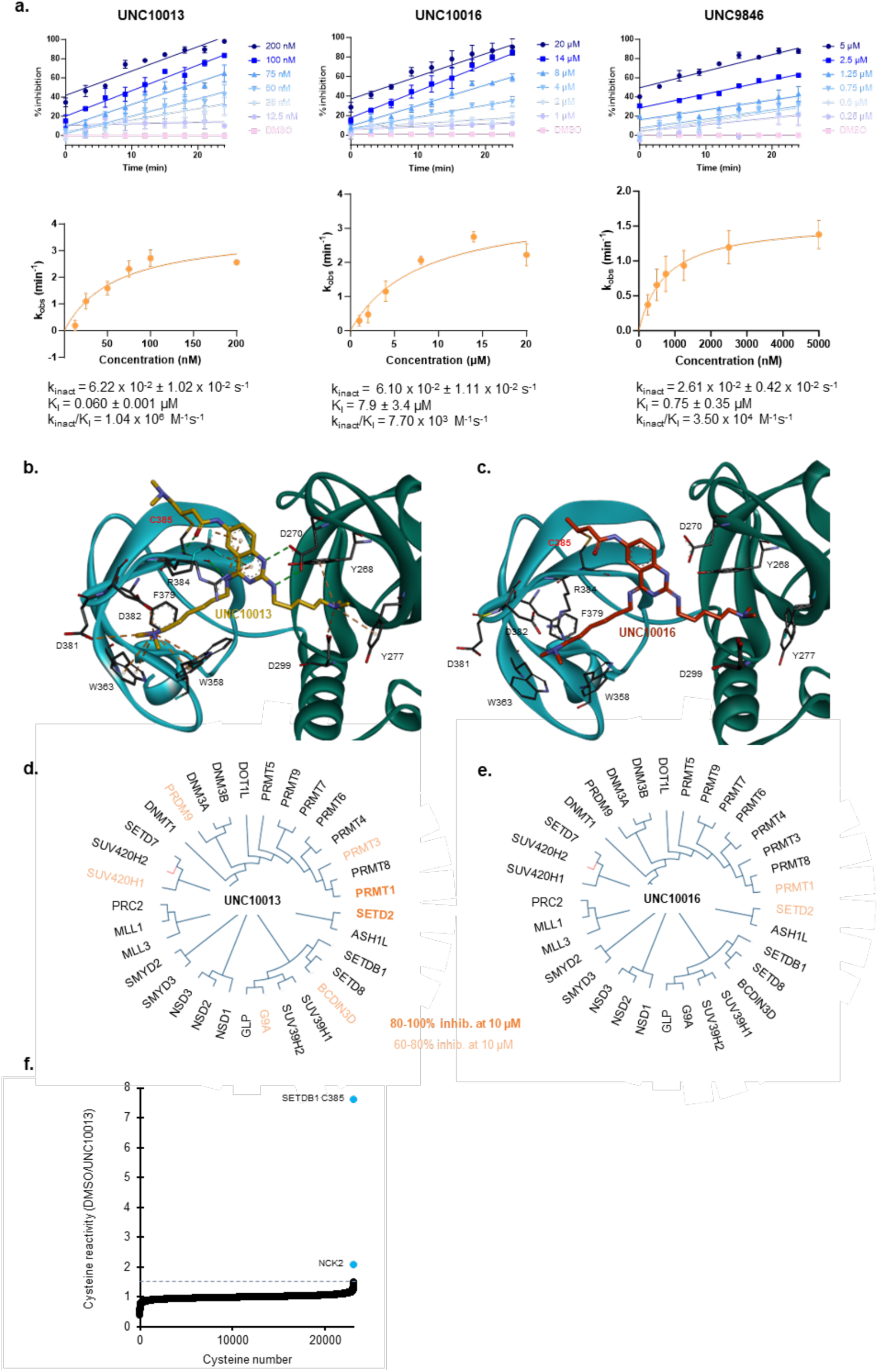
Covalent chemical probe criteria evaluation for UNC10016 (2), UNC9846 (3) and UNC10013 (4). (a) Kinetic parameters k_inact_/K_I_ evaluation for UNC10016 (**2**), UNC9846 (**3**), and UNC10013 (**4**) using a TR-FRET displacement assay. The inhibition at different time points for different concentrations of compound were measured and plotted as a function of time. Resulting slopes, corresponding to k_obs_ were plotted as a function of the concentration to calculate k_inact_ and K_I_ values. Reported values are the average of at least three independent experiments ± s.e.m. (b) Crystal structure of UNC10013 (**4**) (left) and (c) UNC10016 (**2**) (right) bound to SETDB1 3TD (PDB ID: 9CUX, 9CUW) Visualized with Discovery Studio. Cation-pi interactions are shown in orange. H-bond interactions are shown in green. (d) Phylogenetic tree of the 33 methyltransferases tested for off-target activity of UNC10013 (**4**) and (e) UNC10016 (**2**) using a radiometric methyltransferase assay. Values are the average of at least three independent experiments. Proteins in light orange showed 60-80% inhibition by tested compounds at 10 µM. Proteins in dark orange showed over 80% inhibition by tested compounds at 10 µM (see Supporting Information Table S1). (f) Plot showing the relative cysteine reactivity of 23,149 detected cysteines after treatment of MCF-7 cell lysate with 10 µM of UNC10013 (**4**) compared to DMSO. Two cysteines showed significant cysteine reactivity, SETDB1 Cys385 and NCK2 Cys144 (>1.5).

After further characterization of the four lead covalent ligands, UNC10016 (**2**) and UNC10013 (**4**) were deemed more attractive for further development as they did not covalently modify other cysteines of SETDB1 thanks to lower intrinsic reactivities, unlike UN9846 (**3**) and UNC10014 (**1**), while showing potent k_inact_/K_I_ parameters.

### Orthogonal Binding Evaluation of UNC10016 (2) and UNC10013 (4)

Three assays were utilized for the orthogonal binding evaluation of UNC10016 (**2**) and UNC10016 (**4**): fluorescence polarization displacement assay, differential scanning fluorimetry (DSF), and a TR-FRET-based nucleosome displacement assay.^16^ Both UNC10016 (**2**) and UNC10013 (**4**) showed potent ability to displace the H3K9Me2K14Ac peptide in the FP assay, with respective IC_50_s of 0.15 and 0.07 µM at t = 1 h, validating the compounds as potent 3TD ligands (Table 2).

**Table 2.**
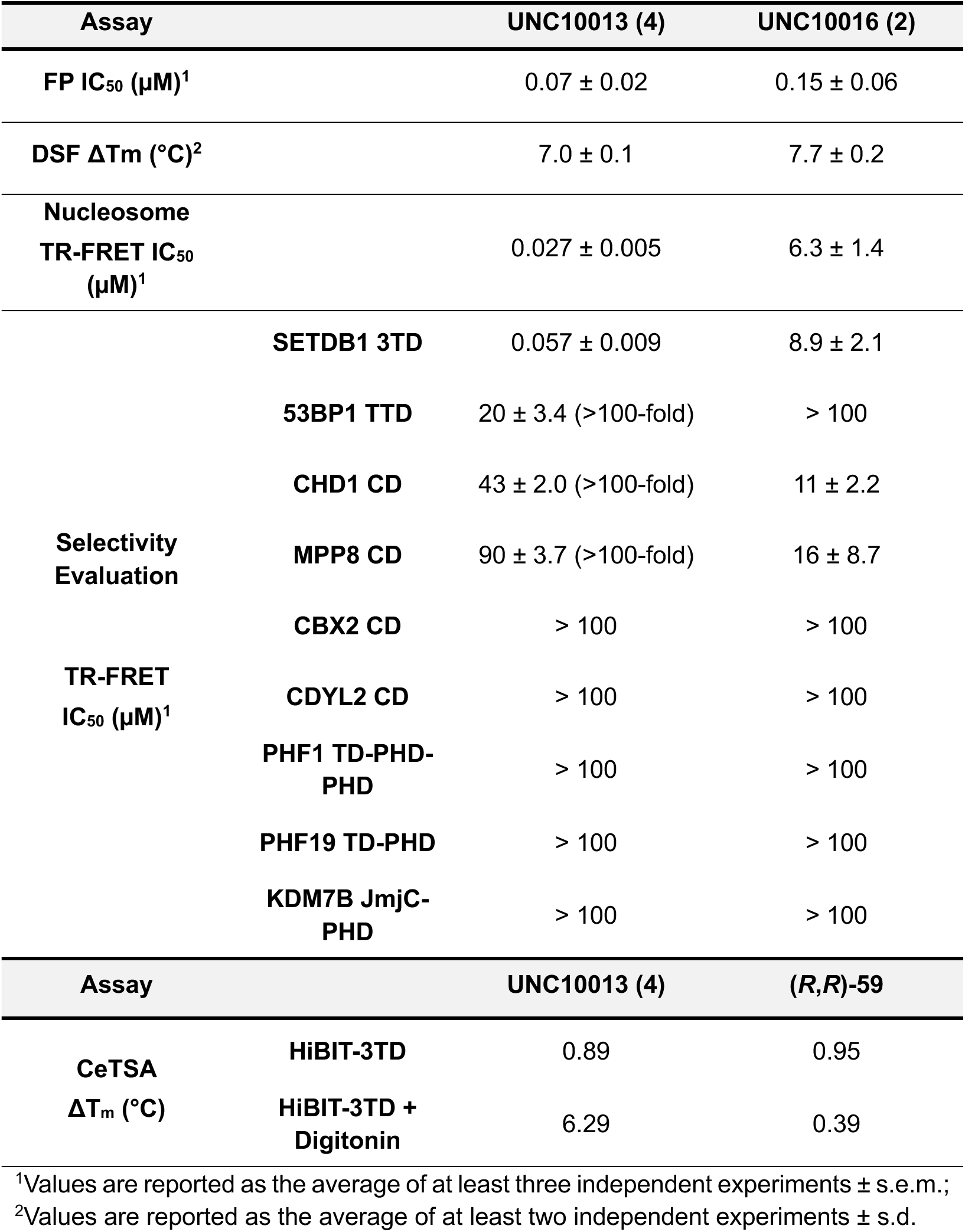
Biological evaluation of UNC10013 and UNC10016.

DSF evaluation of the ability of ligands to stabilize the 3TD showed significant temperature shifts of 7.0 and 7.7 °C for UNC10013 (**4**) and UNC10016 (**2**) respectively (Table 2). This confirmed the ability of the compounds to bind the 3TD.

Finally, since our prior assays looked at the displacement of H3 tail-mimicking peptides while a natural ligand, the nucleosome, could include different binding components, we examined the 3TD binding to a H3K9Me2K14Ac-modified nucleosome using TR-FRET detection.^16^ In this assay, we showed that UNC10013 (**4**) and UNC10016 (**2**) were able to displace the full nucleosome from the 3TD. UNC10013 (**4**) showed a potent IC_50_ of 27 nM, which is similar to the peptide TR-FRET IC_50_ of 57 nM while UNC10016 (**2**) was less potent with an IC_50_ of 6.3 µM (peptide TR-FRET IC_50_ = 8.9 µM) (Table 2). These results enhanced our interest in further developing UNC10013 (**4**) and UNC10016 (**2**) into covalent chemical tools to study the interaction of the 3TD with the nucleosome and other substrates.

### Covalent Chemical Probe Criteria Evaluation for UNC10016 (2) and UNC10013 (4) Candidates

Chemical probe experts have agreed on the criterion required for a covalent chemical probe:^17^

1. Biophysical proof of target engagement
2. Known k_inact_/K_I_
3. Target family selectivity (> 30-fold)
4. Proteome-wide selectivity
5. Availability of inactive control
6. Cellular potency (< 1 µM)

Promising results from the characterization of UNC10016 (**2**) and UNC10013 (**4**) led to the evaluation of both ligands as chemical probe candidates applying each of the six criteria. One of the criteria, knowing the k_inact_/K_I,_ being already addressed in our initial studies.

#### Co-crystallization of UNC10016 (2) and UNC10013 (4) in the 3TD

Co-crystallization of UNC10016 (**2**) and UNC10013 (**4**) within the 3TD confirmed that both compounds maintained binding poses similar to UNC6535 while covalently engaging Cys385 (Fig. 2b and 2c). New interactions between the backbone of Cys385 and the carbonyl of UNC10013 (**4**) and UNC10016 (**2**) were observed while only the UNC10013 (**4**) quinazoline core showed some cation-pi interactions with Arg384 which might explain its improved K_I_. Overall, the X-ray crystallography data met criterion 1 for a covalent chemical probe while highlighting the unique ability of these compounds to occupy the aromatic cages of both TD2 and TD3 of the 3TD of SETDB1 simultaneously while covalently modifying Cys385.

#### Selectivity evaluation: family and proteome-wide

A screen of UNC10016 (**2**) and UNC10013 (**4**) against various Kme reader proteins using similar TR-FRET technology as for SETDB1 was initiated.^18^ Results showed that UNC10013 (**4**) was at least 100-fold selective against the eight Kme readers tested, which included various Kme sub-families such as Tudor domains, chromodomains, and PHD domains (Table 2). Unfortunately, UNC10016 (**2**) lacked selectivity against the chromodomains of CHD1 and MPP8.

SETDB1 also contains a lysine methyltransferase domain which modifies substrates similar to the 3TD ligands, so we expanded the family selectivity evaluation to the methyltransferase family. The ability of thirty-three methyltransferases to modify lysine and arginine-containing substrates was tested in the presence of a single concentration of 10 µM of UNC10013 (**4**) and UNC10016 (**2**). UNC10013 (**4**). Confirming our previous results, no inhibition of the SETDB1 methyltransferase activity was observed at 10 µM. This assay also showed at least 60% inhibition for seven methyltransferases, two of them, SETD2 and PRMT1, showed over 80% inhibition at 10 µM (Fig. 2d, 2e). Accordingly, a full dose-response evaluation of the ability of UNC10013 (**4**) to inhibit SETD2 and PRMT1 methyltransferase activities was performed. UNC10013 (**4**) is selective for the binding of the 3TD of SETDB1 by 30-fold relative to SETD2 inhibition (IC_50_ = 0.8 ± 0.1 µM), and 48-fold relative to PRMT1 inhibition (IC_50_ = 1.3 ± 0.1 µM). Although this data suggests that UNC10013 marginally meets the covalent chemical probe criterion for selectivity (> 30-fold), it should be noted that UNC10013 is covalently modifying the 3TD of SETDB1 while reversibly inhibiting the methyltransferase activities of SETD2 and PRMT1 (Fig. 2f).

Regarding UNC10016 (**2**), while fewer methyltransferases were inhibited with the compound at 10 µM, both SETD2 and PRMT1 showed over 60% inhibition suggesting similar binding affinities compared to the 3TD (3TD IC_50_ = 6.3 ± 1.4 µM). Based on the lack of selectivity for both the Kme reader and methyltransferase proteins, UNC10016 was deprioritized for development as a covalent chemical probe.

Cysteine-targeted activity-based protein profiling (ABPP) was used to evaluate the proteome-wide selectivity of UNC10013 (**4**). Results showed that out of 23,149 detected cysteines in the native MCF-7 proteome, remarkably UNC10013 (**4**) preferentially modified SETDB1 at Cys385 relative to 22 other cysteines also present in SETDB1 (Fig. 2f and Table S2), and only one off-target NCK2 (Cys144) was detected.

Overall, characterization of the selectivity of UNC10013 (**4**) and UNC10016 (**2**) demonstrated that UNC10013 (**4**) has a promising selectivity profile as a covalent chemical probe candidate. We identified two methyltransferase proteins as potential off-targets via a reversible binding mode, which suggests that cellular applications should proceed with consideration of methyltransferase inhibition as a possible confounding activity.

#### Inactive control development

To facilitate cellular studies, a negative control analog that retains the covalent warhead of UNC10013 (**4**) was developed. One strategy was to replace the TD2-occupying basic center with neutral heteroatom-containing groups to prevent the key cation-pi interaction with the TD2 aromatic cage. Therefore, the dimethylamine group was replaced by a methoxy (UNC11277 (**11**)) or an acetamide group (UNC11366 (**12**)) by adapting the synthetic route of UNC10013 (Fig. 3a).

**Figure 3.**
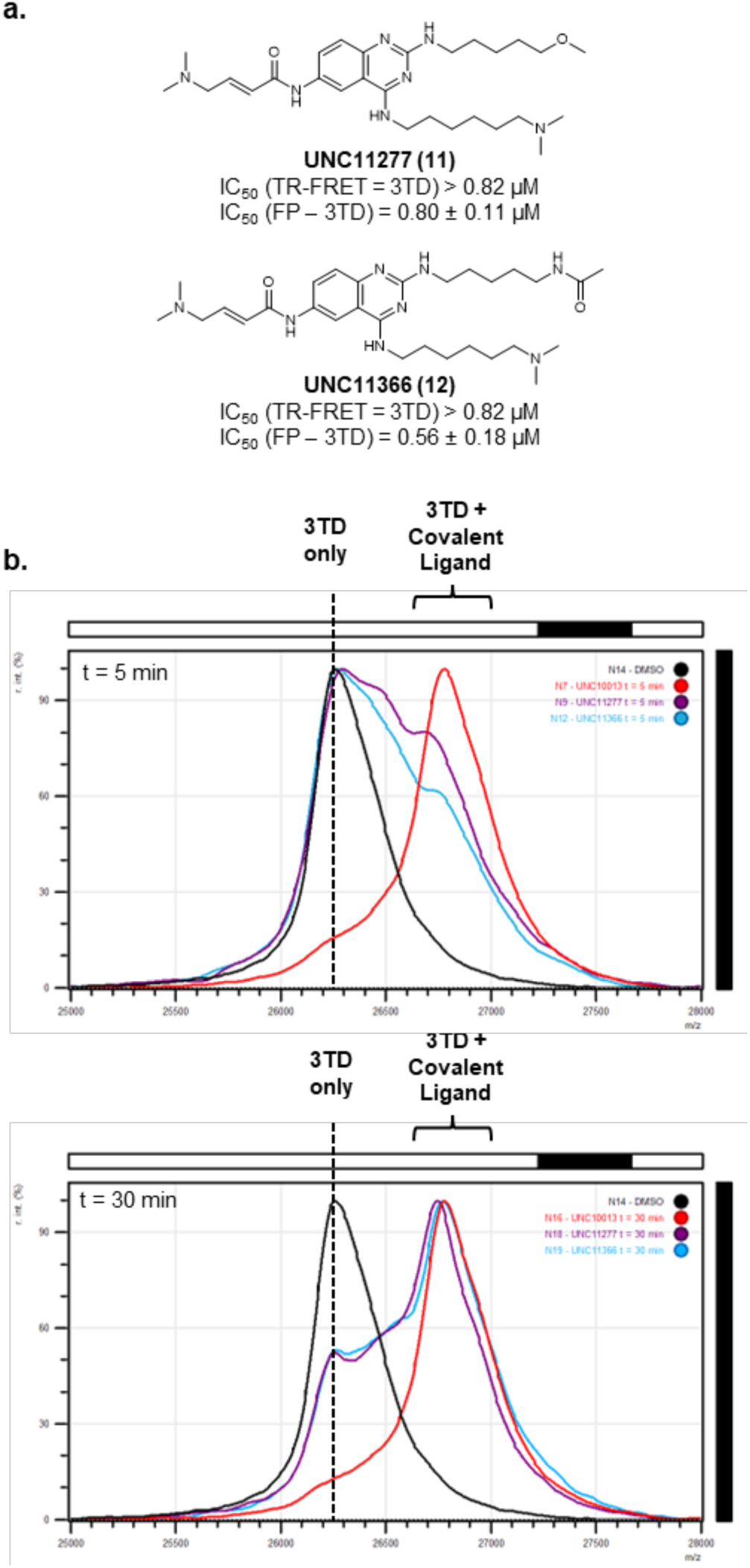
Characterization of negative control candidates UNC11277 (11) and UNC11366 (12). (a) Structure and binding evaluation of UNC11277 (**11**) and UNC11366 (**12**). Reported values are the average of at least three independent experiments ± s.e.m. (b) Representative MALDI-TOF qualitative evaluation of the ability of UNC10013 (**4**) (red) UNC11277 (**11**) (purple), and UNC11366 (**12**) (blue) to covalently modify SETDB1 3TD after t = 5 min (top) and t = 30 min (bottom) incubation on ice.

The two negative controls were evaluated using MALDI-TOF for their ability to form a covalent bond with SETDB1 3TD. Results showed that both compounds **11** and **12** retained the ability to form a covalent bond with the 3TD but at a slower rate than UNC10013 (**4**). This confirmed a decrease in affinity after removal of the basic center in the TD2. After 5 minutes of incubation on ice, most 3TD remained unmodified by UNC11277 (**11**) and UNC11366 (**12**) while UNC10013 (**4**) completely covalently modified the protein (Fig. 3b). While compounds **1**-**8** showed complete covalent labelling after 30 minutes (Extended Data Fig. 2), UNC11277 (**11**) and UNC11366 (**12**) showed partial covalent labelling suggesting a lower reactivity towards the covalent modification of SETDB1 (Fig. 3b). After 2 hours of incubation on ice, both negative control candidates **11** and **12** showed a nearly complete covalent modification of SETDB1, confirming their ability to covalently modify the 3TD of SETDB1 (Extended Data Fig. 5).

In order to quantify their reactivity for SETDB1, both compounds **11** and **12** were tested in the TR-FRET and FP displacement assays. UNC11277 (**11**) and UNC11366 (**12**) showed an IC_50_ > 0.8 µM, providing over 54-fold selectivity in the TR-FRET assay compared to UNC10013 (**4**) (Fig. 3a). However, in the FP assay, UNC11277 (**11**) and UNC11366 (**12**) showed a 10-fold (IC_50_ = 0.80 µM) and a 7-fold selectivity (IC_50_ = 0.56 µM) respectively (Fig. 3a). These results suggest that UNC11277 (**11**) is a promising negative control with lower reactivity towards the covalent modification of the 3TD of SETDB1 than UNC10013 (**4**) and decreased binding ability.

#### Cellular target engagement evaluation

One key characteristic of chemical probes is their ability to potently engage the target of interest in cells. First, a cell viability evaluation was performed to ensure UNC10013 (**4**) and its negative control, UNC11277 (**11**), were not toxic to the cells. Treatment of MCF-7 cells with concentrations up to 50 µM for 24h, 48h, and 72h did not result in a cell viability decrease compared to DMSO (Extended Data Fig. 6).

Another key parameter to ensure that small molecules can engage the target of interest in cells is cell permeability. To evaluate the cell permeability of UNC10013 (**4**), a Caco-2 evaluation was performed. Although no concerning efflux ratio was measured (Efflux ratio = 1.11), a low apparent permeability P_app_ (A-B) was observed (P_app_ (A-B) = 0.37 x 10^-6^ cm/s; P_app_ (B-A) = 0.41 x 10^-6^ cm/s) suggesting modest cell permeability. This result is likely associated with the fact that UNC10013 (**4**) contains three basic amines and three H-bond donor groups.

The cellular thermal shift assay (CeTSA) was used to evaluate target engagement of UNC10013 (**4**). HEK293T cells were transfected with SETDB1 3TD tagged with HiBIT and incubated with 30 µM of UNC10013 (**4**) or (R,R)-59, a known 3TD cellular ligand, used as a positive control. Similar ΔT_m_ were measured for both UNC10013 (**4**) and the positive control (R,R)-59, 0.89, and 0.95 °C respectively, although the thermal shifts remained small (Table 2). Concerns about the low cell permeability of UNC10013 led us to reproduce the CeTSA results using cells permeabilized by digitonin. Indeed, a ΔT_m_ of 6.29 °C was measured under this condition, similar to the *in vitro* ΔT_m_ for UNC10013 (**4**) and the 3TD (Table 2). The significant increase of ΔT_m_ upon permeabilization of the cells confirms that UNC10013 has low cell permeability decreasing its potency in cellular assays. Surprisingly, (R,R)-59 treatment of digitonin-permeabilized cells showed a decrease in ΔT_m_ compared to unpermeabilized cells potentially possibly due to cellular toxicity. Overall, we confirmed that UNC10013 (**4**) is able to engage the 3TD in cells at 30 µM although this ability is impaired due to low cell permeability.

#### Covalent chemical probe summary

To sum up, UNC10013 (**4**) has shown potent k_inact_/K_I_ kinetic parameters for covalently binding the 3TD of SETDB1. We confirmed that the modified cysteine is Cys385 by LC-MS/MS and X-ray crystallography. Proteome-wide selectivity evaluation of UNC10013 (**4**) showed only Cys144 of NCK2, an adaptor protein unrelated to SETDB1, covalently modified by UNC10013 along SETDB1 Cys835, resulting in an off-target rate of 0.004% out of 23,149 detected cysteines. We achieved target family selectivity against a panel of methyltransferases and Kme readers in biophysical assays as well. Then, we identified UNC11277 (**11**) as a structurally related negative control, which contains the same covalent warhead as UNC10013 (**4**), for cellular assays. Finally, UNC10013 did not show toxicity for cellular treatment of MCF-7 for concentrations up to 50 µM and showed cellular target engagement of SETDB1 3TD at 30 µM in CeTSA, Unfortunately, UNC10013 has low cell permeability, as shown in Caco-2 evaluation and in CeTSA, which reduces its ability to potently engage its target in cells.

Five out of the six criteria for a covalent chemical probe were validated for UNC10013 (**4**) demonstrating that UNC10013 is an excellent starting point for the development of a SETDB1 3TD covalent chemical probe. Nevertheless, although we did not engage SETDB1 in cells at concentrations below 1 µM, UNC10013 remains an exciting chemical tool to study the 3TD of SETDB1. Indeed, we observed binding to the 3TD at 30 µM by CeTSA, suggesting that higher concentrations of compound will likely be required to compensate for the low cell permeability.

### UNC10013 is a negative allosteric modulator of the methyltransferase activity of SETDB1

Next, we examined whether UNC10013 would influence the allosteric relationship between the 3TD and the methyltransferase domains that we previously observed.^13^ While H3K9 methylation is not SETDB1-specific, the only known Akt methylation effector is SETDB1.^9,10^ Accordingly, we looked at the ability of UNC10013 (**4**) to modulate the SETDB1-mediated Akt methylation in HeLa cells after insulin stimulation. First, cells were pretreated for 24 h with different concentrations of UNC10013 (**4**) before the Akt methylation was triggered for 60 minutes using insulin (Fig. 4a). Results showed that treatment with 30 µM or more of UNC10013 (**4**) inhibited the Akt methylation and subsequent Akt Tyr308 phosphorylation. Further evaluation of UNC10013 (**4**) showed that the compound was able to inhibit the Akt methylation and subsequent phosphorylation after insulin activation up to 90 minutes, suggesting a robust inhibition (Fig. 4b).

**Figure 4.**
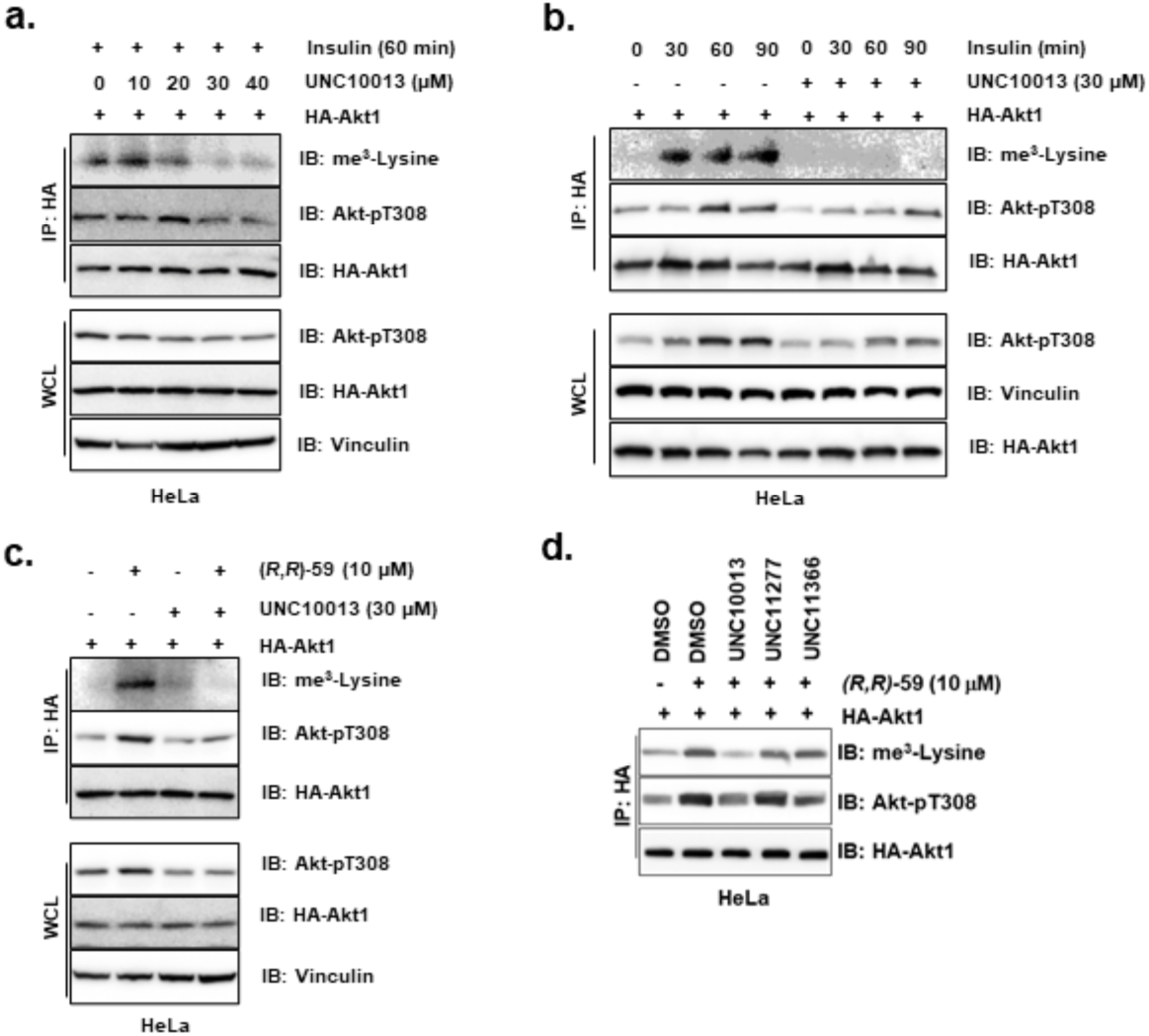
Cellular evaluation of UNC10013 (4) and UNC11277 (11) for the modulation of the SETDB1-mediated Akt methylation activity reveals negative allosteric modulation properties of UNC10013. (a) Pretreatment with different concentration of UNC10013 (**4**) for 24 h followed by 60 min of insulin stimulation of the Akt methylation. (b) Pretreatment with 30 µM of UNC10013 (**4**) for 24 h followed by different insulin stimulation treatment times. (c) Cotreatment with 10 µM of (R,R)-59 and 30 µM of UNC10013 (**4**) for 24 h. (d) Pretreatment with 30 µM of UNC10013 (**4**), UNC11277 (**11**) or UNC11366 (**12**) for 2 h followed by treatment with 10 µM of UNC8830 for 18 h.

Previous studies showed that the small-molecule 3TD ligand for SETDB1, (R,R)-59, is a positive allosteric modulator of the methyltransferase activity of SETDB1.^13^ Therefore, we decided to use (R,R)-59, instead of insulin, to activate the SETDB1-mediated Akt methylation. In this system, we observed that co-treatment with 10 µM of (R,R)-59 and 30 µM of UNC10013 (**4**) inhibited the (R,R)-59-triggered Akt methylation and phosphorylation (Fig. 4c). In another experiment, we found that pretreatment with 30 µM of UNC10013, but not the negative control UNC11277, for 2 h was able to prevent (R,R)-59-induced Akt methylation treatment in HeLa cells (Fig. 4d).

Knowing that (R,R)-59 and UNC10013 share the same binding pocket, these results demonstrate that UNC10013 (**4**) is a negative allosteric modulator of the methyltransferase domain of SETDB1, competing with (R,R)-59, and confirm that the 3TD is an allosteric site able to modulate the catalytic activity of the methyltransferase domain of SETDB1. This is significant because inhibitors of the catalytic SET domain have not yet been advanced as potential anti-cancer drugs and targeting the 3TD with negative allosteric modulators may provide this therapeutic opportunity. These experiments also suggest that UNC11277 is a good negative control for UNC10013 in cells.

## DISCUSSION

In these studies, we identified UNC6535 as the first small molecule ligand for the SETDB1 3TD able to occupy both the TD2 and TD3 aromatic cages simultaneously. UNC6535 showed a K_d_ of 4.2 µM for the 3TD and the ability to displace the H3K9Me2K14Ac peptide-mimicking the endogenous histone tail. Subsequent structure-based design identified UNC10013 (**4**) as a promising covalent chemical tool for the 3TD of SETDB1. X-ray crystallography confirmed that UNC10013 (**4**) maintains the unique binding pose of UNC6535 while covalently modifying Cys385. UNC10013 has a potent k_inact_/K_I_ of 1.04 x 10^6^ M^-1^s^-1^ and excellent proteome-wide selectivity. We also demonstrated that it could potently displace the H3K9Me2K14Ac-modified entire nucleosome from the 3TD. Caco-2 cell permeability, and CeTSA cellular target engagement evaluations revealed low cell permeability. However, UNC10013 (**4**) was able to engage the 3TD in HEK293T cells at 30 µM. Indeed, using SETDB1-mediated Akt methylation as a functional readout, we observed that UNC10013 (**4**) is a negative allosteric modulator of the methyltransferase activity of SETDB1 at concentrations of 30 µM and above revealing the ability to inhibit the methyltransferase activity of SETDB1 by targeting its 3TD.

In conclusion, we identified the first covalent ligand for the 3TD of SETDB1 and demonstrated that it can allosterically inhibit the methyltransferase activity of SETDB1 in cells. This chemical tool complements the previously identified (R,R)-59 compound which is a positive allosteric modulator of SETDB1 catalytic activity, and also binds the 3TD. Indeed, this pair of compounds should allow researchers to gain a better understanding of the function of the 3TD of SETDB1 and its relationship with the other domains, especially the methyltransferase domain.

Allosteric modulators are especially attractive since the SET domain of methyltransferases is highly conserved within the family, therefore drugging the allosteric Kme reader domain is a promising strategy to potentially achieve selectivity for SETDB1 SET domain inhibition. Moreover, decrease in Akt methylation and subsequent phosphorylation achieved by SETDB1 depletion has been linked to decreased tumor growth *in vivo*.^10^ Therefore, the development of an allosteric negative modulator ligand for the 3TD of SETDB1 is a promising approach to develop cancer or neurological disorder-targeting therapies, especially as there are currently no structural data or potent confirmed molecules directly targeting the methyltransferase SET domain.^19,20^

In the future, we intend to address the cell permeability issues of UNC10013 (**4**) by structural modifications and cellular evaluation to provide a more potent 3TD covalent chemical probe for cellular studies.

## METHODS

### Protein and Peptide Production and Purification

#### Expression constructs

The chromo domains of CBX2 (residues 9-66 of NP_005180) and MPP8 (residues 55-116 of NP_059990) were expressed with N-terminal His-tags in modified pET28 expression vectors. The chromo domain of CDYL2 (residues 1-75 of NP_689555) was expressed with a C-terminal His-tag in a pET30 expression vector. The tudor domain of SETDB1 (residues 196-403 of NP_036564) was expressed with an N-terminal His-tag in a modified pET28 expression vector. The SETDB1 C385A mutant was generated using a QuickChange II site-directed mutagenesis kit (Agilent) according to manufacturer’s recommendations. The tudor domain of 53BP1 (residues 1484-1603 of NP_005648) was expressed with an N-terminal His tag in a pET15 expression vector. The tudor and PHD domains of PHF1 (residues 27-360 of NP_077084) and PHF19 (residues 1-207 of NP_001009936) were expressed with N-terminal His-tags in modified pET28 expression vectors. The Jumonji domain of KDM7B (residues 1-447 of NP_055922) was expressed with an N-terminal MBP-tag and a C-terminal His-tag. Full length SETDB1 (residues 1-1291 of NP_001138887) was expressed in a pFastBac baculovirus expression vector with an N-terminal His-tag and a C-terminal FLAG tag. Truncated SETDB1 (residues 570-1291 of NP_001138887) was expressed with a N-terminal GST and His-tag in a pFastBac baculovirus expression vector.

#### Protein expression and purification

All E. coli expression constructs were transformed into Rosetta BL21(DE3)pLysS competent cells (Novagen, MilliporeSigma). Protein expression was induced by growing cells at 37°C with shaking until the OD600 reached ∼0.6-0.8 at which time the temperature was lowered to 18°C and expression was induced by adding 0.5mM IPTG and continuing shaking overnight. Cells were harvested by centrifugation and pellets were stored at −80°C.

All His-tagged proteins except for KDM7B were purified by resuspending thawed cell pellets in 30ml of lysis buffer (50mM sodium phosphate pH 7.2, 50mM NaCl, 30mM imidazole, 1X EDTA free protease inhibitor cocktail (Roche Diagnostics) per liter of culture. Cells were lysed on ice by sonication with a Branson Digital 450 Sonifier (Branson Ultrasonics) at 40% amplitude for 12 cycles with each cycle consisting of a 20 second pulse followed by a 40 second rest. The cell lysate was clarified by centrifugation and loaded onto a HisTrap FF column (Cytiva) that had been preequilibrated with 10 column volumes of binding buffer (50mM sodium phosphate pH 7.2, 500mM NaCl, 30mM imidazole) using an AKTA FPLC (Cytiva). The column was washed with 15 column volumes of binding buffer and protein was eluted in a linear gradient to 100% elution buffer (50mM sodium phosphate pH 7.2, 500mM NaCl, 500mM imidazole) over 20 column volumes. Peak fractions containing the desired protein were pooled and concentrated to 2ml in Amicon Ultra-15 concentrators 3,000 molecular weight cut-off (Merck Millipore). Concentrated protein was loaded onto a HiLoad 26/60 Superdex 75 prep grade column (Cytiva) that had been preequilibrated with 1.2 column volumes of sizing buffer (25mM Tris pH 7.5, 250mM NaCl, 0.5mM TCEP, 5% glycerol for MPP8 or 25mM Tris pH 7.5, 250mM NaCl, 2mM DTT, 5% glycerol for all other proteins) using an ATKA Purifier (Cytiva). Protein was eluted isocratically in sizing buffer over 1.3 column volumes at a flow rate of 2ml/min collecting 3ml fractions. Peak fractions were analyzed for purity by SDS-PAGE and those containing pure protein were pooled and concentrated using Amicon Ultra-15 concentrators 3,000 molecular weight cut-off (Merck Millipore).

#### KDM7B purification

KDM7B was purified as described above for His-tagged proteins with the following exceptions. Peak fractions from the HisTrap column were pooled and concentrated in Amicon Ultra-15 concentrators 50,000 molecular weight cut-off (Merck Millipore). The N-terminal MBP tag was removed by TEV cleavage as described above for MPP8. The cleavage reaction was then passed over a HisTrap FF column (Cytiva) to remove any protein that still retained the tag as well as the His-tagged TEV protease. The column flow through was collected and concentrated to 2ml using Amico Ultra-15 concentrators, 30,000 molecular weight cut-off (Merck Millipore). Concentrated protein was loaded onto a HiLoad 26/60 Superdex 200 prep grade column (Cytiva) that had been preequilibrated with 1.2 column volumes of sizing buffer (25mM Tris pH 7.5, 250mM NaCl, 2mM DTT, 5% glycerol) using an ATKA FPLC (Cytiva). Protein was eluted isocratically in sizing buffer over 1.3 column volumes at a flow rate of 2ml/min collecting 3ml fractions. Peak fractions were analyzed for purity by SDS-PAGE and those containing pure protein were pooled and concentrated using Amicon Ultra-15 concentrators 30,000 molecular weight cut-off (Merck Millipore).

#### Baculovirus expression and purification

Baculoviruses for full length and truncated SETDB1 were generated using the Bac-to-Bac expression system (Invitrogen). Sf9 cells were grown in Sf900III SFM media (Gibco) supplemented with 2%FBS to 1x106 cells/ml and infected at an MOI of 5 with a baculovirus containing either full length SETDB1 or truncated SETDB1. Cells were harvested after 72hr, and cell pellets were stored at −80⁰C until purification.

#### Truncated SETDB1 purification

Cell pellets from 1L of infected cells were thawed and resuspended in 30ml of cytobuster (Merck Millipore) containing 1X EDTA free protease inhibitor cocktail (Pierce), 1X phosphatase inhibitor cocktail (Pierce) and 30mM imidazole, pH 8.0. Cells were lysed for 15 minutes at room temperature with gentle mixing. Cell lysates were clarified by centrifugation and the clarified supernatant was loaded onto a 1ml HisTrapFF column (Cytiva) that had been preequilibrated with 10 column volumes of binding/wash buffer (50mM sodium phosphate, pH 7.2, 500mM NaCl, 30mM imidazole) at a flow rate of 1ml/min using an AKTA FPLC (Cytiva). The column was washed with 15 column volumes of binding/wash buffer and bound protein was eluted in a linear gradient over 20 column volumes to 100% elution buffer (50mM sodium phosphate buffer, pH 7.2, 500mM NaCl, 500mM imidazole). Peak fractions were run on SDS-PAGE gels and analyzed by Coomassie staining. Truncated SETDB1 containing fractions were pooled and concentrated in Amicon Ultra 30K cut off concentrators (Merck Millipore) and buffer was exchanged into 25mM HEPES, pH 7.5, 300mM NaCl, 0.5mM TCEP, 10% glycerol, 0.04% Triton. Protein concentration was determined by Bradford assay. Protein was aliquoted and stored at −80⁰C.

#### Full-length SETDB1 purification

Cell pellets from 1L of infected cells were thawed and resuspended in 30ml of cytobuster (Merck Millipore) containing 1X EDTA free protease inhibitor cocktail (Pierce), and 1X phosphatase inhibitor cocktail (Pierce). Cells were lysed for 15 minutes at room temperature with gentle mixing. Cell lysates were clarified by centrifugation and the clarified supernatant was incubated with ANTI-FLAG M2 affinity gel (Sigma) overnight at 4°. The FLAG gel was washed with TBS and protein was eluted by competition with FLAG peptide (Sigma). Elution fractions were run on SDS-PAGE gels and analyzed by Coomassie staining. Full length SETDB1 containing fractions were pooled and concentrated in Amicon Ultra 50K cut off concentrators (Merck Millipore) and buffer was exchanged into 25mM HEPES, pH 7.5, 300mM NaCl, 0.5mM TCEP, 10% glycerol, 0.04% Triton. Protein concentration was determined by Bradford assay. Protein was aliquoted and stored at −80⁰C.

#### Peptide production

H3K9Me2K14Ac-Biotin (aa 1-19) peptide was purchased from Genscript. The countersalts are hydrochloric salts. The sequence is as followed:

H3K9Me2K14Ac-Biotin (aa 1-19): MARTKQTAR{Lys(me3)}STGG{Lys-Ac}APRKQ{K(Biotin)}.

### Nucleosome Production and Purification

**Scheme 1.**
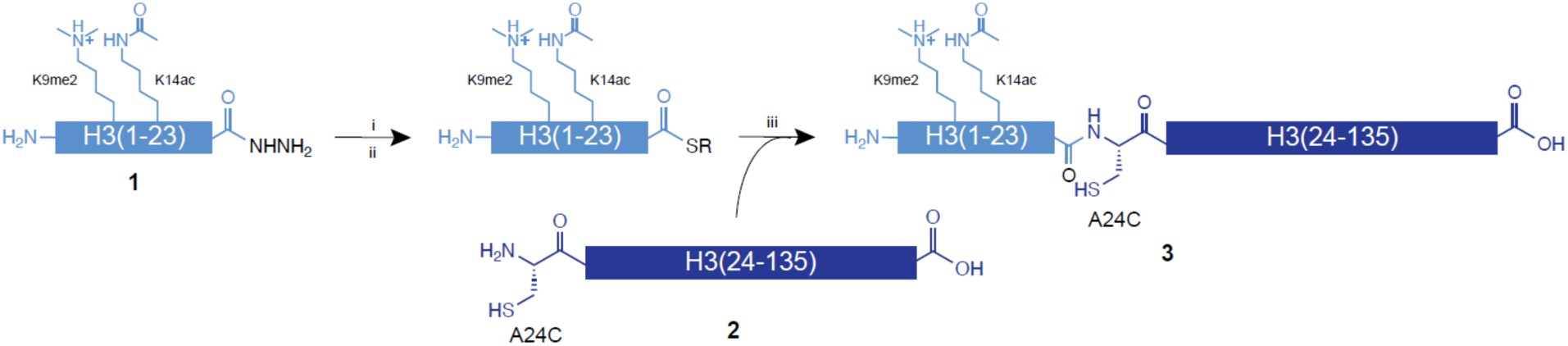
Semisynthesis of H3K9Me2K14Ac(A24C, C110A). H3K9Me2K14Ac(A24C, C110A), **3**, was generated via expressed protein ligation using peptide **1** and protein **2**. i) 0.2 M phosphate, 6 M Guanidine·HCl (GnCl), NaNO2 (10 eq), pH 3.0-3.1, −15 °C,15-20 min. ii) 4-mercaptophenylacetic acid (MPAA, 100 eq.), pH 6.8-7.0, room temperature, R = phenylacetic acid. iii) Expressed protein ligation, 0.2 M phosphate, 6 M GnCl, MPAA, room temperature, 24 h.

#### Preparation of histone peptide and protein fragments for expressed protein ligation

Peptide **1**, containing H3.2(1-23, K9Me2, K14Ac, C-terminal hydrazide) was synthesized by the UNC High-throughput Peptide Synthesis and Array Facility, purified by reverse phase HPLC (RP-HPLC), and confirmed by MALDI spectrometry. The gene sequence encoding human H3.2 residue 24-135 was cloned into the pST50Tr vector with an N-terminal TEV protease cleavage sequence (ENLYFQ) and A24C and C110A mutations. ENLYFQ-H3.2 (24-135, A24C, C110A, hereafter, H3.2-C) was expressed in E. coli BL21(DE3)pLysS cells at 37 °C for 3 hours. The ENLYFQ-H3.2-C protein was extracted from inclusion bodies as previously described^21^ and dialyzed overnight into 0.1% TFA in water at 4 °C. The dialyzed protein was cleared by centrifugation and filtration (0.45 µm) and purified by RP-HPLC (Vydac 218TP C18 column, 250 x 22 mm, 10 µm) using a 40– 70% B gradient over 60 min (A: 0.1% TFA in water; B: 90% acetonitrile, 0.1% TFA in water) and lyophilized. The identity of the ENLYFQ-H3.2-C protein was verified by LC/MS. The purified H3.2-C protein was dissolved in 50 mM Tris-HCl, pH 8.0, 1 mM DTT, 0.5 mM EDTA, 6 M guanidine HCl (GnCl) and then diluted in the same buffer lacking GnCl to a final concentration of 1 M GnCl and 1 mg/ml protein. The ENLYFQ sequence was cleaved with the addition of 1/100 molar eq TEV protease for 2 hours at room temperature and purified by RP-HPLC exactly as described above, yielding H3.2-C protein **2**.

#### Expressed protein ligation

The expressed protein ligation reaction between peptide **1** and H3.2-C protein **2** was performed essentially as previously described.^22^ Briefly, peptide **1** (0.9 mg, 0.35 µmol) was dissolved in 70 µl of 0.2 M phosphate containing 6 M GnCl (pH 3.0–3.1). The peptide hydrazide was converted to an azide by addition of 10 molar equivalents of NaNO_2_ using a 0.5 M NaNO_2_ stock. This reaction was allowed to proceed for 15–20 min at −15 °C in an ice/salt bath. The H3.2-C protein **2** (4.1 mg, 0.32 µmol) was dissolved in 70 µl of 0.2 M phosphate containing 6 M GnCl (pH 6.8-7.0) containing 100 molar eq of 4-mercaptophenylacetic acid (MPAA, Sigma) and added to the peptide. The pH of the ligation mixture was adjusted to 6.8-7.0 and the ligation was allowed to proceed for 24 h at room temperature, affording H3.2K9me2K14ac(A24C, C110A), **3**. Ligation product **3** was purified by RP-HPLC (Vydac 218TP C18 column, 250 x 10 mm, 5 µm) with a 43– 53% B gradient over 45 min, yielding 2.3 mg. The identity of the ligation product was verified by LC/MS.

#### Preparation of nucleosomes

The human H3.2(A24C, C110A) mutant was cloned in the pST50Tr vector by site-directed mutagenesis. Human histones H2A, H2B, H3.2(A24C, C110A), and H4 were expressed and purified from inclusion bodies as previously described.^21^ H2A(K119C)/H2B dimer, H3.2K9me2K14ac(A24C, C110A)/H4 tetramer, and unmodified, control H3.2(A24C, C110A)/H4 tetramer were reconstituted and purified by cation exchange chromatography as previously described.^23^ The H2A(K119C)/H2B dimer was labeled with biotin-maleimide exactly as previously described.^16^ Modified H2A(K119C-biotin)/H2B/H3.2K9me2K14ac(A24C, C110A)/H4 and unmodified H2A(K119C-biotin)/H2B/H3.2(A24C, C110)/H4 nucleosomes were assembled with 185 bp 601 nucleosome positioning sequence^24^ with 20 bp symmetric linker DNA and purified as previously described.^21,23^

### TR-FRET Displacement Assay

#### H3K9Me2K14Ac Peptide Displacement from SETDB1 3TD

The TR-FRET assay protocol was developed and adapted from a previously reported protocol.^18^ Briefly, the assay was run using white, low-volume, flat-bottom, nonbinding, 384-well microplates (Greiner, 784904) containing a total volume of 10 μL per well. The buffer was made of 1X PBS pH 7.0, 0.005% Tween 20, and 2 mM DTT. Lance Europium (Eu)-W1024 Streptavidin conjugate (2nM) and LANCE Ultra ULightTM-anti-6x-His antibody (10 nM) were used as donor and acceptor fluorophores associated with the tracer ligand and protein respectively. Final assay concentrations of 40 nM 6X histidine-tagged SETDB1 (residues 195-403, N-terminal tag) and 40 nM of H3K9Me2K14Ac-Biotin (aa 1-19) were used for compound testing. A 10-point, three-fold serial dilution of each compound was tested as the primary hit validation. Assay components were added to an assay plate using a Multidrop Combi (ThermoFisher). After addition, the plates were sealed with clear covers, mixed gently on a shaker for 1 min, centrifuged at 1000x g for 2 minutes, and allowed to equilibrate for 1 hour in the dark. Plates were read with an EnVision 2103 Multilabel Plate Reader (PerkinElmer) using an excitation filter at 320 nm and emission filters at 615 and 655 nm. Emission signals (615 and 665 nm) were measured simultaneously using a dual mirror D400/D630 (using a 100-microsecond delay). TR-FRET output sign was expressed as emission ratios of acceptor/donor (665/615 nm) counts. Percent inhibition was calculated on a scale of 0% (i.e. activity with DMSO vehicle only) to 100% (100 μM H3K9Me2K14Ac) using two full columns of control wells on each plate. The data was fitted with a four-parameter nonlinear regression analysis using ScreenAble to determine IC50 values, plotted on GraphPad for visualization, and reported as an average of at least three technical replicates ± s.e.m.

#### Kme Reader Selectivity Evaluation

Similar protocols to those described above were used for the TR-FRET evaluation of the Kme reader selectivity of compounds using the parameters below:

| Protein | Tracer Ligand | Donor Fluorophore | Acceptor Fluorophore | Buffer |
| --- | --- | --- | --- | --- |
| **CBX2** (30 nM) | UNC4195 (30 nM) | 10 nM Europium-W1024 | 2 nM and LANCE Ultra ULight-anti6x-His | 20 mM Tris-HCl, 150 mM NaCl, 0.05% Tween-20 (v/v), and 2 mM DTT |
| **CDYL2** (10 nM) | UNC4848 (10 nM) | 10 nM Europium-W1024 |  |  |
| **MPP8** (5 nM) | P42 (20 nM) | 10 nM Europium-W1024 |  |  |
| **PHF1** (60 nM) | UNC6957 (90 nM) | 10 nM Europium-W1024 |  |  |
| **PHF19** (20 nM) | UNC6957 (40 nM) | 10 nM Europium-W1024 |  |  |
| **KDM7B** (30 nM) | P3883 (50 nM) | 10 nM Europium-W8044 |  |  |
| **CHD1** (30 nM) | H3K4Me2-biotin (60 nM) | 10 nM Europium-W1024 |  | 20 mM Tris-HCl, 75 mM NaCl, 0.05% Tween-20 (v/v), and 2 mM DTT |

#### Nucleosome TR-FRET Displacement

The assay was run using white, low-volume, flat-bottom, nonbinding, 384-well microplates (Greiner, 784904) containing a total volume of 10 μL per well. The buffer was made of 1X PBS pH 7.0, 0.005% Tween 20, and 2 mM DTT. Lance Europium (Eu)-W1024 Streptavidin conjugate (2nM) and LANCE Ultra ULightTM-anti-6x-His antibody (50 nM) were used as donor and acceptor fluorophores associated with the tracer ligand and protein respectively. Final assay concentrations of 15 nM 6X histidine-tagged SETDB1 (residues 195-403, N-terminal tag) and 15 nM of H3K9Me2K14Ac-modified nucleosome were used for compound testing. A 10-point, three-fold serial dilution of each compound was tested as the primary hit validation. After addition, the plates were sealed with covers, mixed gently on a shaker for 1 min, centrifuged at 1000x g for 2 minutes, and allowed to equilibrate for 1 hour in the dark. Plates were read with an EnVision 2103 Multilabel Plate Reader (PerkinElmer) using an excitation filter at 320 nm and emission filters at 615 and 655 nm. Emission signals (615 and 665 nm) were measured simultaneously using a dual mirror D400/D630 (using a 100-microsecond delay). TR-FRET output sign was expressed as emission ratios of acceptor/donor (665/615 nm) counts. Percent inhibition was calculated on a scale of 0% (i.e. activity with DMSO vehicle only) to 100%. The data was fitted with a four-parameter nonlinear regression analysis using GraphPad, where IC_50_ values were extracted after constraining the bottom to 0% and the top to 100%, and reported as an average of at least three technical replicates ± s.e.m.

### Isothermal Titration Calorimetry

ITC experiments were carried out at 10 °C on an Auto-ITC calorimeter (Malvern). Protein and peptide were kept in an identical buffer of 20 mM Tris/HCl [pH 7.0], 150 mM NaCl, and 2 mM TCEP. The final DMSO concentration was kept consistent throughout the cell and syringe. ITC experiments were carried out with protein and ligand concentrations of 100 μM and 1.5 mM, respectively. Binding isotherms were analyzed by nonlinear least-squares fitting of the data using Microcal ORIGIN software (Microcal) using a one-site binding model and reported as an average of two technical replicates ± s.e.m.

### Radiometric Methyltransferase Activity Evaluation

#### Dose-response Evaluation

IC_50_ values were determined for the compounds under the following conditions: For SETDB1, the assay used 2 nM SETDB1, 5 µM SAM, and 100 nM biotinylated H3 1-25, with a 15-minute incubation at 23 °C. For SETD2, the assay included 150 nM SETD2, 5 µM SAM, and 1 µM biotinylated H3 (21-44), with 20 µL of reaction mixtures incubated for 1 hour at 23 °C in buffer (20 mM Tris, pH 8.0; 0.01% Triton X-100; 5 mM DTT). The PRMT1 assay was performed as previously described.^25^

#### Methyltransferase Selectivity Evaluation

The inhibitory effects of two compounds, UNC10013 and UNC10016, on the enzymatic activities of 33 epigenetic targets were examined using radioactivity-based activity assays as previously described.^25^

### MALDI-TOF-based Methyltransferase Activity Assay

A catalytic assay was developed based on a previously described protocol by Basavapathruni et al.^26^ The selected buffer was 20 mM bicine [pH 7.6], 0.002% Tween 20, and 1 mM DTT with the DTT being added fresh before use. The final conditions of reagents were as follows: 5 nM full-length SETDB1 (ActiveMotif), 1 μM biotinylated H3 unmodified peptide substrate (ARTKQTARKSTGGKAPRKQL-K(Biot)-NH2, GenScript), and 20 μM SAM. 384-well assay-ready plates were prepared with 10-point dose-response curves. For compound testing post-EpiG set screening, column 1 contained complete enzymatic inhibition by UNC00000421A at 100 μM. Column 2 contained a 16-point, three-fold dilution dose-response curve of UNC00000421A starting at 200 μM and going down the column. Columns 3-12 and 13-22 were used for compound dose-response curves. Column 23 was reserved for 2% DMSO only and Column 24 was a no enzyme control. Assay ready plates were centrifuged for 5 minutes at 1500xg. After centrifugation, 9 μL of a mixture of SAM and SETDB1 was added using Multidrop Combi (ThermoFisher) to columns 1-23. 9 μL of solution of SAM was added using a multi-channel pipette. The plate was incubated plates for 15 minutes in a dark cabinet before adding 1 μL of the substrate to all columns using the MultiDrop Combi. The reactions were allowed to proceed for 1 hour in a dark cabinet (not stacked). The reactions were quenched at 1 hour with 1 μL of 5% formic acid (0.5% final) using the MultiDrop Combi, mixed gently on a tabletop mixer for 1 minute, and then centrifuged for 2 minutes at 1500xg. Plates were shipped for analysis by SAMDI Tech, Inc.

### Fluorescence Polarization

Fluorescence polarization (FP) experiments were conducted in black low-volume 384-well plates (Greiner Bio-One, US) with a final volume of 20 µL per well. Each compound was serially diluted (12 or 16 concentrations) at 1.1X the tested concentrations in 18 µL of an optimized buffer containing 20 mM Tris-HCl pH 8, 0.01% (v/v) Triton X-100, and 275 nM SETDB1 TTD (final concentration 250 nM in 20 µL). The mixtures were preincubated at room temperature (∼25 °C) for 1 hour, maintaining a final DMSO concentration of 2% or 4% in the assay. Following preincubation, 2 µL of a 200 nM stock of 5’ FITC-labeled H3K9Me2K14Ac peptide in the optimized assay buffer was added to each well using a Multidrop Combi dispenser (Thermo Scientific), resulting in a final fluorescent peptide concentration of 20 nM. After a brief centrifugation at 300 g (Eppendorf Centrifuge 5810R) for 5 minutes, the assay plates were covered to protect from light and incubated for 15 minutes at room temperature. The fluorescence polarization (FP) ratio was then measured using a BioTek Synergy H1 microplate reader (BioTek, Winooski, VT) with excitation at 485 nm and emission at 528 nm. The polarization values obtained were converted to percentage activity relative to the control with 100% activity and processed in GraphPad Prism using Sigmoidal, 4PL, X is log(concentration) fit with the bottom and top constrained to 0% and 100%, respectively. Each experiment was performed in triplicate, and the results were presented as the average and standard deviation of the three measurements.

### Crystallography

#### Protein expression and purification

PCR amplified cDNA encoding SETDB1 TTD residues 197-403 was subcloned into the pET28-MHL vectors, downstream of the poly-histidine coding region. Following the transfection into *E. coli* BL21-CodonPlus-RIL, cultured cells were grown overnight at 37 ^0^C in Terrific Broth supplemented with 50 µg/mL Kanamycin and 35 µg/ml chloramphenicol. SETDB1 TTD expression was induced by adding 1 mM IPTG (isopropyl-1-thio-D-galactopyranoside) and allowing overnight expression at 18 ^0^C. The obtained culture was pelleted at low speed (7000 rpm for 10 minutes at 4 ^0^C in a Beckman Coulter centrifuge). Harvested cells were resuspended (1 g of cell pellet per 10 mL) in the extraction buffer (20 mM HEPES pH 7.5, 500 mM NaCl, 2 mM TCEP, 5% glycerol) with protease inhibitor (0.1 mM phenylmethyl sulfonyl fluoride, PMSF). The cell suspension was supplemented with 5 µl/L Benzonase Nuclease (in-house prepared) and sonicated for mechanical lysis on ice for 10 min total (10s pulses with 7s interruptions) (Sonicator 3000, Misoni). The crude extract was then clarified by high-speed centrifugation (60 min at 20,000 ×g at 4^0^C) in a Beckman Coulter centrifuge to remove the cellular debris. Clarified lysate was first sent through the HispurTM Ni-NTA resin (Thermo Scientific) column (∼ 20 ml bed volume for 60-70 g of the pellet) and eluted with 250 mM imidazole. Eluted fractions were passaged through Gel filtration HiLoadTM 26/600 Superdex (Cytiva) with 20 mM HEPES, pH 7.5, 250 mM NaCl, 0.5 mM TCEP, and 5% glycerol and further enriched with ion exchange Mono S HR 16/10 (SOURCE^TM^ 30Q GE Healthcare) to 95% purity. Following the identification of SETB1 TTD fractions on SDS-PAGE gels and confirmation of identity with LC-MS enriched fractions were pooled, concentrated to 7-8 mg/ml, snap-frozen, and stored at −80 ^0^C till use.

#### Crystallization and structural determination

SETDB1-UNC6535 structure was obtained by first generating Apo SETDB1-TTD crystals using 96-well vapor diffusion sitting drop plates by mixing equimolar amounts of protein and reservoir solution containing 20% (w/v) PEG8000, 0.2M Sodium Chloride, 0.1M Hepes, pH 7.5, 5% MPD. Apo crystals were then soaked with 5 mM UNC6535 ligand at room temperature for 2 hours, then mounted by cryo-cooling in liquid nitrogen.

SETDB1-UNC10013 and SETDB1-UNC10016 structures were obtained by co-crystallization method. SETDB1-TTD protein, at 7 mg/mL (0.266 mM) was pre-incubated for 30 minutes at room temperature with 2.6 mM (10X times molar excess) of UNC10013 and UNC10016 compounds in the presence of 1:1000 ratio (trypsin: SETDB1 TTD) before crystallization. Co-crystallization of SETDB1-UNC10013 and SETDB1-UNC10016 were carried out in 96-well vapor diffusion sitting-drop plates by mixing 0.5 μL of protein-compound mixture and 0.5 μL of crystallization screen using Phoenix liquid dispensing robot (Art Robbins). Initial crystals were obtained after 72 hours at 18 ^0^C. After further optimization of the protein:compound ratio (2X times molar excess) and seeding, well-diffracting co-crystals of UNC10013 and UNC10016 were obtained in crystallization conditions containing 20% PEG 8000, 0.2M NaCl, 0.1M HEPES pH 7.5, 5% Ethyl Glycol, and 25% P3350, 0.1M NH4SO4, 0.1M Bis-Tris pH 6.5, respectively. Crystals were then cryocooled in liquid nitrogen. Dataset for SETDB1-UNC6535 was collected in-house using FRE Rigacu superbright. Datasets for SETDB1-UNC10013 and SETDB1-UNC10016 were collected at the CMCF-ID beamline at the Canadian Light Source (CLS). Datasets were processed with HKL3000. Initial phases for SETDB1-UNC6535 was obtained by using (PDB ID:5KE2) as initial model in Fourier transform with Refmac. Initial phases for SETDB1-UNC10013 and SETDB1-UNC10016 were obtained by using (PDB ID: 3DLM) as initial model in PHASER for molecular replacement. Model building was performed in COOT and the structure was validated with Molprobity. Ligand restraints were generated using Grade Web Server (http://grade.globalphasing.org).

### MALDI-TOF

#### Initial Assessment of Covalent Bond Formation

The buffer was made of 25 mM Tris/HCl, 50 mM NaCl (pH 7.5). Experiments were carried out using 98 µL of SETDB1-3TD (residues 195-403, N-terminal 6X His tag) protein and 2 µL of compound for final concentrations of 10 µM of protein and 100 µM of compound with a 2% final DMSO concentration. After 2 hours of incubation at room temperature, 1 µL of the mixture and 1 µL of α-cyano-4-hydroxycinnamic acid (CHCA) MALDI matrix were spotted on a MALDI sample plate. Each spot was analyzed by MALDI-TOF on an AB Sciex 5800 MALDI-TOF/TOF instrument using the linear mode. Data is visualized using mMass version 5.5.0 with normalized intensities, baseline corrections (precision: 15, relative offset: 25) and smoothing (window size: 30 m/z, 5 cycles).

#### Qualitative Reactivity Comparison

A similar protocol than described above was used except that the final concentrations of protein and compound were set at 10 µM and the mixture was incubated on ice. At the desired time point, an 8 µL sample was quenched by adding 2 µL of 5% TFA aqueous solution (v/v) before being spotted.

### Intrinsic Reactivity Evaluation with Glutathione

The protocol was adapted from Resnick et al.^27^ Briefly, 500 µM of compound was incubated with 2.5 mM of glutathione and 800 µM of Rhodamine B, used an internal standard, in 100 mM potassium phosphate buffer (pH 7.4) and MeCN (9:1 ratio respectively). Every hour, for up to 8 h, 6 µL were analyzed on the LC-MS instrument. LC-MS data for all compounds was acquired using an Agilent 6110 Series system with the UV detector set to 254 nm. Samples were injected (<10 μL) onto an Agilent Eclipse Plus 4.6 × 50 mm, 1.8 μm, C18 column at room temperature. Mobile phases A (H2O + 0.1% acetic acid) and B (MeCN + 1% H2O + 0.1% acetic acid) were used with a linear gradient from 10% to 100% B in 5.0 min, followed by a flush at 100% B for another 2 minutes with a flow rate of 1.0 mL/min. Mass spectra (MS) data were acquired in positive ion mode using an Agilent 6110 single quadrupole mass spectrometer with an electrospray ionization (ESI) source. The % unreacted compound was calculated by normalizing the area under curve ratio of the compound to Rhodamine B. The resulting % unreacted compound was plotted in function of time and the data was fitted with a one-phase decay regression analysis using GraphPad providing the t_1/2_ values.

### Intact Protein Mass Spectrometry

40 µM of 3TD protein was incubated with 400 µM of compound in 25 mM HEPES pH 7.5, 300 mM NaCl for a total volume of 20 µL. The sample was incubated overnight at 4 °C before undergoing sample preparation for mass spectrometry.

Samples were diluted to a concentration of 0.5 µg/µL in peptide background electrolyte (BGE). Samples were injected onto an HR Chip (908Devices Inc.) using a ZipChip system (908Devices Inc.) interfaced to a Thermo QExactive HF Biopharma mass spectrometer. The MS data was acquired through Tune (Thermo, v. 2.9) and settings included: scan range 500-2000 m/z, in-source CID 15eV, mass resolution 15,000, 3 microscans, 10 ms maximum injection time. The ZipChip settings included: field strength 500 V/cm, injection volume 1 nL, pressure assist start time 0.5 min, analysis run time 3.5 min. Intact mass spectra were deconvoluted using PMI Intact Mass software (Protein Metrics Inc. v5.1.1). Provided sequences were used to assign peaks in Byos. Threshold for protein assignment = +/- 5 Da. Relative abundance was calculated by dividing the intensity of a given peak by the max peak intensity.

### Liquid Chromatography coupled to Tandem Mass Spectrometry (LC-MS/MS)

The same samples used aboved were diluted to 50 μl with 2M urea. Samples were reduced with 5 mM DTT at 37°C for 45 min, then alkylated with 15 mM iodoacetamide at room temperature in the dark for 45 min. The samples were diluted to 1M urea and subjected to digestion with trypsin (Promega) overnight at 37°C at a 1:20 enzyme: protein ratio. The resulting peptide samples were acidified to 0.5% TFA, desalted using C18 Desalting Spin Columns (Thermo), then the eluates were dried via vacuum centrifugation. The samples were reconstituted in a 2% ACN, 0.1% formic acid solution prior to LC-MS/MS analysis.

Samples were analyzed in technical duplicate by LC-MS/MS using a Thermo Easy nLC 1200 – QExactive HF Mass Spectrometer. Samples were injected onto an Easy Spray PepMap C18 column (75 μm id × 25 cm, 2 μm particle size) (Thermo Scientific) and separated over a 45 min method. The gradient for separation consisted of 5-38% mobile phase B at a 250 nl/min flow rate, where mobile phase A was 0.1% formic acid in water and mobile phase B consisted of 0.1% formic acid in 80% ACN. The QExactive HF was operated in data-dependent acquisition mode (DDA) where the 15 most intense precursors were selected for subsequent fragmentation. Resolution for the precursor scan (m/z 350-1600) was set to 60,000 with a target value of 3 × 106 ions, 100 ms max injection time. MS/MS scans resolution was set to 15,000 with a target value of 5 × 104 ions, 60 ms max injection time. The normalized collision energy was set to 27% for HCD. Dynamic exclusion was set to 30 s, peptide match was set to preferred, and precursors with unknown charge or a charge state of 1 and ≥ 7 were excluded.

Raw data were analyzed using Proteome Discoverer 2.5 (Thermo Scientific). The samples were searched against the Recombinant 3TD protein sequence, along with an E. coli database from Uniprot (UP000000625; containing 4,448 protein sequences). A common contaminants database was appended to the reference proteome. Parameters used in PD included: precursor ion tolerance – 20ppm; product ion tolerance – 20ppm; enzyme specificity, set to trypsin, up to two missed cleavages were allowed. Variable modifications: methionine oxidation cysteine carbamidomethylation, compound covalent modification.

### k_inact_/K_I_ Measurement using mass dilution SETDB1 3TD TR-FRET

Mass dilution TR-FRET assays were run using the white, high volume, flat-bottom, nonbinding, 384-well OptiPlateTM from PerkinElmer (Part No. 6007290) containing a total assay volume of 100 μL per well. All experiments were run at room temperature with the same buffer composition and final concentrations of protein and FRET reagents as the TR-FRET methods described above. The test compounds and SETDB1 3TD protein were diluted in Eppendorf tubes for incubation at the desired testing concentrations (2.5X). In the OptiPlate, the remaining components necessary for the FRET assay were dispensed at 60 μL (1.67X) per well in preparation for mass dilution. Starting at time point zero, 40 μL aliquots of the protein-peptide stock were transferred from the Eppendorf tube to the Optiplate for mass dilution every three minutes for 30 minutes total (11 total wells per concentration, 100 μL total). The dilution factors were predetermined so the mass dilution step reduces the peptide concentration below the reported IC50. All TR-FRET components are diluted to the final concentrations listed in the TR-FRET methods above (1X). A DMSO control sample was also included for data normalization at each time point tested. 100% (saturated with known inhibitor) and 0% (no inhibitor present) inhibition controls were incorporated to calculate the percentage of inhibition for all data points. Following mass dilution, each time point was incubated for 15 minutes to allow for any reversible compound binding to be diluted off. Then, each data point was read on an EnVision 2103 Multilabel Plate Reader (PerkinElmer) using a 320 nm excitation filter and 615 and 665 nm emission filters. Emission signals were measured simultaneously using a dual mirror D400/D630 and 100 μsec delay. The TR-FRET output signal was expressed as a ratio of acceptor/donor (665 nm/615 nm) emission counts. The normalized percent inhibition for each concentration tested was plotted at each time point. The data was fitted using a linear fit, and the slope (kobs) of each concentration was plotted against the different peptide concentrations tested. From this, the max k_obs_ from these experiments is taken as the k_inact_, while the concentration at which ½kinact is observed is taken as the K_I_. From here, the inactivation efficiency can be calculated using k_inact_/K_I_. GraphPad Prism 9.0 was used for data visualization, fitting, and figure preparation.

### Differential Scanning Fluorimetry

DSF assays were performed using AB ViiA 7 Real-Time PCR System. The buffer was made of 20 mM HEPES, 200 mM NaCl, pH 7.5. Experiments were carried out using 8 μL of SETDB1-3TD (residues 196-403, N-terminal His tag) and 20X concentration Sypro Orange dye (5000X DMSO stock, ThermoFisher) and 2 uL of ligand for a final concentration of 20 μM of protein and 200 μM of ligand with 1.1% of DMSO. Plates were incubated for 15 min before running the temperature gradient (1 °C/min, from 25 to 90 °C). Values are presented as an average of at least two replicates ± s.d. calculated using the Boltzmann method from the Protein Thermal Shift software (ThermoFisher).

### Cell Culture and Transfection

MCF-7 cells were obtained from the UNC Lineberger Tissue Culture facility and were cultured in DMEM (Sigma or Corning) supplemented with 10% FBS, 100 U/mL penicillin, 100 μg/mL streptomycin (Sigma), 0.01 mg/mL of insulin from bovine pancreas (Sigma), with (Cell Titer Glo assay) or without (ABPP experiment) 1X Minimum Essential Medium Non-Essential Amino Acids Solution (Gibco). Cells were grown and maintained in 10 cm-diameter tissue culture dishes in an incubator at 37°C with 5% CO_2_.

Human cervical cancer cell line HeLa was purchased from ATCC and cultured in DMEM medium supplemented with 10% FBS, 100 Units/mL penicillin and 100 mg/mL streptomycin. Cell transfection was performed using polyethylenimine (23866-1, Polysciences, Inc) according to manufacturer instructions.

### Cysteine-targeted Activity-Based Protein Profiling

#### In-lysate reactive cysteine profiling

The streamlined reactive cysteine profiling was performed as described in previous work.^28,29^ To briefly describe the process, the native and soluble proteome of MCF-7 cells was generated through homogenization by sonication (5 min, 3-s on, 5-s off, 50% amplitude) in native lysis buffer (PBS, pH 7.4, 0.1% NP-40) on ice. To profile reactive cysteine, 30 µg of lysate (2 µg/µL, spiked with recombinant SETDB1 at 0.5% w/w of the total proteome) in 15 µL of lysis buffer was loaded into a 96-well plate. The lysate was treated sequentially with 10 µM of the compound and 500 µM of the DBIA probe for 1 hour respectively at room temperature. The reaction was quenched by adding 3 µL of a 1:1 mixture of SP3 beads (50 mg/mL, Cat. #45152105050250 and Cat. #65152105050250) and 30 µL of ∼98% ethanol containing 20 mM DTT. After a 15-minute incubation, the plate was placed on a magnetic stand, and the liquid was aspirated. The beads were washed once with 200 µL of 80% ethanol, resuspended in 25 µL of lysis buffer containing 20 mM IAA, incubated in the dark for 30 minutes with vigorous shaking, and then 50 µL of ∼98% ethanol containing 20 mM DTT was added. After another 15-minute incubation, the beads were washed twice with 80% ethanol. The remaining beads were resuspended in 30 µL of 200 mM EPPS buffer (pH 8.5) containing 0.3 µg of Lys-C and 0.3 µg of trypsin and incubated at 37 °C overnight. The next day, 60 µg of TMT reagent was added to the mixture of digested peptides and beads to achieve a final acetonitrile concentration of ∼30%, followed by gentle mixing at room temperature for 60 minutes and quenching with 7 µL of 5% hydroxylamine. All TMT-labeled samples were combined, dried using a SpeedVac, and desalted using a 100-mg Sep-Pak column. The labeled peptides were enriched using 80 µL of Pierce™ High-capacity Streptavidin Agarose (Cat. #20359) in 100 mM HEPES buffer (pH 7.4) for 3 hours at room temperature. The remaining agarose resin was washed sequentially with 300 µL of 100 mM HEPES (pH 7.4) with 0.05% NP-40 twice, 350 µL of 100 mM HEPES (pH 7.4) three times, and 400 µL of water once on an Ultrafree-MC centrifugal filter (hydrophilic PTFE, 0.22 µm pore size). The peptides were eluted sequentially with elution buffer (80% acetonitrile, 0.1% formic acid) twice and elution buffer at 72 °C once. The combined eluate was dried, desalted via StageTip, and resuspended in loading buffer (5% ACN and 5% FA) prior to LC-MS analysis.

#### LC-FAIMS-MS/MS analysis

Enriched cysteines were loaded onto a 100-µm capillary column packed with 25 cm of Accucore 150 resin (2.6 μm, 150Å; Thermo Fisher Scientific) and separated using a 180-minute method on a NanoLC-1200 UPLC system. Cysteine data were collected using a high-resolution MS/MS method on an Orbitrap Eclipse mass spectrometer coupled with a FAIMS Pro device. Each cysteine sample was analyzed twice with different sets of FAIMS compensation voltages (CVs): 1) −60, −45, and −35V, and 2) −70, −55, and −30V. MS1 scans were collected in the Orbitrap with 60K resolution, a 400-1600 m/z scan range, standard AGC, and a 50 ms maximum injection time (IT). Data-dependent MS2 scans were acquired in Top Speed mode with a cycle time of 1 second, selecting and fragmenting precursors using 36% HCD. MS2 scans were collected in the Orbitrap with 50K resolution, a fixed 110-2000 m/z scan range, 500% AGC, and a maximum IT of 86 ms. Dynamic exclusion was set to 120 seconds with a mass tolerance of ± 10 ppm. The flowthrough was analyzed using a 60-minute LC-FAIMS-MS/MS in data-dependent analysis, similar to the settings used for analyzing cysteine samples.

#### Data analysis for cysteine identification, localization, and quantification

Raw files were searched using the Comet search engine (ver. 2019.01.5)^30^ with the Uniprot human proteome database (downloaded 11/24/2021). The settings included a 50 ppm precursor error tolerance, a 0.9 Da fragment error tolerance, and peptide modifications such as fixed Cys carboxyamidomethylation (+57.0215), fixed Lys/N-terminal TMTpro (+304.2071), variable Met oxidation (+15.9949), and variable Cys DBIA mass shift (+239.1628) based on carboxyamidomethylation. Peptide spectral matches were filtered to a peptide false discovery rate (FDR) of <1% using linear discriminant analysis and then filtered to obtain a 1% protein FDR at the entire dataset level employing a target-decoy strategy.^31,32^ Cysteine-modified peptides were then filtered for site localization using the AScorePro algorithm with a cutoff of 13 (P < 0.05) as previously described.^33^ Unique peptides and unique cysteine sites were summarized from all peptide spectral matches (PSMs) and reported. For TMT reporter quantification, a total sum signal-to-noise ratio of all reporter ions of 180 was required, with fewer than three missing values. Peptide and cysteine site quantitative values were normalized based on sample loading differences obtained from analyzing the flowthrough sample.

### Cell Titer Glo Proliferation Evaluation

5,000 MCF-7 cells in 25 µL of media were seeded in each well of a 384-well white tissue culture-treated plate (Corning #3570). After 24 h, the media was replaced with media containing the desired compound concentration to obtain a dose-response. Treated cells were then incubated for 24 h, 48 h, or 72 h at 37 °C. Cell proliferation analysis was performed using the Cell Titer Glo Kit (G9241, Promega) according to the manufacturer’s instructions. Luminescent signals were measured using the Promega GloMax Plate Reader with an integration time of 0.3 seconds per well. Percent viability was calculated as a percentage of the related DMSO-treated cells.

### Caco-2 Cell Permeability Evaluation

The Caco-2 cell permeability evaluation data was obtained from the *In vitro* ADME laboratory at Pharmaron (Beijing, China).

### Cellular Thermal Shift Assay

SETDB1 (3TD, 197-403aa) was cloned into pBiT3.1-N (Promega, #N2361) and pBiT3.1-C (Promega, #N2371) plasmids. HEK293T cells were plated in 6-well plates (1e6/well) and reversely transfected with 0.2 µg HiBIT –tagged SETDB1 (3TD) and 1.8 µg of empty plasmid using X-tremeGene XP transfection reagent (Roche), following manufacturer’s instructions. The next day cells were trypsinized and resuspended in OptiMEM (no phenol red, Gibco) at density 2e5/ml. After compound or DMSO addition (the same DMSO concentration in each sample) cells were transferred to 96-well pcr plates (40 µl/well), incubated for 1 h at 37 °C and heated for 3 min. For permeabilized CETSA cells were incubated in the presence of digitonin (50 ug/ml) for 20 min at RT. After 3 min, 40 µl of LgBIT solution (200 nM LgBIT, 2% NP-40, protease inhibitors in OptiMEM no phenol red) was added and incubated at RT for 10 min. Next, 20 µl of NanoGlo substrate (8 µl/ml, Promega) was added, mixed gently and 20 µl was transferred to 384 white plates in triplicates and the bioluminescence signal was read in the ClarioStar plate reader. The results and mean +/- s.d.

### Cellular Akt Methylation and Phosphorylation Evaluation

#### Plasmid construction

pcDNA3-HA-Akt1 was constructed by cloning corresponding PCR fragments into pcDNA3-HA vector by BamHI and SalI sites.

Akt1-BamHI-Forward: 5′-GCATGGATCCAGCGACGTGGCTATTGTG-3′ Akt1-SalI-Reverse: 5′-GCATGTCGACTCAGGCCGTGCCGCTGGC-3′

#### Antibodies

All antibodies were used at a 1:1,000 dilution in TBST buffer with 5% non-fat milk for western blotting unless specified. Anti-HA-Tag antibody (3724), anti-Tri-Methyl Lysine motif antibody (14680) and Anti-phospho-Akt-Thr308 antibody (9275) were obtained from Cell Signaling Technology. Ant-Vinculin antibody (sc-25336) was purchased from Santa Cruz Biotechnology.

#### Immunoblot and Immunoprecipitations Analyses

Cells were lysed in EBC buffer (50 mM Tris pH 7.5, 120 mM NaCl, 0.5% NP-40) or Triton X-100 buffer (50 mM Tris, pH 7.5, 150 mM NaCl, 1% Triton X-100) supplemented with protease inhibitor cocktail and phosphatase inhibitor cocktail. The protein concentrations of whole cell lysates were measured by NanoDrop OneC using the Bio-Rad protein assay reagent. Equal amounts of whole cell lysates were loaded by SDS-PAGE and immunoblotted with indicated antibodies. For immunoprecipitations analysis, 1 mg total lysates were incubated with the anti-HA agarose beads (A-2095, Sigma) for 3-4 hr at 4°C. The recovered immuno-complexes were washed three times with NETN buffer (20 mM Tris, pH 8.0, 100 mM NaCl, 1 mM EDTA and 0.5% NP-40) before being resolved by SDS-PAGE and immunoblotted with indicated antibodies.

## EXTENDED DATA

**Extended Data Figure 1.**
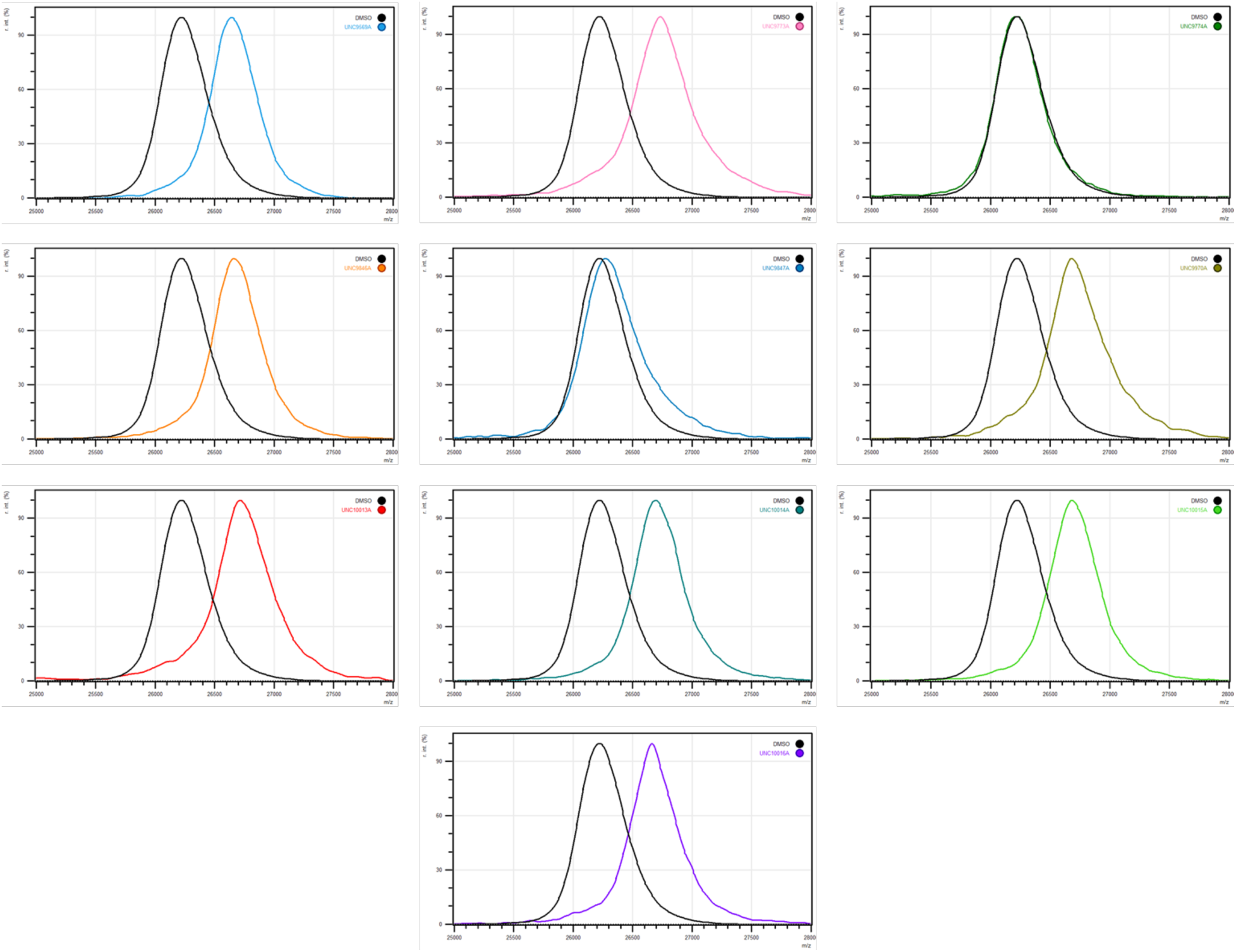
MALDI-TOF evaluation of the ability of compounds 1-9 to covalently modify SETDB1. Representative MALDI-TOF qualitative evaluation of the ability of **1**-**9** to covalently modify SETDB1 3TD after 2 h incubation at room temperature using 10 µM of protein and 100 µM of ligand.

**Extended Data Figure 2.**
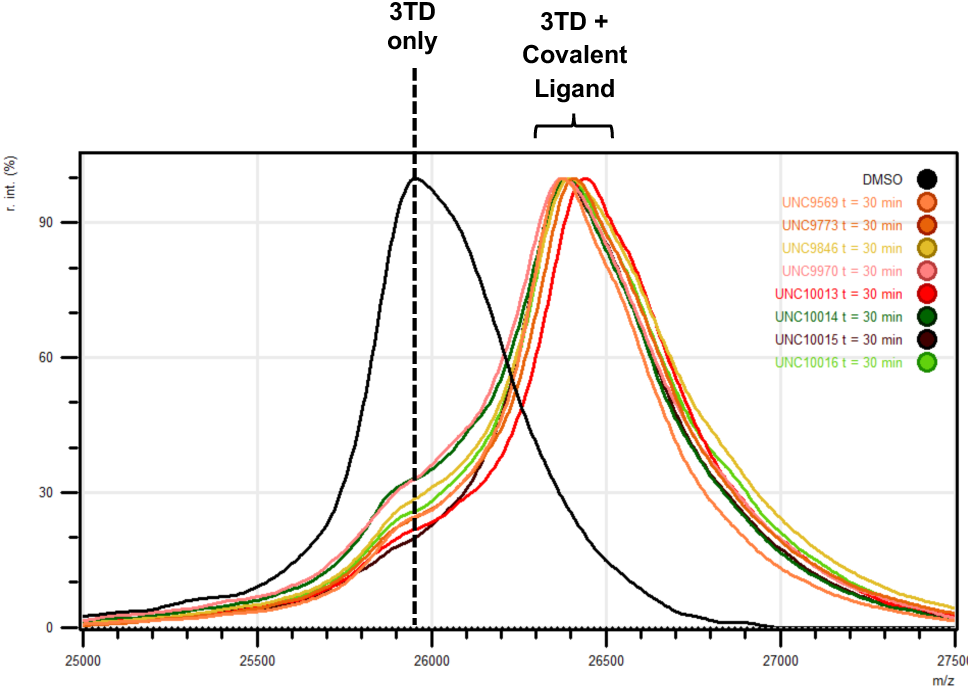
MALDI-TOF evaluation of the ability of compounds 1-8 to covalently modify SETDB1 after 30 minutes incubation on ice. Representative MALDI-TOF qualitative evaluation of the ability of **1**-**8** to covalently modify SETDB1 3TD after 30 minutes incubation on ice using 10 µM of protein and 10 µM of ligand.

**Extended Data Figure 3.**
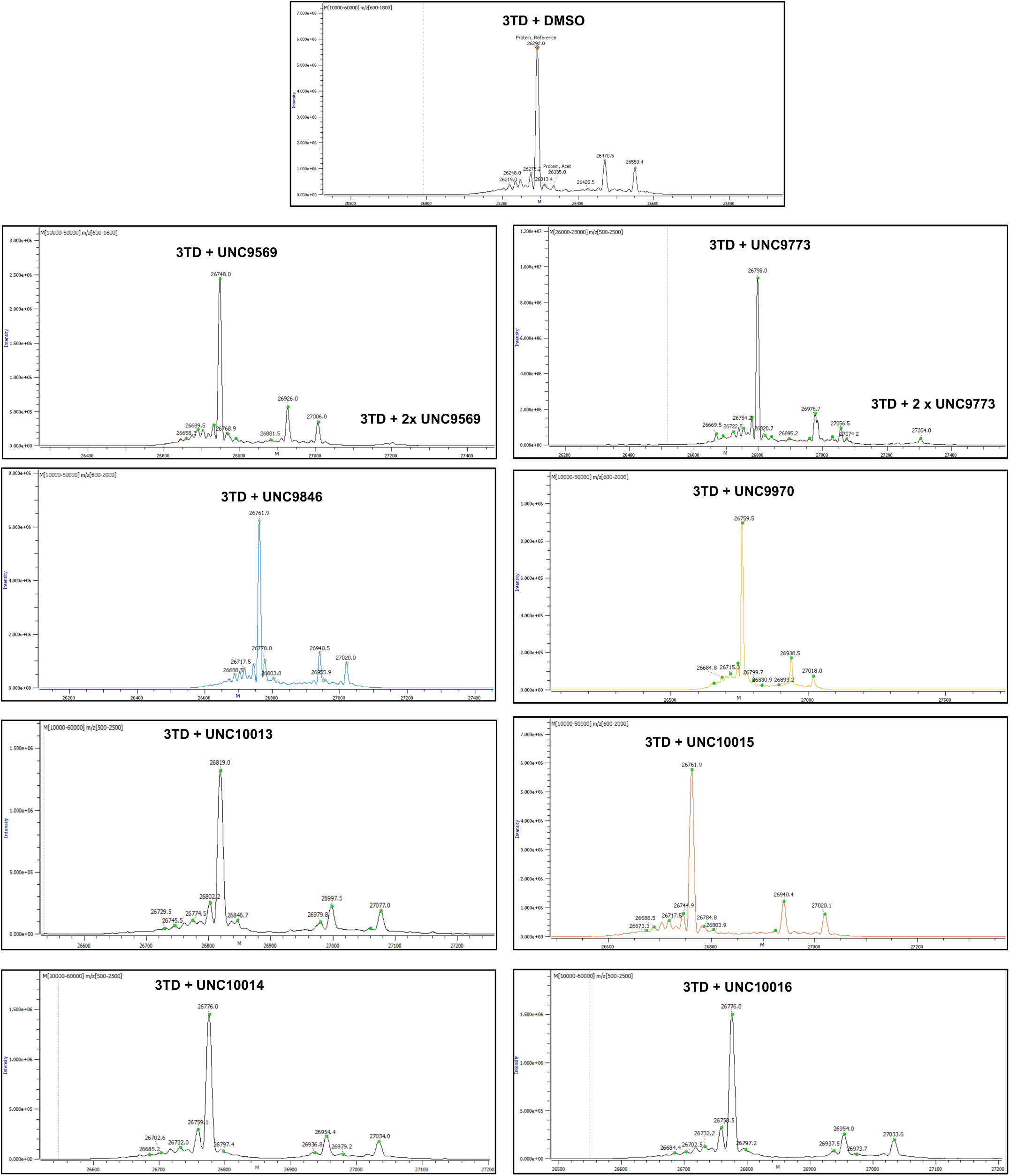
High resolution intact protein mass spectrometry to evaluate the ability of compounds 1-8 to covalently modify SETDB1. Representative deconvoluted mass spectra for evaluation of the ability of **1**-**8** to modify SETDB1 3TD.

**Extended Data Figure 4.**
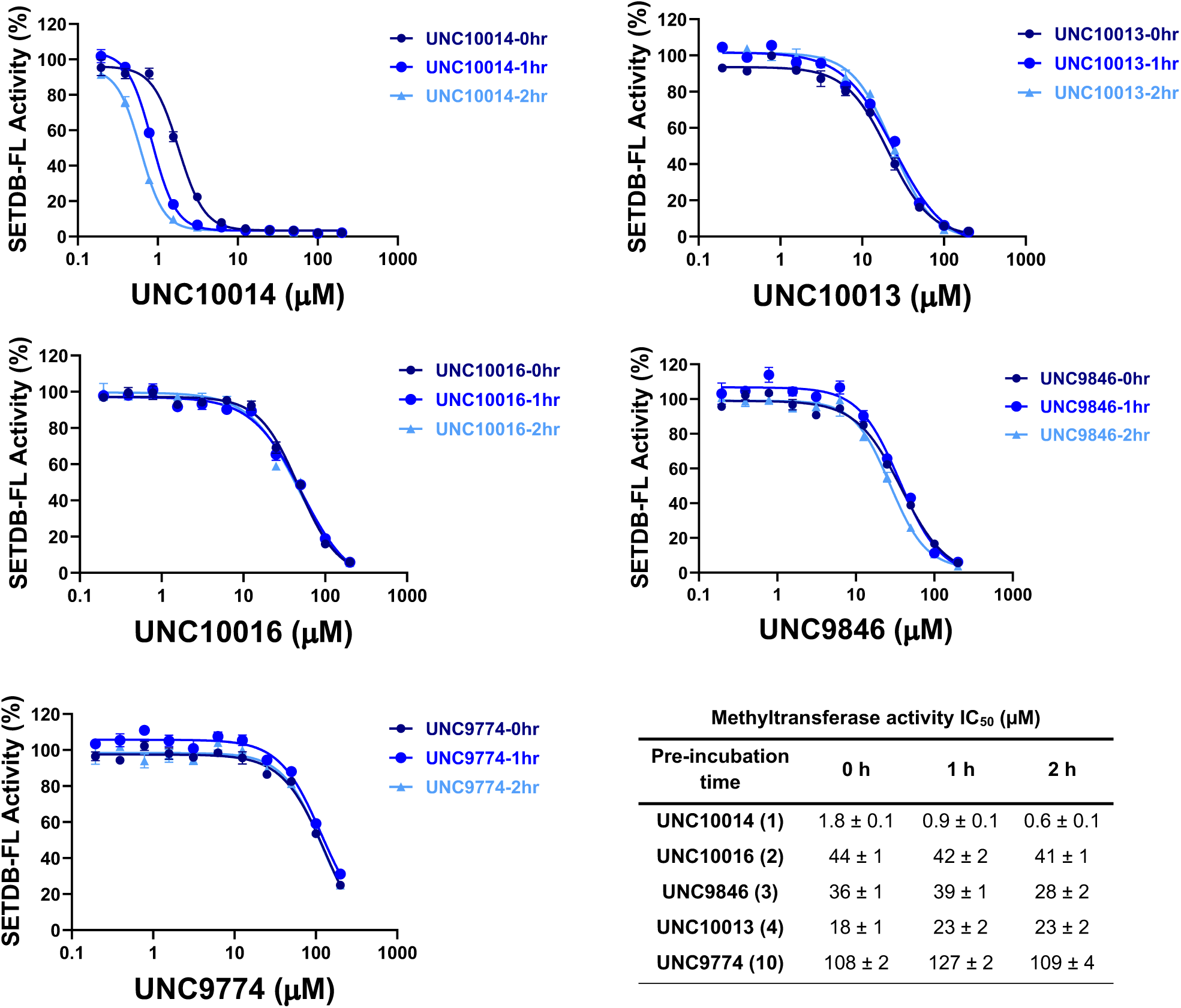
Evaluation of the ability of compounds 1-4 and 10 to inhibit the methyltransferase activity of SETDB1. Compounds **1**-**4** and **10** were preincubated for 0, 1 or 2 h with the catalytically active full-length SETDB1 protein before measurement of the activity inhibition using a radiometric methyltransferase assay. Values are reported as the average of three independent experiments ± s.d.

**Extended Data Figure 5.**
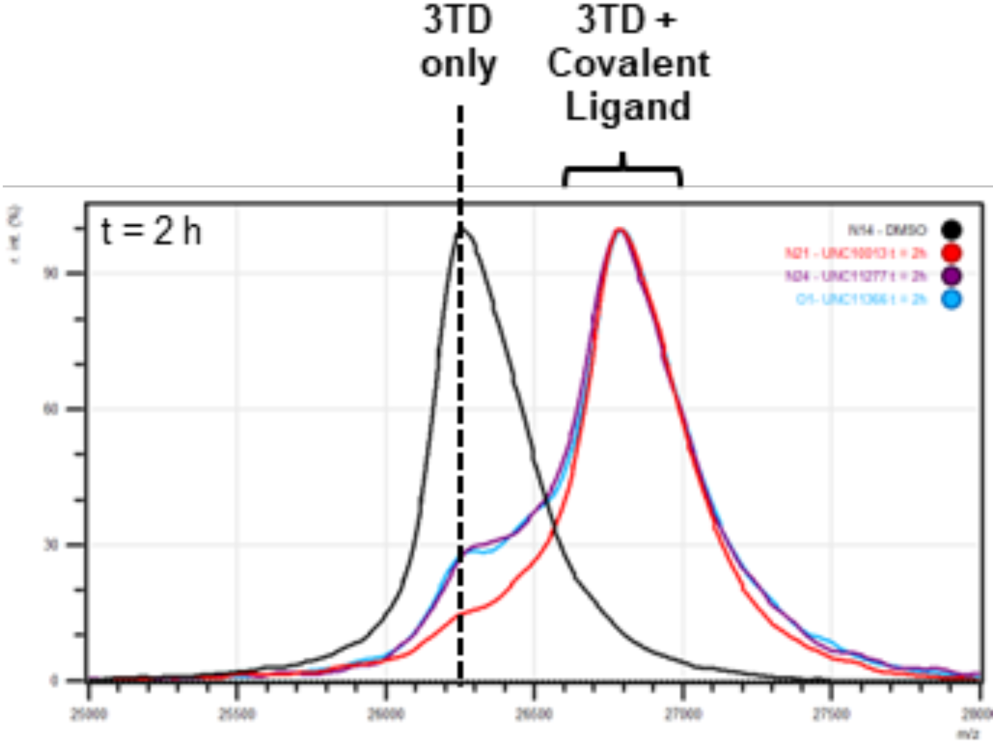
MALDI-TOF qualitative evaluation of the ability of UNC11277 (11) and UNC11366 (12) to covalently modify the 3TD of SETDB1. Representative MALDI-TOF spectra after incubation of 10 µM of UNC10013 (**4**) (red) **UNC11277** (**11**) (purple), and **UNC11366** (**12**) (blue) incubated with 10 µM of SETDB1 3TD for t = 2 h on ice.

**Extended Data Figure 6.**
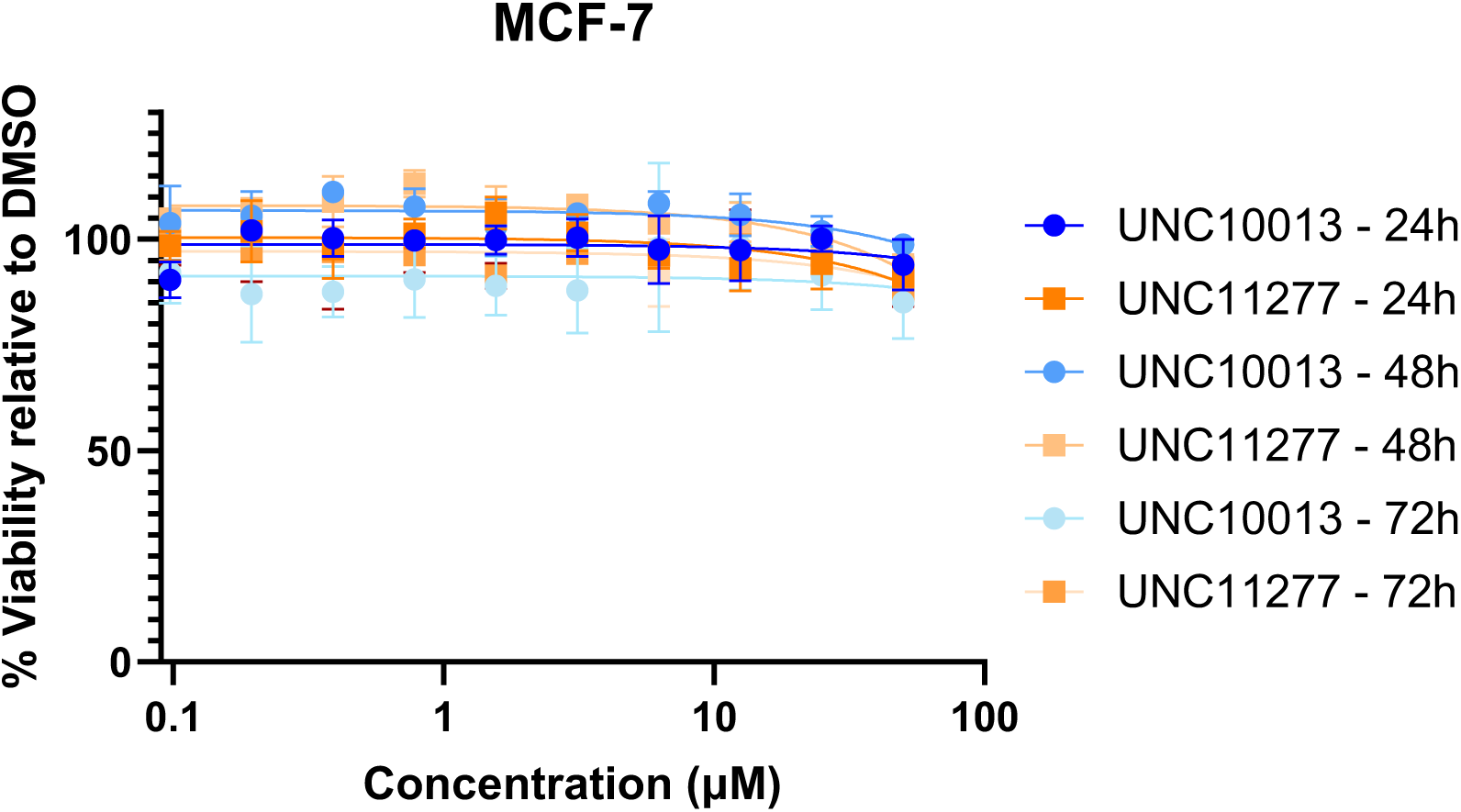
Graph showing the results of a Cell Titer Glo assay to evaluate the toxicity of UNC10013 (4) and UNC11277 (11) treatment of MCF-7 cells compared to DMSO. Concentrations up to 50 µM were tested for up to 72 h treatment. Results shown are the average of three independent replicates ± s.e.m.

## ACKNOWLEDGMENT

This work was supported by the PhRMA Foundation (Drug Discovery postdoctoral fellowship) to M.U. and the National Institute of General Medical Sciences, NIH (grant R35GM139514) to S.V.F. This work was also supported by the National Cancer Institute of the National Institutes of Health under award number P30CA016086. The content is solely the responsibility of the authors and does not necessarily represent the official views of the National Institutes of Health. The authors thank Juanita Rubiano Sánchez and Tiffany Peters for reviewing the experimental chemistry and biology data. The authors thank B. Hardy for assembling the screening plate for TR-FRET. Part or all of the research described in this paper was performed using beamline CMCF-ID at the Canadian Light Source, a national research facility of the University of Saskatchewan, which is supported by the Canada Foundation for Innovation (CFI), the Natural Sciences and Engineering Research Council (NSERC), the National Research Council (NRC), the Canadian Institutes of Health Research (CIHR), the Government of Saskatchewan, and the University of Saskatchewan. The Structural Genomics Consortium is a registered charity (no. 1097737) that receives funds from Bayer AG, Boehringer Ingelheim, Bristol Myers Squibb, Genentech, Genome Canada through Ontario Genomics Institute [OGI-196], EU/EFPIA/OICR/McGill/KTH/Diamond Innovative Medicines Initiative 2 Joint Undertaking [EUbOPEN grant 875510], Janssen, Merck KGaA (aka EMD in Canada and US), Pfizer, and Takeda.

