## Supplementary Information for "Potent and Selective SETDB1 Covalent Negative Allosteric Modulator Reduces Methyltransferase Activity in Cells"

### 1. Chemistry

#### 1.1. General Information

Chemicals were purchased from commercial suppliers and used without further purification. Thin layer chromatography (TLC) was performed on glass plates coated with 60 F254 silica. Flash chromatography was carried out using a Teledyne Isco Combiflash Rf200, Rf200i or NextGen 300+ automated flash system with RediSep Rf normal phase or C18 RediSep Rf Gold reverse phase silica gel pre-packed columns. Fractions were collected at 220 nm and/or 254 nm. Preparative HPLC was performed using an Agilent Prep 1200 series with the UV detector set to 220 nm and 254 nm. Samples were injected onto either a Phenomenex Luna 250 × 30 mm (5 μm) C18 column or a Phenomenex Luna 75 x 30 mm (5 μm) C18 column at room temperature. Microwave irradiation was performed in a CEM Discover SP microwave reactor in sealed vials. Reactions were irradiated at temperatures between 60 and 250 °C using low or medium absorbance mode depending on the solvent.

#### 1.2. Analytical Equipment

$^{1}$H NMR spectra were obtained using a Varian 400MR Inova spectrometer using a frequency of 400 MHz. $^{13}$C spectra were acquired using the Varian 400MR Inova spectrometer operating at a frequency of 101 MHz or a Bruker Avance III HD 700 MHz at a frequency of 176 MHz. The abbreviations for spin multiplicity are as follows: s = singlet; d = doublet; t = triplet; q = quartet, p = quintuplet, h = sextuplet and m = multiplet. Combinations of these abbreviations are employed to describe more complex splitting patterns (e.g. dd = doublet of doublets). Analytical LCMS data for all compounds was acquired using an Agilent 6110 Series system with the UV detector set to 254 nm. Samples were injected (<10 μL) onto an Agilent Eclipse Plus 4.6 × 50 mm, 1.8 μm, C18 column at room temperature. Mobile phases A (H2O + 0.1% acetic acid) and B (MeCN + 1% H2O + 0.1% acetic acid) were used with a linear gradient from 10% to 100% B in 5.0 min, followed by a flush at 100% B for another 2 minutes with a flow rate of 1.0 mL/min. Mass spectra (MS) data were acquired in positive ion mode using an Agilent 6110 single quadrupole mass spectrometer with an electrospray ionization (ESI) source. Analytical LCMS (at 254 nm) was used to establish the purity of targeted compounds. All compounds that were evaluated in biochemical and biophysical assays had >95% purity as determined by LCMS.

#### 1.3. General Procedures

*General Procedure 1: Amide formation using acyl chloride reagent.*

In a round-bottom flask containing the aniline derivative (1.0 eq.) in 3:1 DCM/DMF (33 mL/mmol) cooled down to 0 °C using an ice bath, the acyl chloride reagent (1.2 eq.) followed by triethylamine (2.0 eq.) were added. The mixture was stirred at 0 °C for 30 min and then at room temperature for another 30 min. Methanol was then added to the mixture and the solvent was removed under reduced pressure.

*General Procedure 2: Amide formation with in situ acyl chloride formation.*

In a round-bottom flask containing the carboxylic acid derivative (1.5 eq.) in THF (4.2 mL/mmol) cooled down to 0 °C using an ice bath, oxalyl chloride (1.5 eq.) was added dropwise followed by a catalytic amount of DMF. The mixture was stirred at 0 °C for 15 min before being stirred at room temperature for another 2 h. Then, the mixture was cooled down to 0 °C before the aniline derivative (1.0 eq.) dissolved in THF (8.4 mL/mmol) was added dropwise followed by DIPEA (2.0 eq.). The mixture was then stirred at room temperature for 1 h.

#### 1.4. Synthetic Schemes

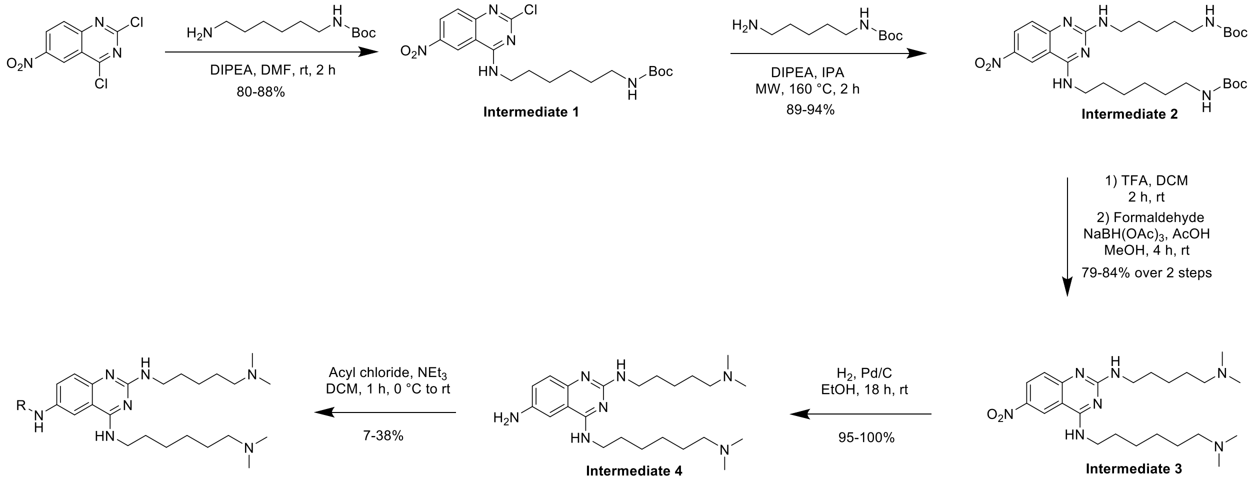

**Scheme S1.** General synthetic route for the synthesis of **1-10**.

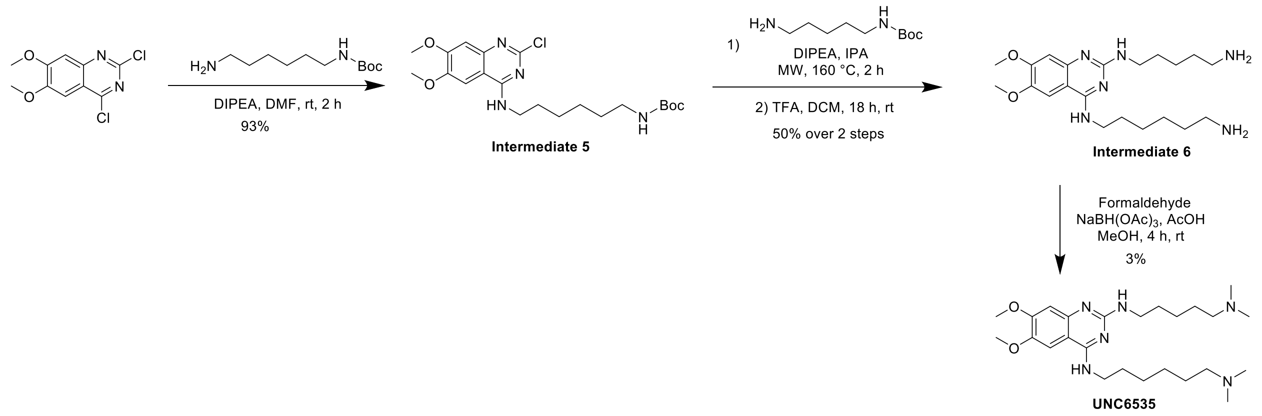

**Scheme S2.** Synthetic route for the synthesis of **UNC6535**.

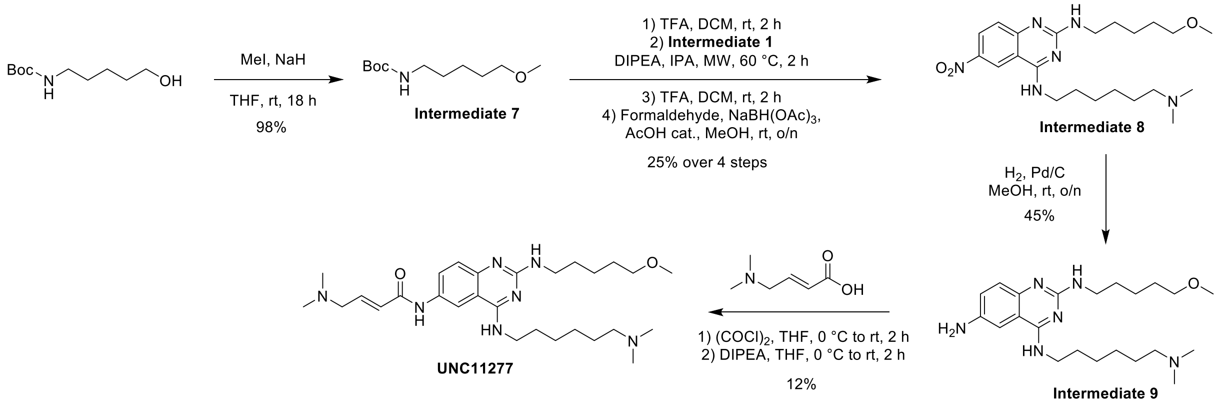

**Scheme S3.** Synthetic route for the synthesis of **UNC11277**.

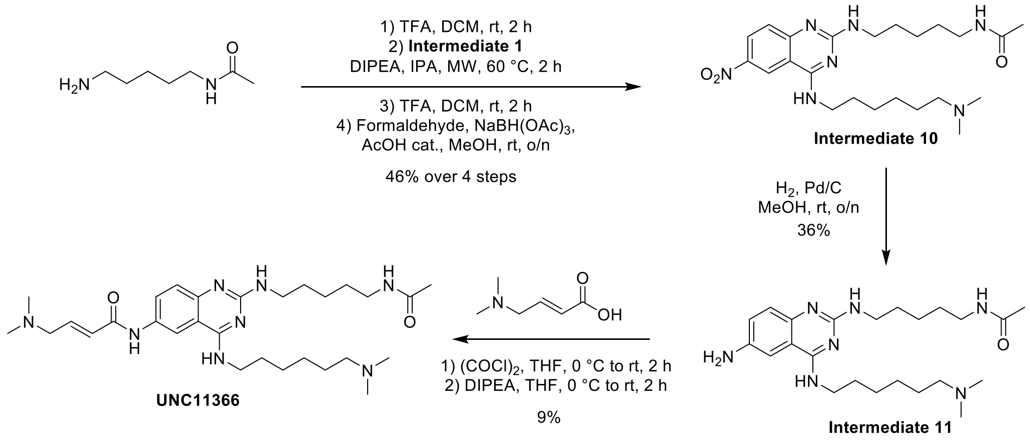

**Scheme S4.** Synthetic route for the synthesis of **UNC11366**.

##### 1.5. Compound Data

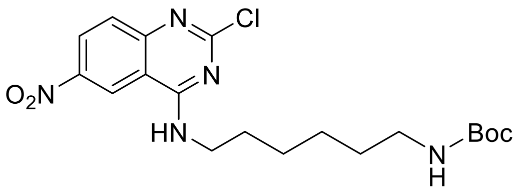

***tert*-butyl (6-((2-chloro-6-nitroquinazolin-4-yl)amino)hexyl)carbamate (Intermediate 1)**

In a round-bottom flask containing 2,4-dichloro-7-nitroquinazoline (200 mg, 1.0 eq.) dissolved in 1.0 mL of DMF, *N*-Boc-1.6-diaminohexanediamine (202 µL, 1.1 eq.) and DIPEA (286 µL, 2.0 eq.) were added. The mixture was stirred at room temperature for 2 h. The solvent was removed by reduced pressure. Flash chromatography (0 to 60% EtOAc in Hexane) yielded the desired product as a yellow solid (305 mg, 88%).$^{1}$H NMR (400 MHz, $CDCl_3$) δ 9.11 (s, 1H), 8.48 (dd, *J* = 9.2, 2.4 Hz, 1H), 7.81 (d, *J* = 9.2 Hz, 2H), 4.71 – 4.61 (m, 1H), 3.70 (q, *J* = 6.1 Hz, 2H), 3.24 (q, *J* = 6.2 Hz, 2H), 1.76 (p, *J* = 6.3 Hz, 2H), 1.54 – 1.47 (m, 4H), 1.45 (s, 8H), 1.44 – 1.39 (m, 2H). MS(ES+) m/z 424.2 [M + H]$^{+}$.

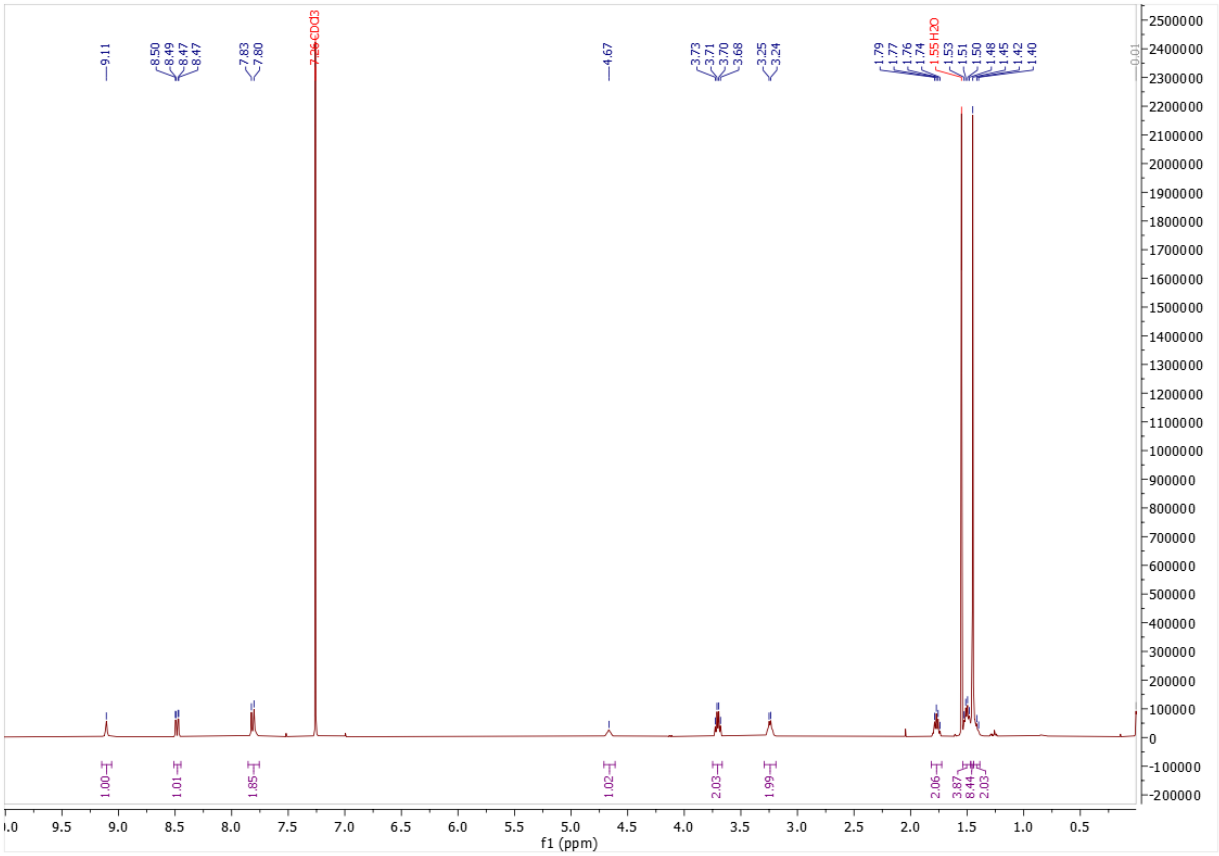

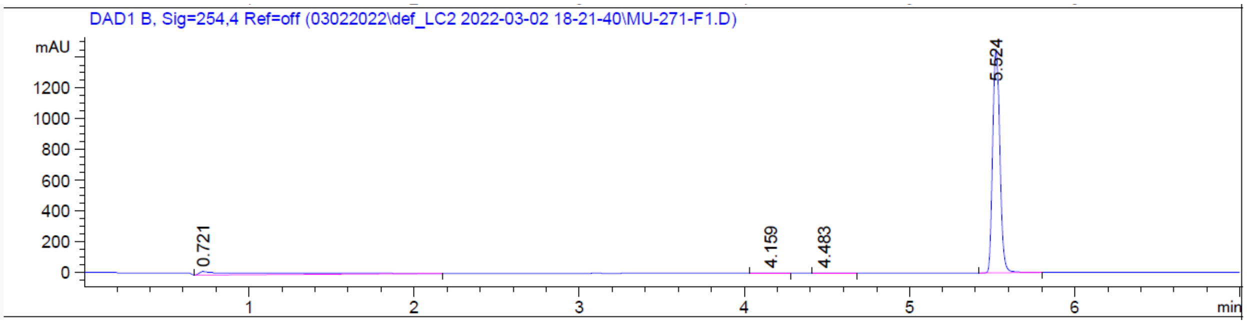

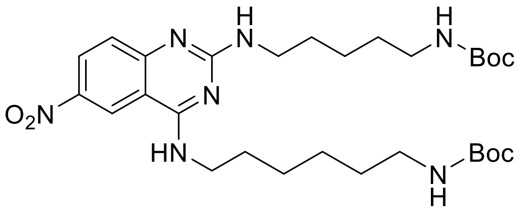

***tert*-butyl (6-((2-((5-((*tert*-butoxycarbonyl)amino)pentyl)amino)-6-nitroquinazolin-4-yl)amino)hexyl)carbamate (Intermediate 2)**

In a microwave vial, **Intermediate 1** (284 mg, 1.0 eq.), *N*-Boc-1,5-diaminoheptane (167 µL, 1.2 eq.) and DIPEA (233 µL) were dissolved in 3.3 mL of IPA. The mixture was heated at 160 °C for 2 h under microwave irradiation. The solvent was removed under reduced pressure. Flash chromatography (0 to 20% MeOH in DCM) yielded the desired product as a yellow sticky oil (351 mg, 89%). $^1$H NMR (400 MHz, cdcl$_3$) δ 8.70 – 8.58 (m, 1H), 8.28 (dd, *J* = 9.3, 2.4 Hz, 1H), 7.39 (s, 2H), 5.34 – 5.17 (m, 0H), 4.69 – 4.47 (m, 2H), 3.71 – 3.56 (m, 2H), 3.52 (q, *J* = 6.7 Hz, 2H), 3.22 – 3.05 (m, 4H), 1.77 – 1.69 (m, 2H), 1.69 – 1.63 (m, 2H), 1.53 – 1.48 (m, 4H), 1.44 (d, *J* = 3.7 Hz, 22H). MS(ES+) m/z 590.3 [M + H]$^+$.

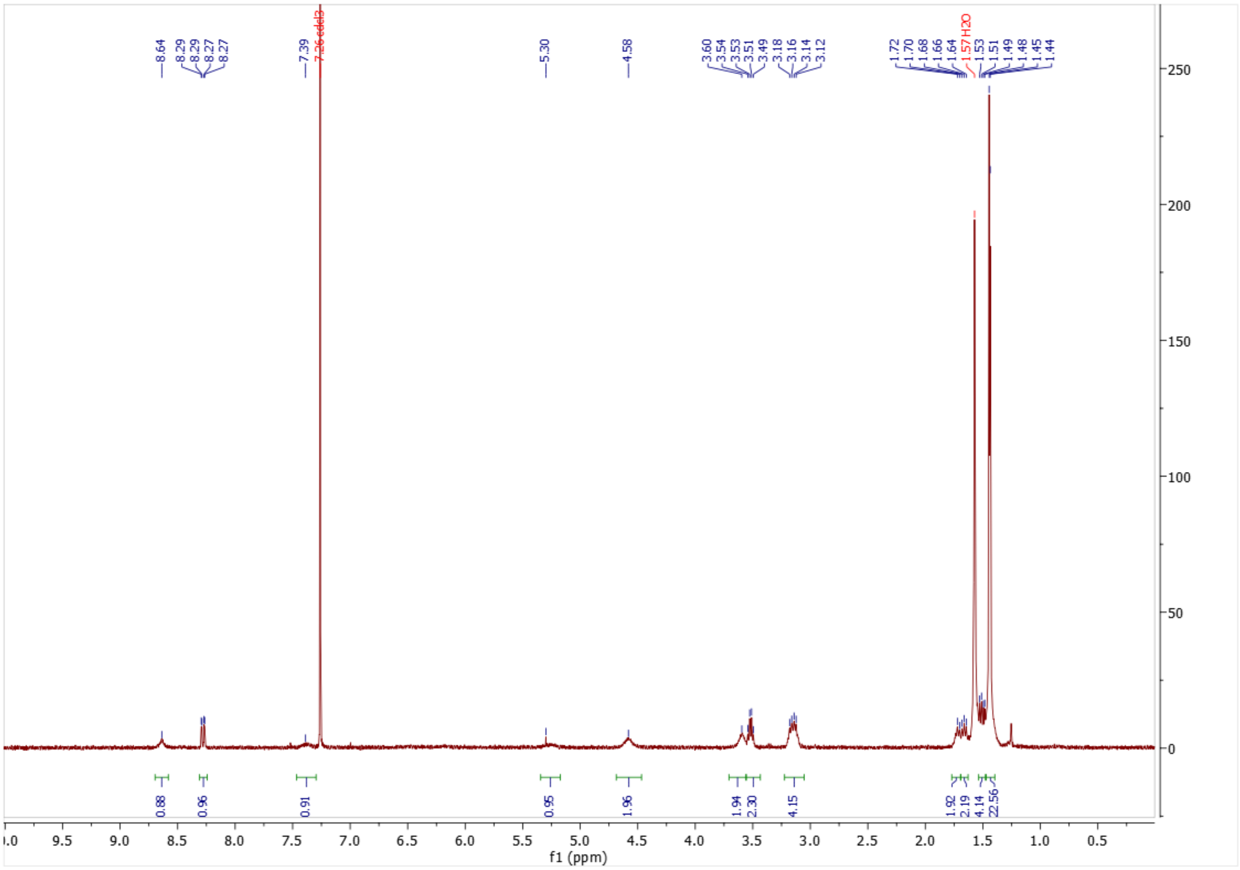
f1 (ppm)

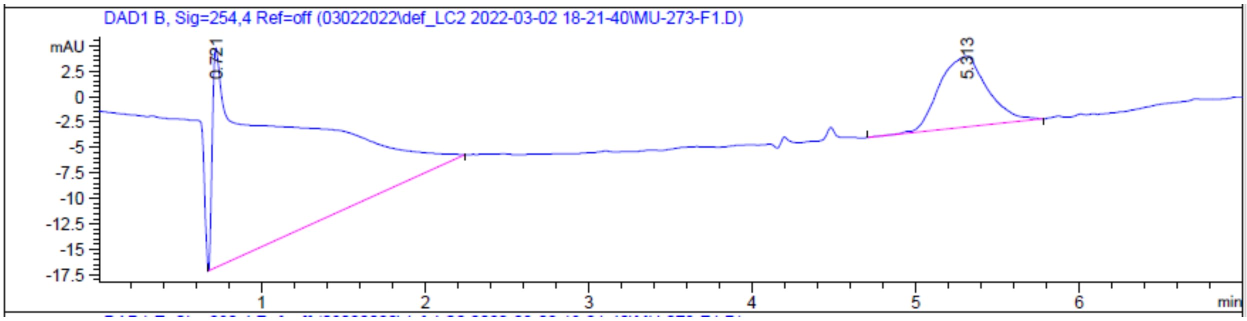
DAD1 B, Sig=254,4 Ref=off (03022022\def_LC2 2022-03-02 18-21-40\MU-273-F1.D)
mAU
0.721
5.313
min

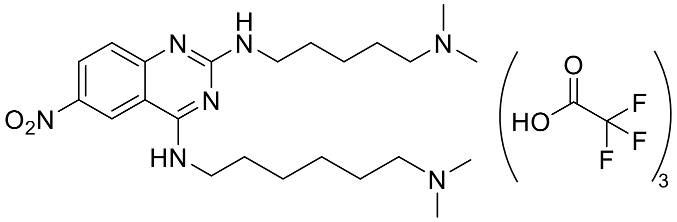

**$N^4$-(6-(dimethylamino)hexyl)-$N^2$-(5-(dimethylamino)pentyl)-6-nitroquinazoline-2,4-diamine • 3 TFA salts (Intermediate 3)**

In a round-bottom flask containing **Intermediate 2** (348 mg, 1.0 eq) in 4.9 mL of DCM, TFA (455 µL, 10 eq.) was added. The mixture was stirred at room temperature overnight. The solvent was removed under pressure. The crude mixture was dissolved in 2.8 mL of MeOH. Formaldehyde 37% in $H_2O$ solution (264 µL, 6.0 eq.) was added. The mixture was stirred at room temperature for 1 h before sodium triacetoxyborohydride (1.67g, 8.0 eq.) was added partwise. The mixture was stirred at room temperature overnight. Water and MeOH were then added to the mixture before flash chromatography loading. Flash chromatography (5 to 100% MeOH in 0.1% TFA in H2O) yielded the desired product as a yellow oil (342 mg, 74%) as a TFA salt. $^1$H NMR (400 MHz, cd$_3$od) δ 9.17 (d, *J* = 2.3 Hz, 1H), 8.53 (dd, *J* = 9.1, 2.4 Hz, 1H), 7.57 (d, *J* = 9.2 Hz, 1H), 3.74 (t, *J* = 7.1 Hz, 2H), 3.63 (t, *J* = 6.7 Hz, 2H), 3.22 – 3.10 (m, 4H), 2.91 (s, 6H), 2.90 (s, 7H), 1.88 – 1.72 (m, 8H), 1.60 – 1.42 (m, 6H). $^{13}$C NMR (101 MHz, cd$_3$od) δ 163.27 (TFA), 162.91 (TFA), 161.37, 155.22, 144.94, 130.31, 121.95, 119.56 (TFA), 118.90, 116.63 (TFA), 114.61, 111.00, 58.88, 58.83, 43.39, 43.35, 43.18, 42.13, 29.39, 29.20, 27.66, 27.21, 25.55, 25.22, 24.70. MS(ES+) m/z 446.3 $[M + H]^+$.

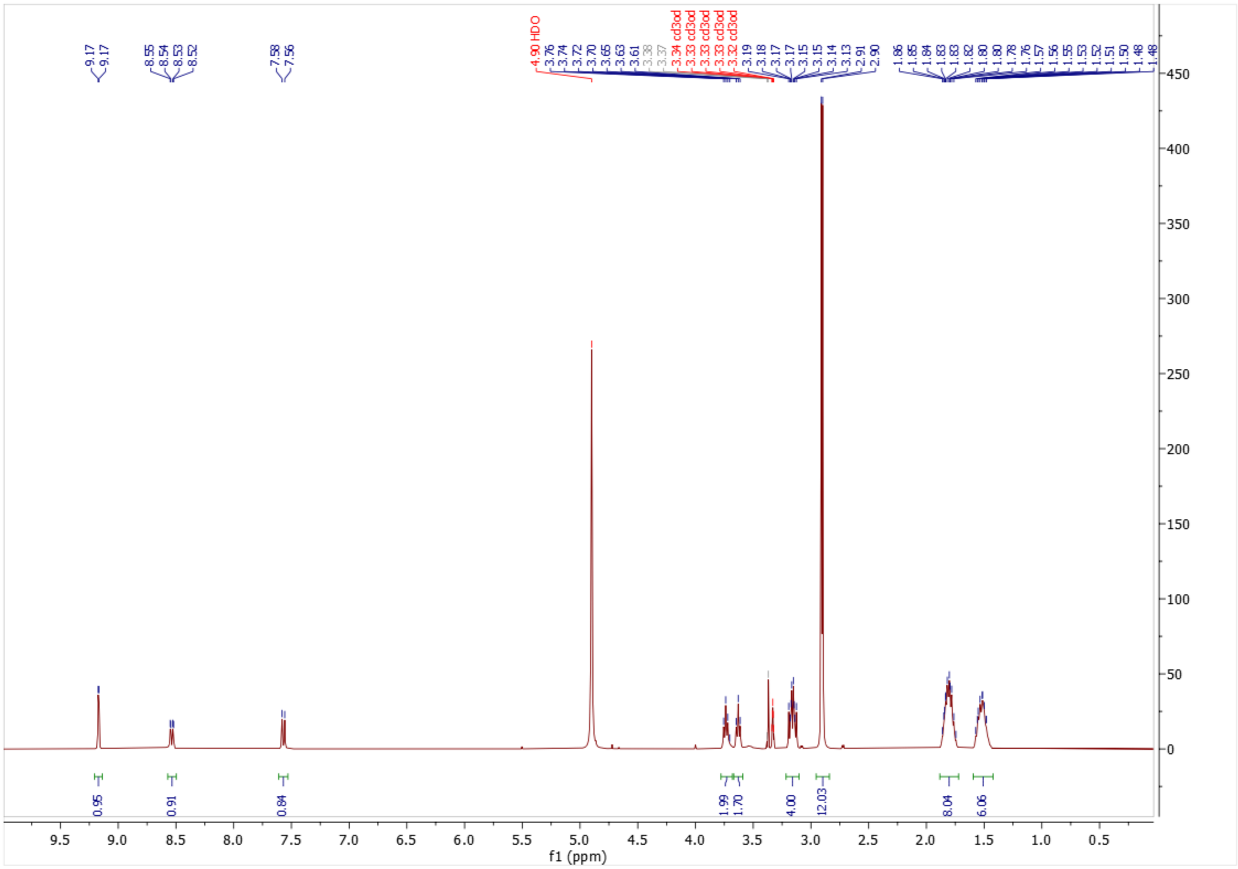

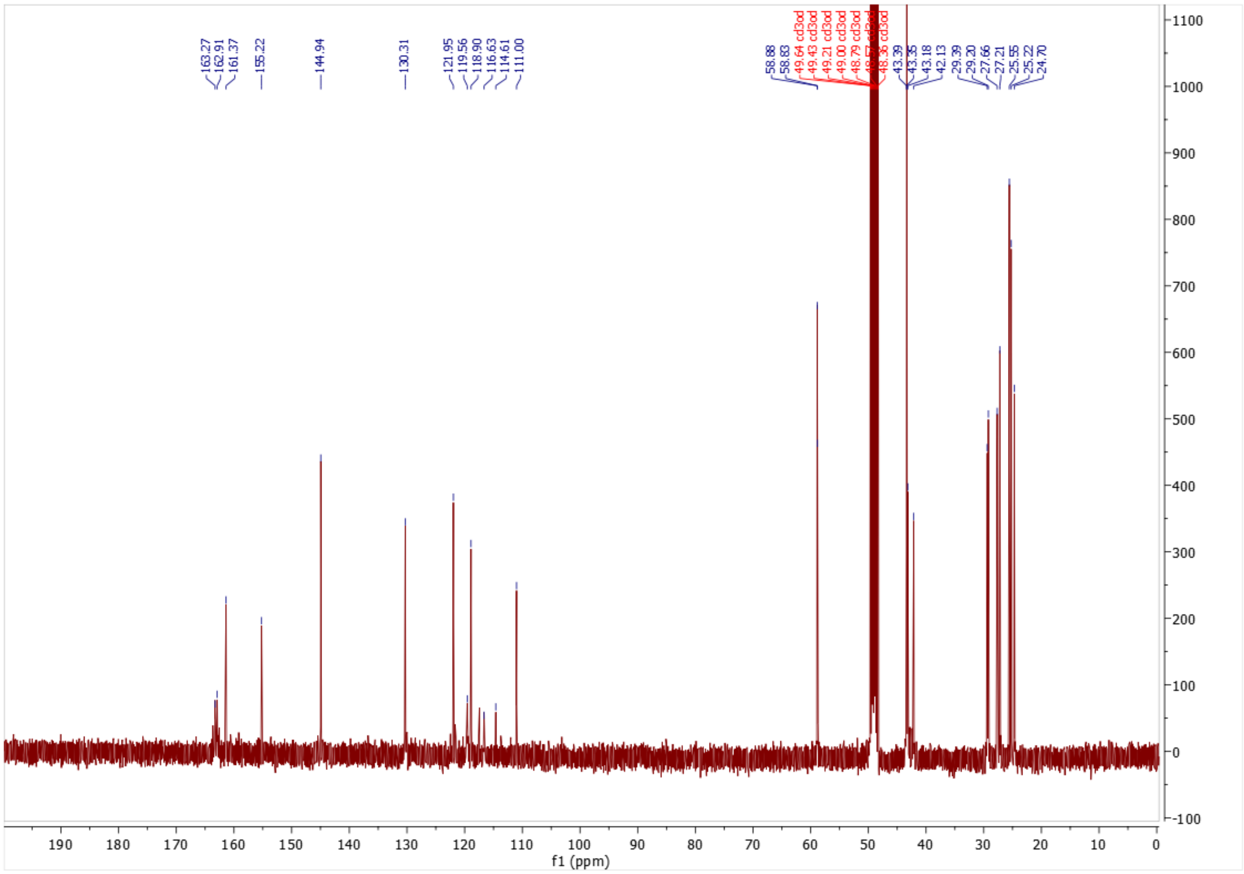

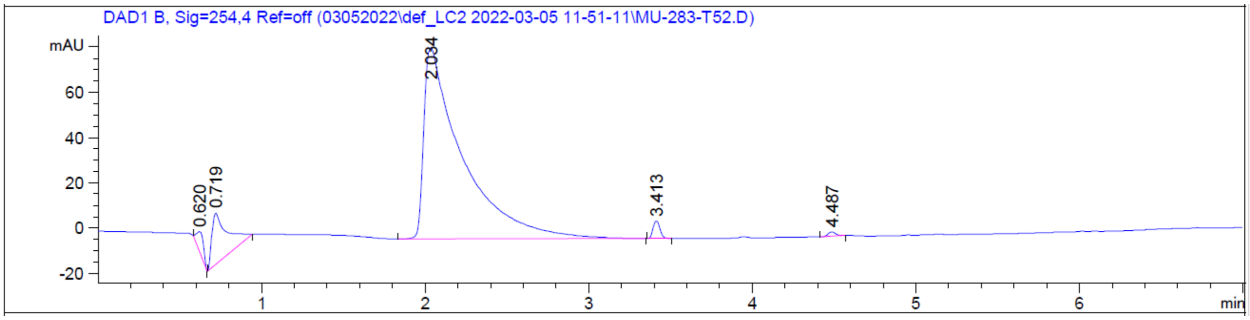

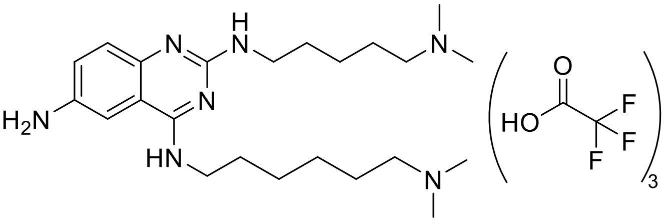

**$N^4$-(6-(dimethylamino)hexyl)-$N^2$-(5-(dimethylamino)pentyl)quinazoline-2,4,6-triamine • 3 TFA salts (Intermediate 4)**

In a round-bottom flask containing **Intermediate 3** (342 mg, 1.0 eq.) in 3.7 mL of MeOH, 10 % palladium on carbon (82 mg, 0.1 eq.) was added. A $H_2$ balloon was fitted on the

flask after removing the air. The mixture was stirred at room temperature overnight. After reaction completion, the mixture was filtered over celite using MeOH for rinsing and then the solvent was removed under reduced pressure. This yielded the desired compound as a brown oil (556 mg, 96%) as a TFA salt. $^{1}$H NMR (400 MHz, cd$_3$od) δ 7.36 (s, 1H), 7.33 – 7.22 (m, 2H), 3.66 (t, *J* = 7.1 Hz, 2H), 3.51 (t, *J* = 7.0 Hz, 2H), 3.18 – 3.07 (m, 4H), 2.88 (s, 6H), 2.87 (s, 6H), 1.84 – 1.68 (m, 8H), 1.54 – 1.39 (m, 6H). MS(ES+) m/z 416.4 [M + H]$^+$.

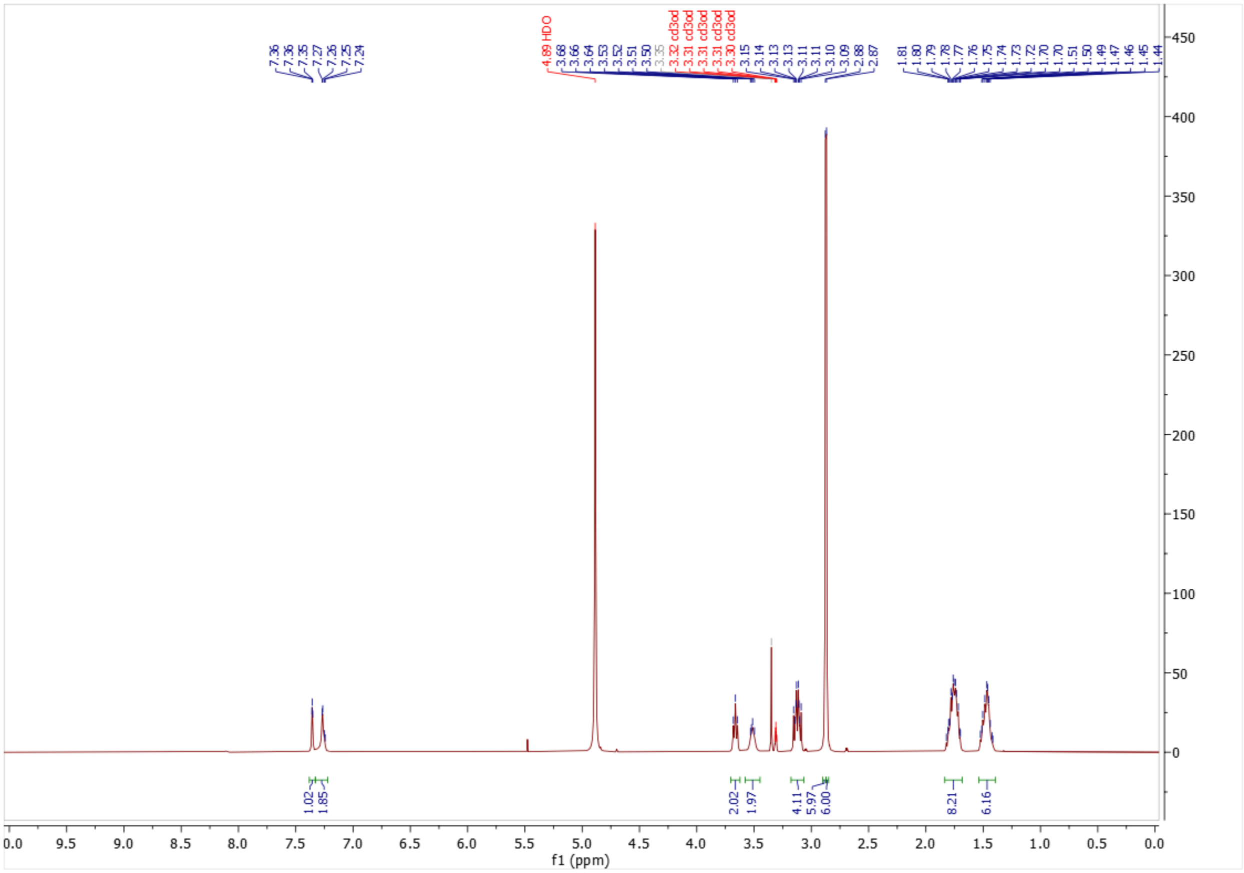

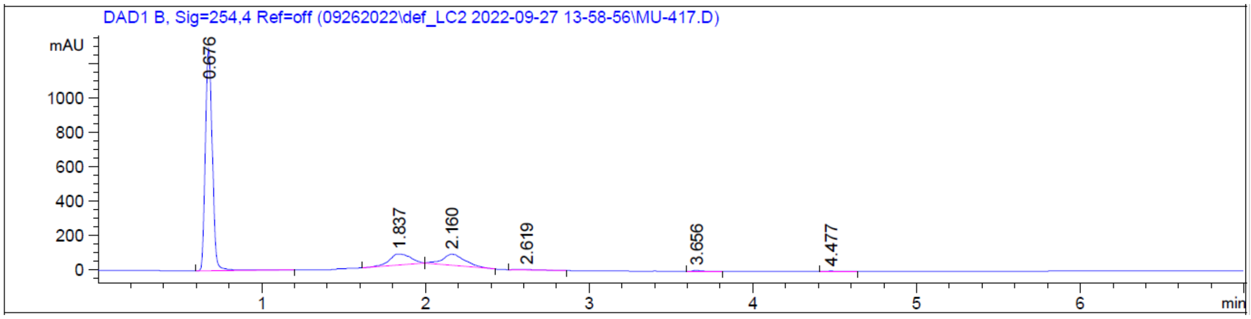

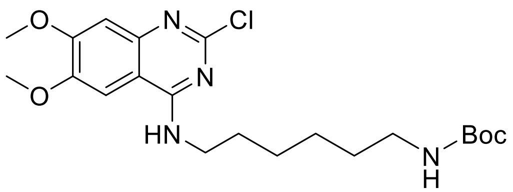

**tert-butyl (6-((2-chloro-6,7-dimethoxyquinazolin-4-yl)amino)hexyl)carbamate (Intermediate 5)**

In a scintillation vial containing 2,4-dichloro-6.7-dimethoxyquinazoline (100 mg, 1.0 eq.) in 0.48 mL of DMF, *N*-Boc-1,6-diaminohexane (95 µL, 1.1 eq.) and DIPEA (134 µL, 2.0 eq.) were added. The mixture was stirred at room temperature for 2 h. The solvent was removed under pressure. Flash chromatography (0 to 100% EtOAc in Hex) yielded the desired product as a white solid (158 mg, 93%). $^{1}$H NMR (400 MHz, $CDCl_3$) δ 7.43 (s, 1H), 7.09 – 7.02 (m, 1H), 3.92 (d, *J* = 7.2 Hz, 6H), 3.65 – 3.55 (m, 2H), 3.12 (q, *J* = 6.5 Hz, 2H), 1.65 (p, *J* = 6.9 Hz, 2H), 1.50 – 1.40 (m, 11H), 1.39 – 1.25 (m, 4H). $^{13}$C NMR (100 MHz, $CDCl_3$) δ 160.25, 156.55, 156.19, 154.60, 148.78, 147.48, 107.24, 106.49, 101.21, 79.23, 56.15, 56.04, 40.58, 39.71, 29.95, 28.49, 28.38, 25.55, 25.26. MS(ES+) m/z 439.2 $[M + H]^{+}$.

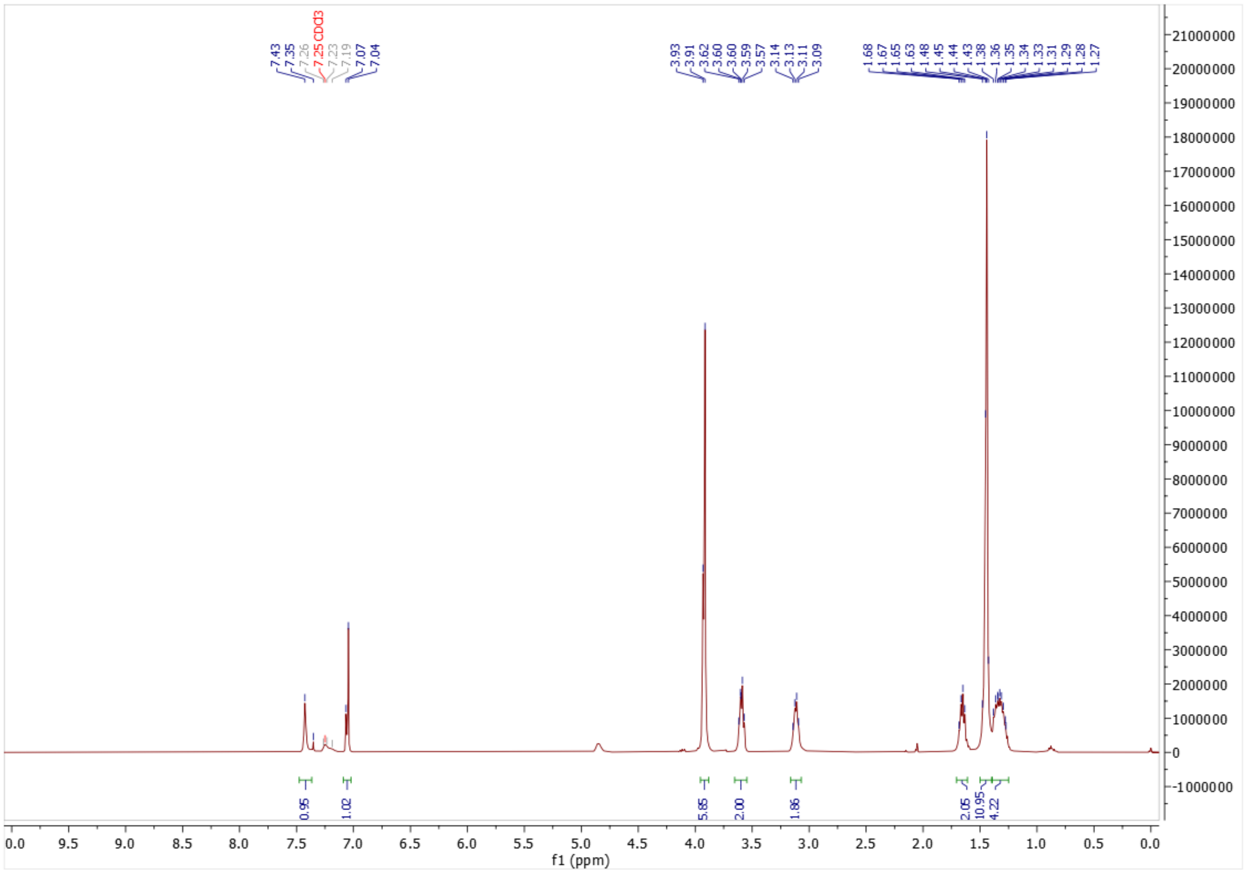

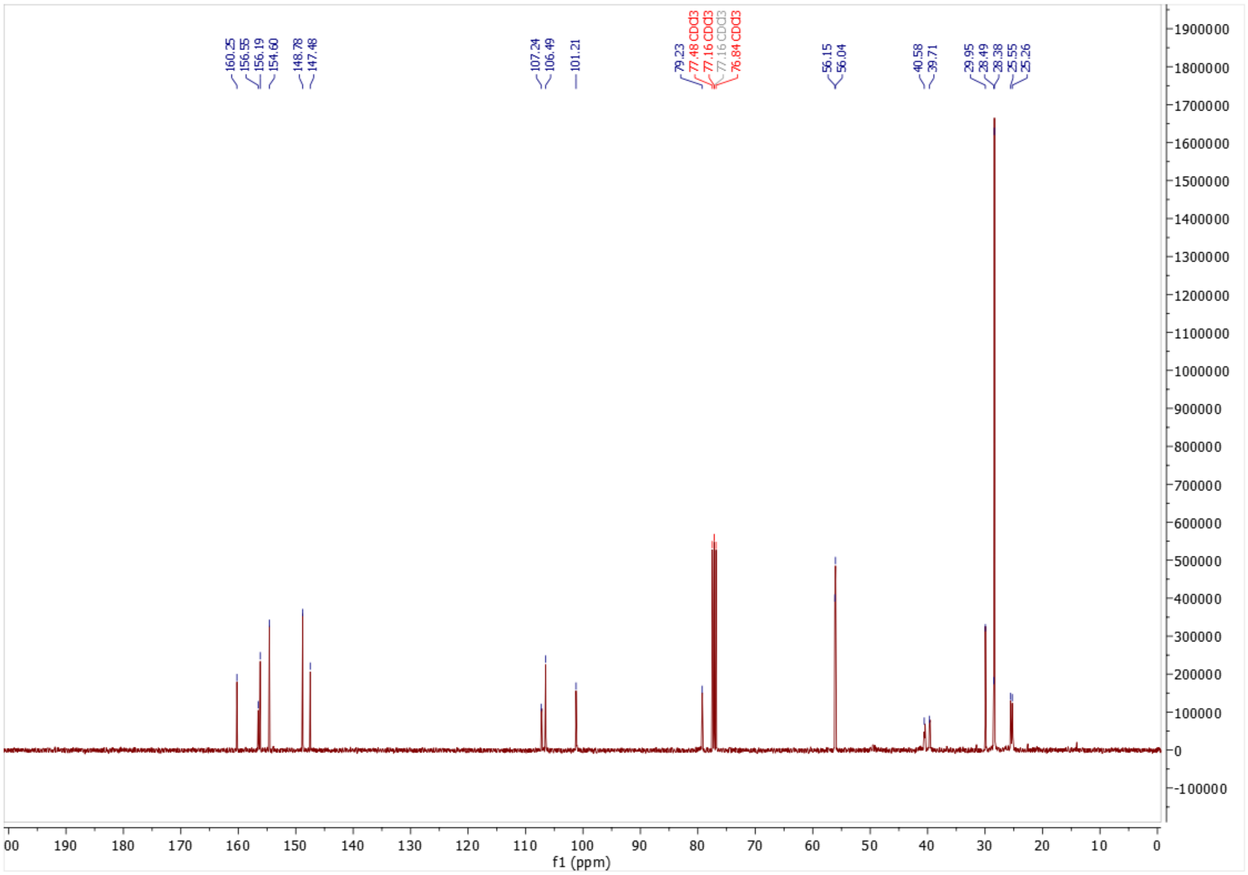

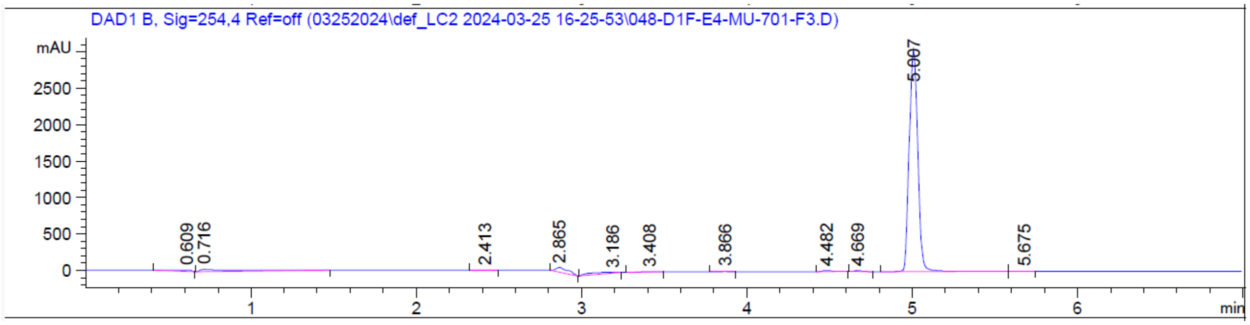

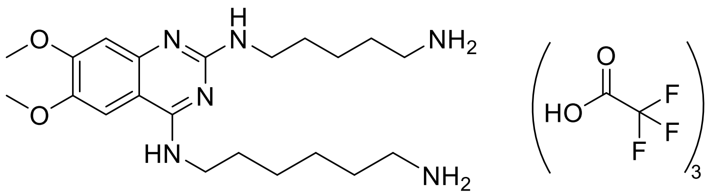

**_N_[4]-(6-aminohexyl)-_N_[2]-(5-aminopentyl)-6,7-dimethoxyquinazoline-2,4-diamine • 3 TFA salts (Intermediate 6)**

In a microwave vial containing **Intermediate 5** (215 mg, 1.0 eq.) in 0.61 mL of IPA, *N*-Boc-1.5-diaminohexane (123 µL, 1.2 eq.) and DIPEA (171 µL, 2.0 eq.) were added. The mixture was heated at 160 °C for 2 h under microwave irradiation. The solvent was then removed under reduced pressure. The crude mixture was dissolved in 3.7 mL of DCM and TFA (378 µL, 10 eq.) was added. The mixture was stirred at room temperature overnight. The solvent was then removed under reduced pressure. Flash chromatography (5 to 100% MeOH in 0.1% TFA in $H_2O$) yielded the desired product as a yellow oil (99 mg, 50%) as a TFA salt. $^1$H NMR (400 MHz, MeOD) δ 7.51 (s, 1H), 6.90 (s, 1H), 3.95 (s, 3H), 3.90 (s, 3H), 3.66 (t, *J* = 7.1 Hz, 2H), 3.58 – 3.45 (m, 2H), 3.00 – 2.89 (m, 4H), 1.81 – 1.64 (m, 8H), 1.57 – 1.41 (m, 6H). MS(ES+) m/z 405.3 $[M + H]^+$.

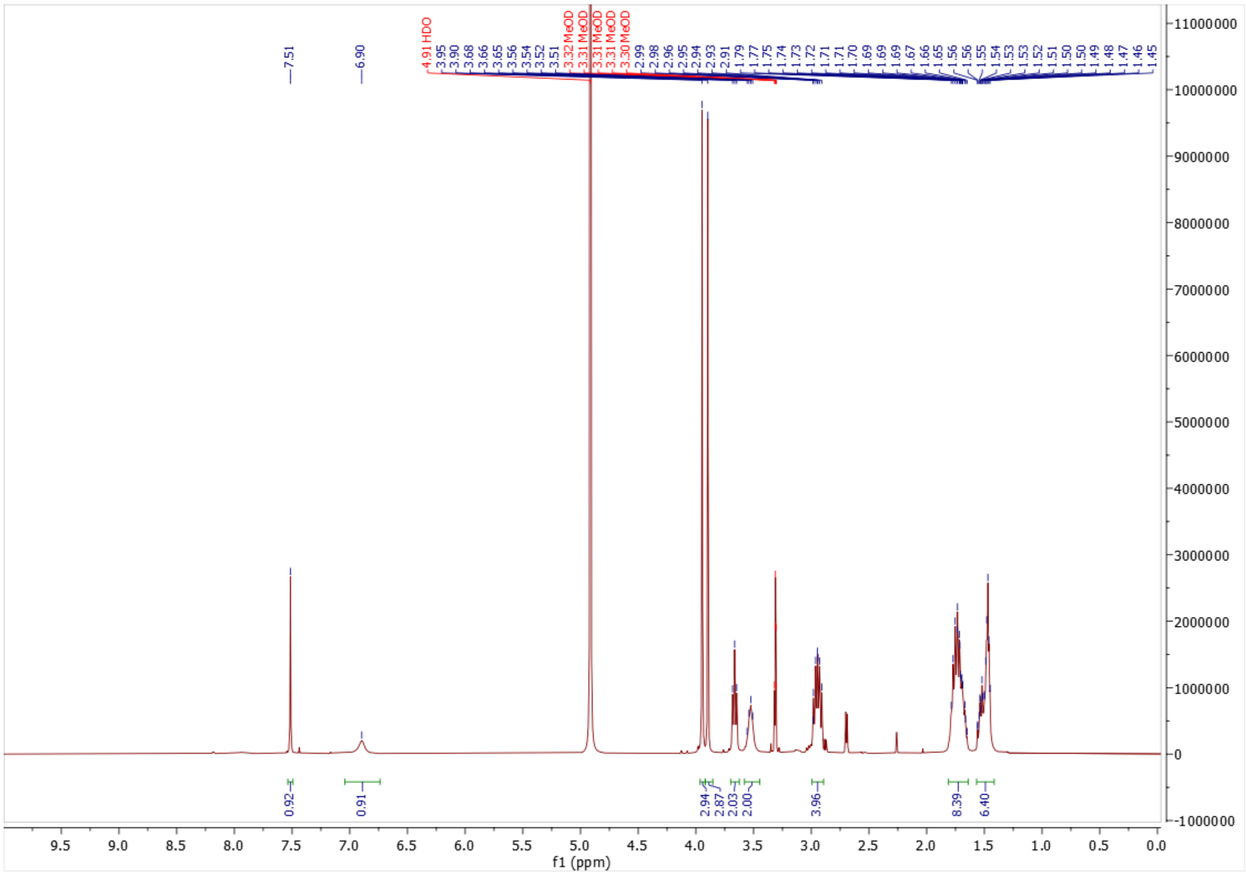

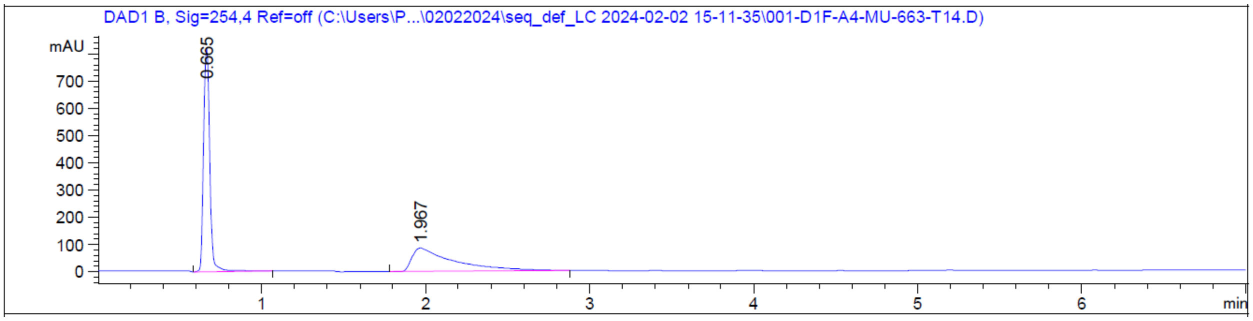

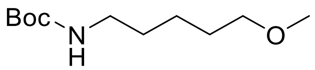

**tert-butyl (5-methoxypentyl)carbamate (Intermediate 7)**

To a round-bottom flask containing *tert*-butyl (5-hydroxypentyl)carbamate (300 mg, 1.0 eq.) in 6.0 mL of anhydrous THF, 60% NaH in mineral oil (70.8 mg, 1.2 eq.) and methyl iodide (111 µL, 1.0 eq.) were added. The mixture was stirred at room temperature overnight under nitrogen. TLC (30% EtOAc in hexane with $KMnO_4$ stain) showed reaction completion. A saturated aqueous solution of $NH_4Cl$ was added to the mixture and extracted with EtOAc. The organic fractions were combined, dried over $MgSO_4$ and the solvent was removed under reduced pressure. Flash chromatography (0 to 50% EtOAc in hexane) yielded **Intermediate 7** as a transparent oil (316 mg, 98%). $^1$H NMR (400 MHz, CDCl3) δ 4.57 (s, 1H), 3.37 – 3.31 (m, 2H), 3.31 – 3.24 (m, 3H), 3.13 – 3.00 (m, 2H), 1.61 – 1.51 (m, 2H), 1.51 – 1.44 (m, 2H), 1.44 – 1.37 (m, 9H), 1.38 – 1.27 (m, 2H).

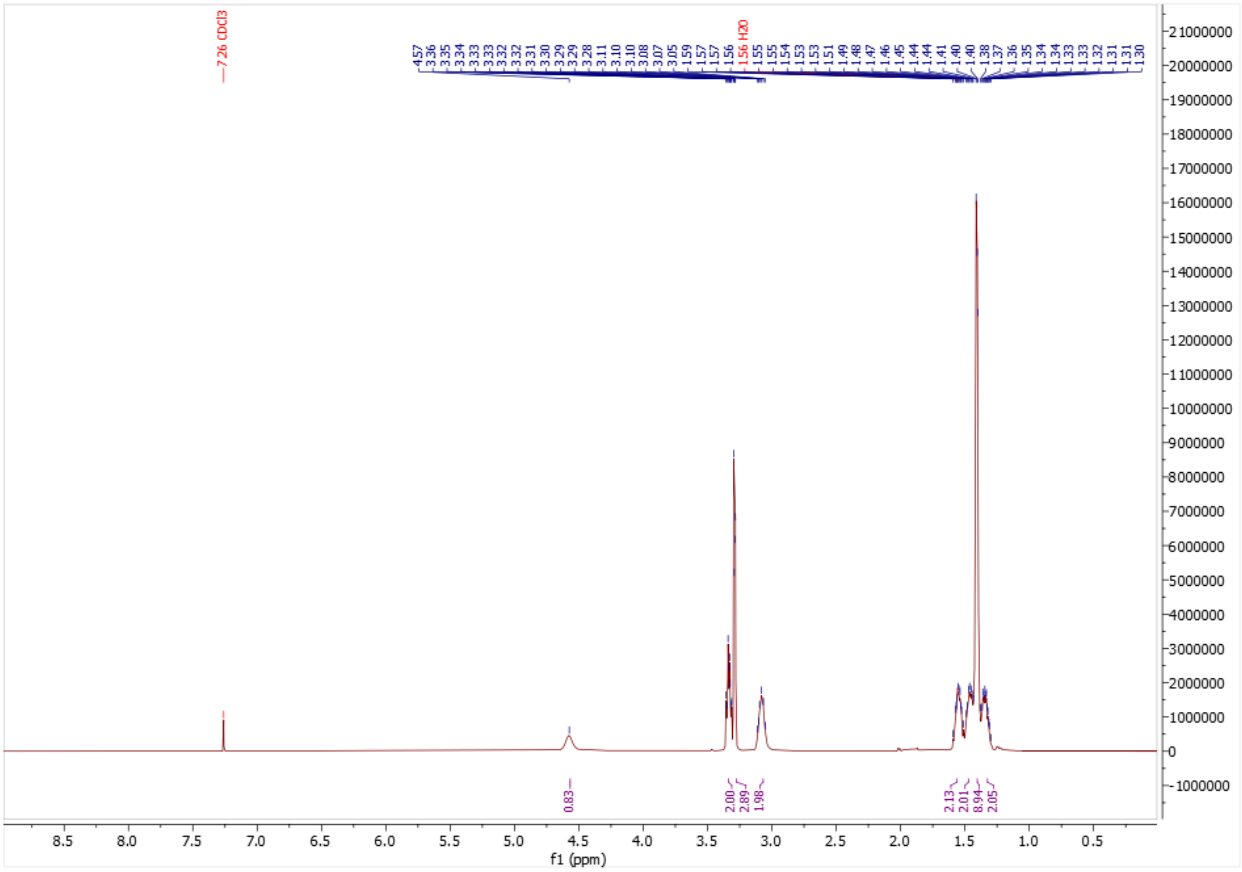

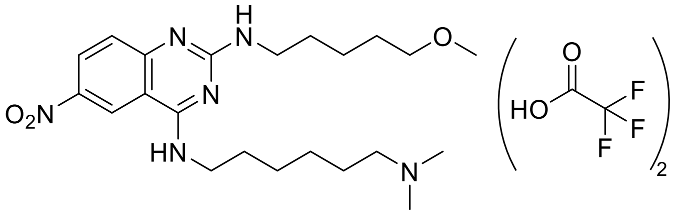

**$N^4$-(6-(dimethylamino)hexyl)-$N^2$-(5-methoxypentyl)-6-nitroquinazoline-2,4-diamine • 2 TFA salts (Intermediate 8)**

In a round-bottom flask, **Intermediate 7** (315.5 mg, 1.0 eq.) was dissolved in 10.0 mL of DCM. TFA (1.1 mL, 10 eq.) was then added. The mixture was stirred at room temperature overnight. The solvent and TFA were then removed under reduced pressure yielding 5-methoxypentan-1-amine as a transparent oil (257 mg, 77%). Then, crude 5-methoxypentan-1-amine (120.0 mg, 1.1 eq.) and **Intermediate 1** (200.0 mg, 1.0 eq.) were dissolved in 1.2 mL of IPA in a microwave vial. DIPEA (164 µL, 2.0 eq.) was then added and the mixture was heated at 160 °C for 2 h under microwave irradiation. The solvent was removed under reduced pressure. The crude mixture was dissolved in 1.2 mL of DCM, and TFA (364 µL, 10 eq.) was added. The mixture was stirred at room temperature overnight. The solvent and TFA were then removed under reduced pressure. The resulting crude was dissolved in 1.2 mL of MeOH before a 37% formaldehyde solution in water (106 µL, 3.0 eq.) and a drop of acetic acid were added. The mixture was stirred at room temperature for 1 h before sodium triacetoxyborohydride (300 mg, 3.0 eq.) was added partwise. The mixture was stirred at room temperature overnight. The solvent was then removed under reduced pressure. Flash chromatography (5 to 60% MeOH in 0.1% TFA in $H_2O$) yielded **Intermediate 8** as a transparent oil (77 mg, 25%). $^1$H NMR (400 MHz, MeOD) δ 9.08 (s, 1H), 8.45 (d, J = 8.8 Hz, 1H), 7.49 (d, J = 9.1 Hz, 1H), 3.67 (t, J = 7.2 Hz, 2H), 3.53 (t, J = 7.1 Hz, 2H), 3.37 (t, J = 6.4 Hz, 2H), 3.27 (s, 3H), 3.09 (t, J = 8.1 Hz, 2H), 2.84 (s, 6H), 1.73 (dh, J = 30.4, 7.0 Hz, 6H), 1.59 (p, J = 6.8 Hz, 2H), 1.45 (dh, J = 15.0, 7.1 Hz, 6H). MS(ES+) m/z 433.4 $[M + H]^+$.

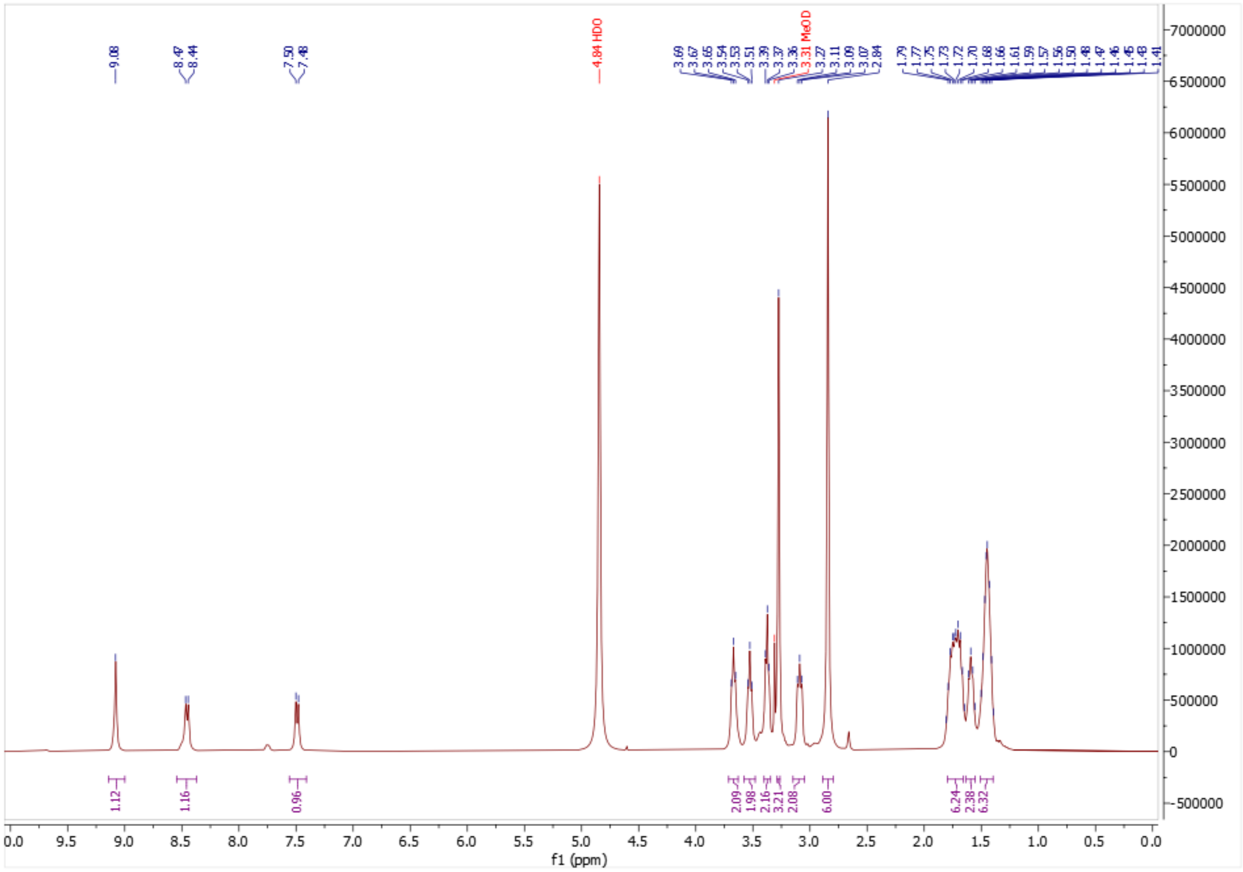

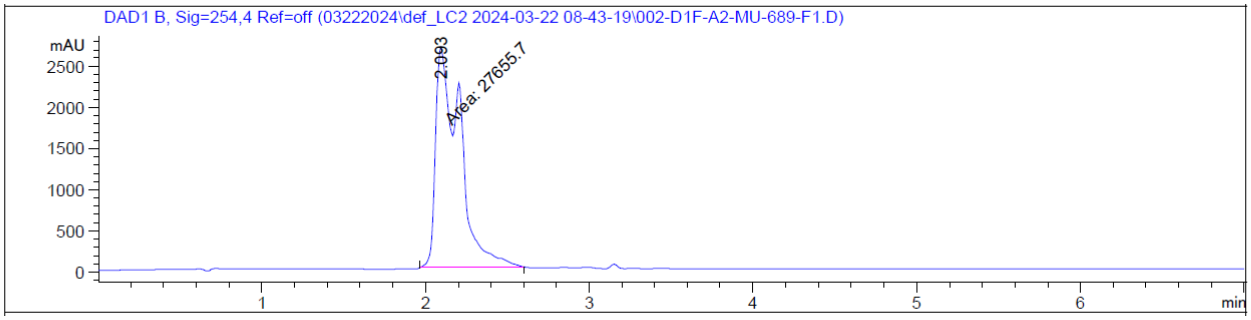

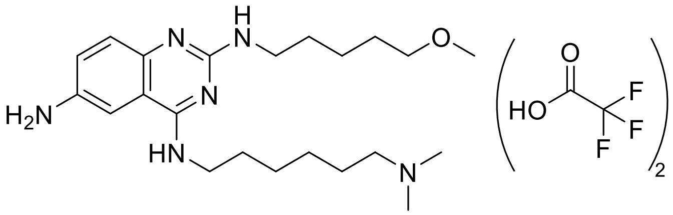

**$N^4$-(6-(dimethylamino)hexyl)-$N^2$-(5-methoxypentyl)quinazoline-2,4,6-triamine • 2 TFA salts (Intermediate 9)**

In a round-bottom flask containing **Intermediate 8** (77 mg, 1.0 eq.) in 0.4 mL of MeOH, 10 % palladium on carbon (19 mg, 0.1 eq.) was added. A $H_2$ balloon was fitted on the

flask after removing the air. The mixture was stirred at room temperature overnight. After reaction completion, the mixture was filtered over celite using MeOH for rinsing and then the solvent was removed under reduced pressure. Flash chromatography (5 to 100% MeOH in 0.1% TFA in $H_2O$) yielded the desired compound as a brown oil (50 mg, 45%) as a TFA salt. $^1$H NMR (400 MHz, MeOD) δ 7.67 (s, 1H), 7.45 (dd, *J* = 8.6, 2.3 Hz, 1H), 7.40 (s, 1H), 3.70 (t, *J* = 7.2 Hz, 2H), 3.51 (s, 2H), 3.41 (t, *J* = 6.4 Hz, 2H), 3.31 (s, 3H), 3.15 – 3.07 (m, 2H), 2.87 (s, 6H), 1.84 – 1.67 (m, 6H), 1.66 – 1.58 (m, 2H), 1.55 – 1.39 (m, 6H). MS(ES+) m/z 403.4 [M + H]$^+$.

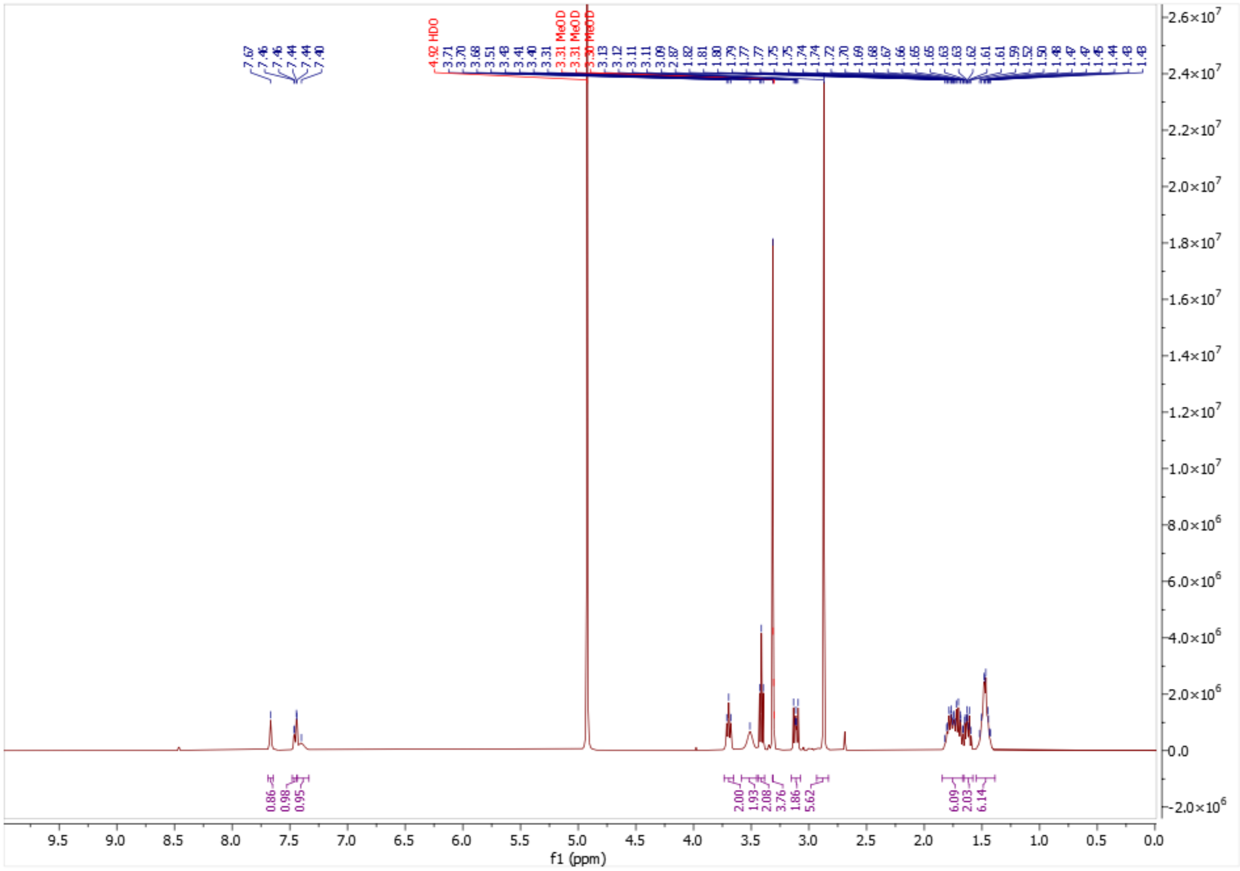

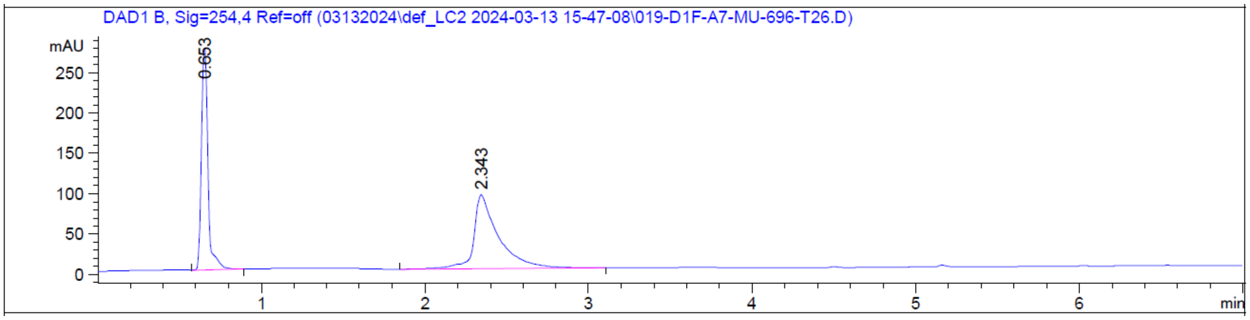

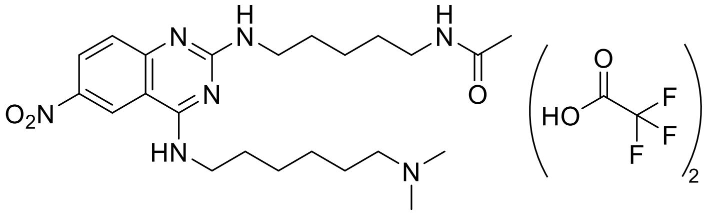

**N-(5-((4-((6-(dimethylamino)hexyl)amino)-6-nitroquinazolin-2-yl)amino)pentyl)acetamide • 2 TFA salts (Intermediate 10)**

In a round-bottom flask, **Intermediate 1** (200 mg, 1.0 eq.) and *N*-(5-aminopentyl)acetamide (134 mg, 1.1 eq.) were dissolved in 1.2 mL of IPA in a microwave vial. DIPEA (164 µL, 2.0 eq.) was then added and the mixture was heated at 160 °C for 2 h under microwave irradiation. The solvent was removed under reduced pressure. The crude mixture was dissolved in 1.2 mL of DCM, and TFA (364 µL, 10 eq.) was added. The mixture was stirred at room temperature overnight. The solvent and TFA were then removed under reduced pressure. The resulting crude was dissolved in 1.2 mL of MeOH before a 37% formaldehyde solution in water (106 µL, 3.0 eq.) and a drop of acetic acid were added. The mixture was stirred at room temperature for 1 h before sodium triacetoxyborohydride (300 mg, 3.0 eq.) was added partwise. The mixture was stirred at room temperature overnight. The solvent was then removed under reduced pressure. Flash chromatography (5 to 100% MeOH in 0.1% TFA in $H_2O$) yielded **Intermediate 10** as a light yellow oil (150 mg, 46%). $^1H$ NMR (400 MHz, MeOD) δ 9.13 (s, 1H), 8.50 (d, *J* = 9.3 Hz, 1H), 7.54 (d, *J* = 9.1 Hz, 1H), 3.71 (t, *J* = 7.4 Hz, 2H), 3.57 (t, *J* = 7.2 Hz, 2H), 3.22 – 3.06 (m, 4H), 2.88 (s, 6H), 1.92 (s, 3H), 1.85 – 1.68 (m, 6H), 1.62 – 1.26 (m, 8H).MS(ES+) m/z 460.3 $[M + H]^+$.

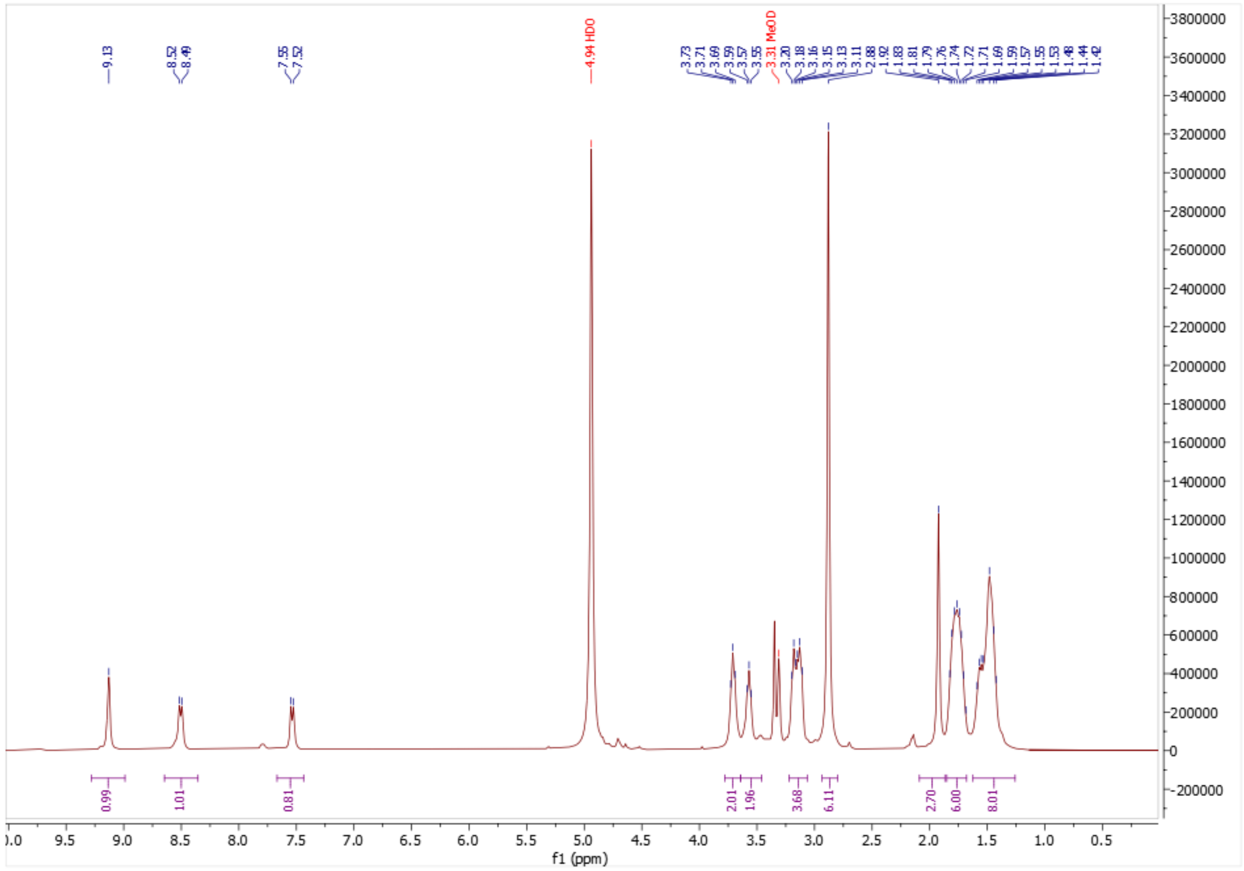

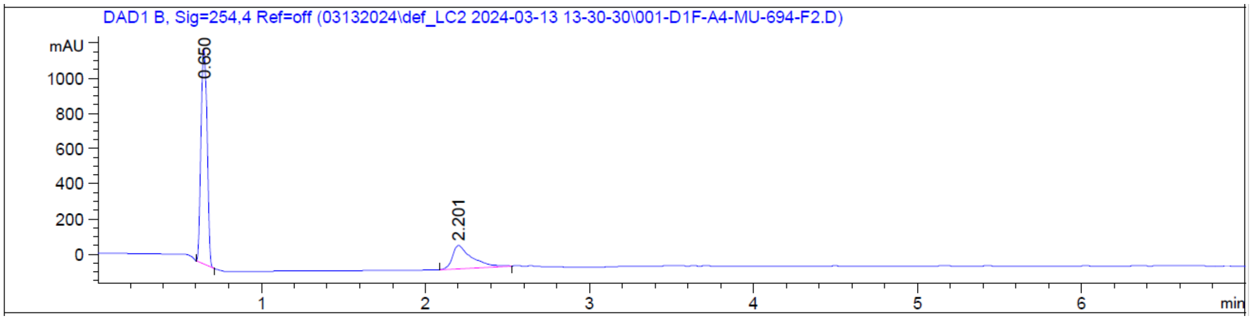

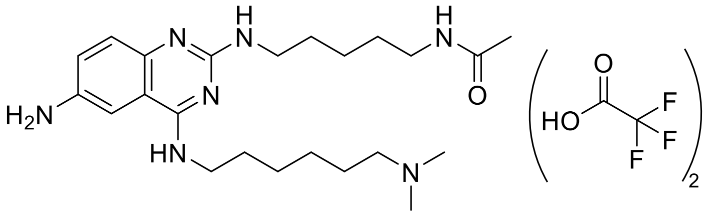

***N*-(5-((6-amino-4-((6-(dimethylamino)hexyl)amino)quinazolin-2-yl)amino)pentyl)acetamide • 2 TFA salts (Intermediate 11)**

In a round-bottom flask containing **Intermediate 10** (150 mg, 1.0 eq.) in 0.8 mL of MeOH, 10 % palladium on carbon (35 mg, 0.1 eq.) was added. A $H_2$ balloon was fitted on the

flask after removing the air. The mixture was stirred at room temperature overnight. After reaction completion, the mixture was filtered over celite using MeOH for rinsing and then the solvent was removed under reduced pressure. Flash chromatography (5 to 100% MeOH in 0.1% TFA in $H_2O$) yielded the desired compound as a light brown oil (78 mg, 36%) as a TFA salt. $^1$H NMR (400 MHz, MeOD) δ 7.79 (s, 1H), 7.52 (dd, $J$ = 8.8, 2.3 Hz, 1H), 7.44 (s, 1H), 3.70 (t, $J$ = 7.2 Hz, 2H), 3.52 (s, 2H), 3.18 (t, $J$ = 7.0 Hz, 2H), 3.15 – 3.08 (m, 2H), 2.87 (s, 6H), 1.92 (s, 3H), 1.84 – 1.65 (m, 6H), 1.62 – 1.37 (m, 8H). MS(ES+) m/z 430.4 [M + H]$^+$.

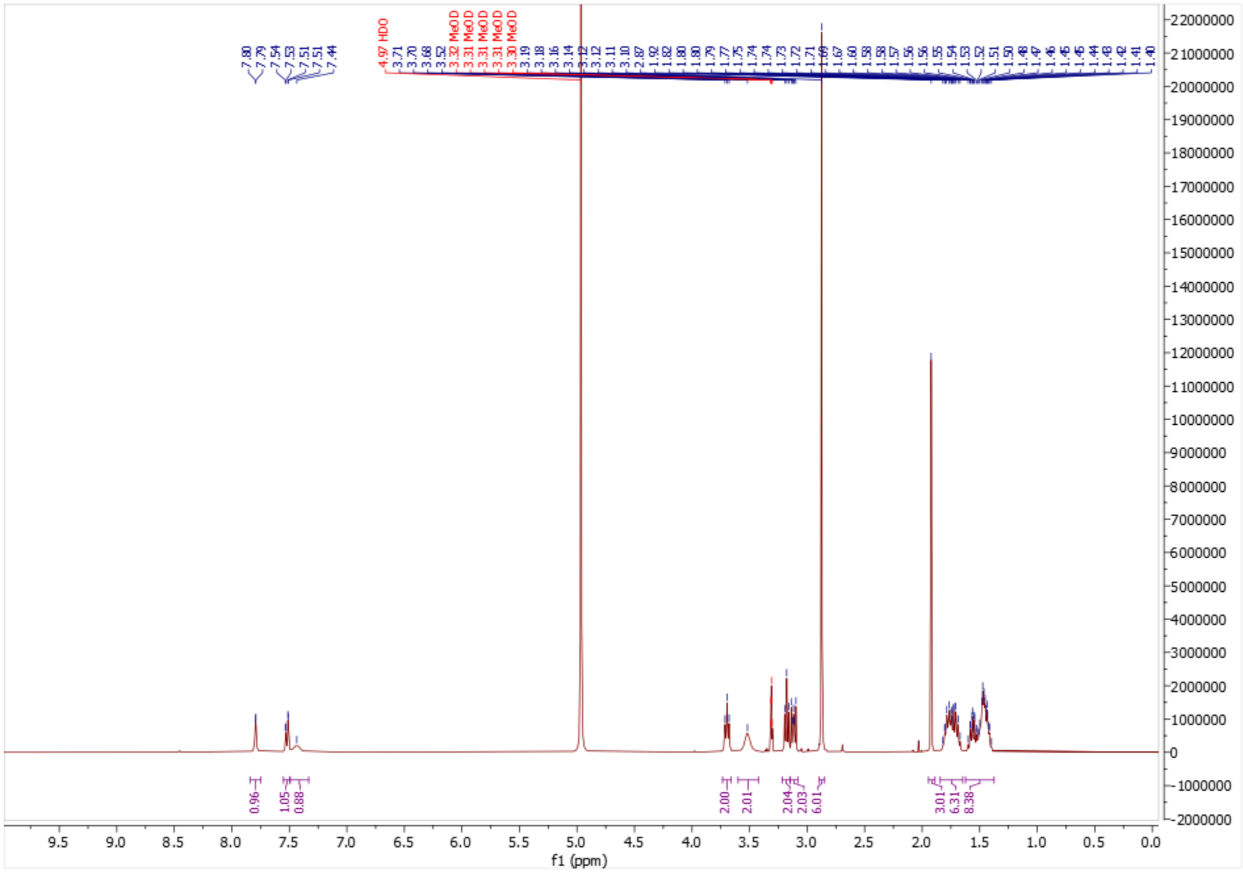

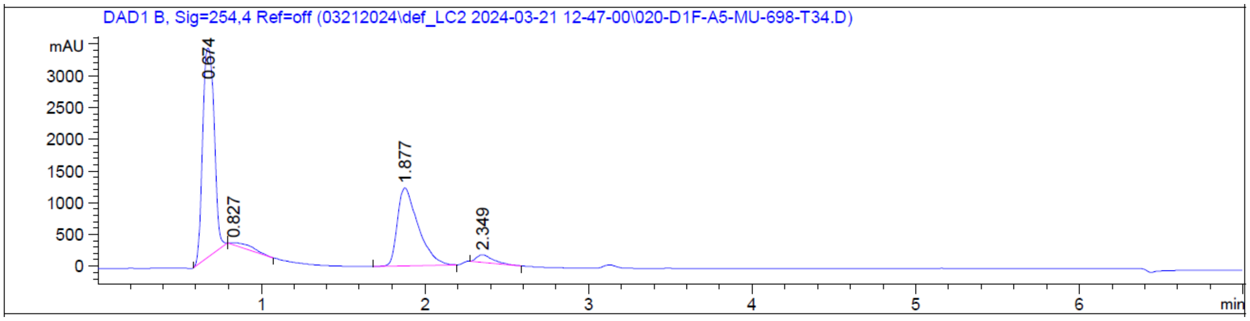

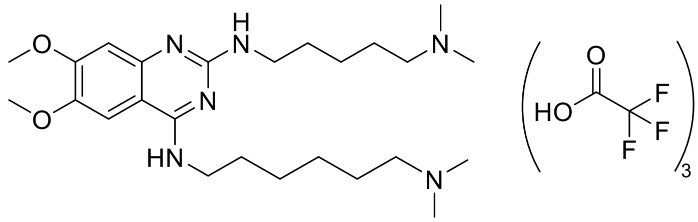

**$N^4$-(6-(dimethylamino)hexyl)-$N^2$-(5-(dimethylamino)pentyl)-6,7-dimethoxyquinazoline-2,4-diamine • 3 TFA salts (UNC6535)**

In a round-bottom flask containing **Intermediate 6** (99 mg, 1.0 eq.) in 1.2 mL of MeOH, a 37% formaldehyde solution in $H_2O$ (110 µL, 6.0 eq.) was added. The mixture was stirred at room temperature for 1 h before sodium triacetoxyborohydride (416 mg, 8.0 eq.) was added partwise. The mixture was stirred at room temperature overnight. Water and MeOH were then added to the mixture before flash chromatography loading. Flash chromatography (5 to 100% MeOH in 0.1% TFA in $H_2O$) followed by semi-prep HPLC (5 to 50% MeOH in 0.05% TFA in $H_2O$) and strong cation exchange desalting (MeOH followed by 10% NH3 in MeOH) yielded the desired product as a yellow oil (4 mg, 3%) as a TFA salt. $^1$H NMR (400 MHz, MeOD) δ 7.36 (s, 1H), 6.83 (s, 1H), 3.91 (s, 3H), 3.89 (s, 3H), 3.59 (t, *J* = 7.2 Hz, 2H), 3.44 (t, *J* = 7.0 Hz, 2H), 2.43 – 2.33 (m, 4H), 2.29 (s, 6H), 2.26 (s, 6H), 1.78 – 1.63 (m, 4H), 1.61 – 1.49 (m, 4H), 1.48 – 1.36 (m, 6H). $^{13}$C NMR (176 MHz, MeOD) δ 161.17, 159.77, 156.09, 147.04, 104.91, 104.25, 104.14, 100.81, 60.61, 56.83, 56.31, 45.33, 45.30, 42.25, 42.03, 30.89, 30.27, 28.35, 28.17, 28.10, 27.99, 25.97. MS(ES+) m/z 461.4 $[M + H]^+$.

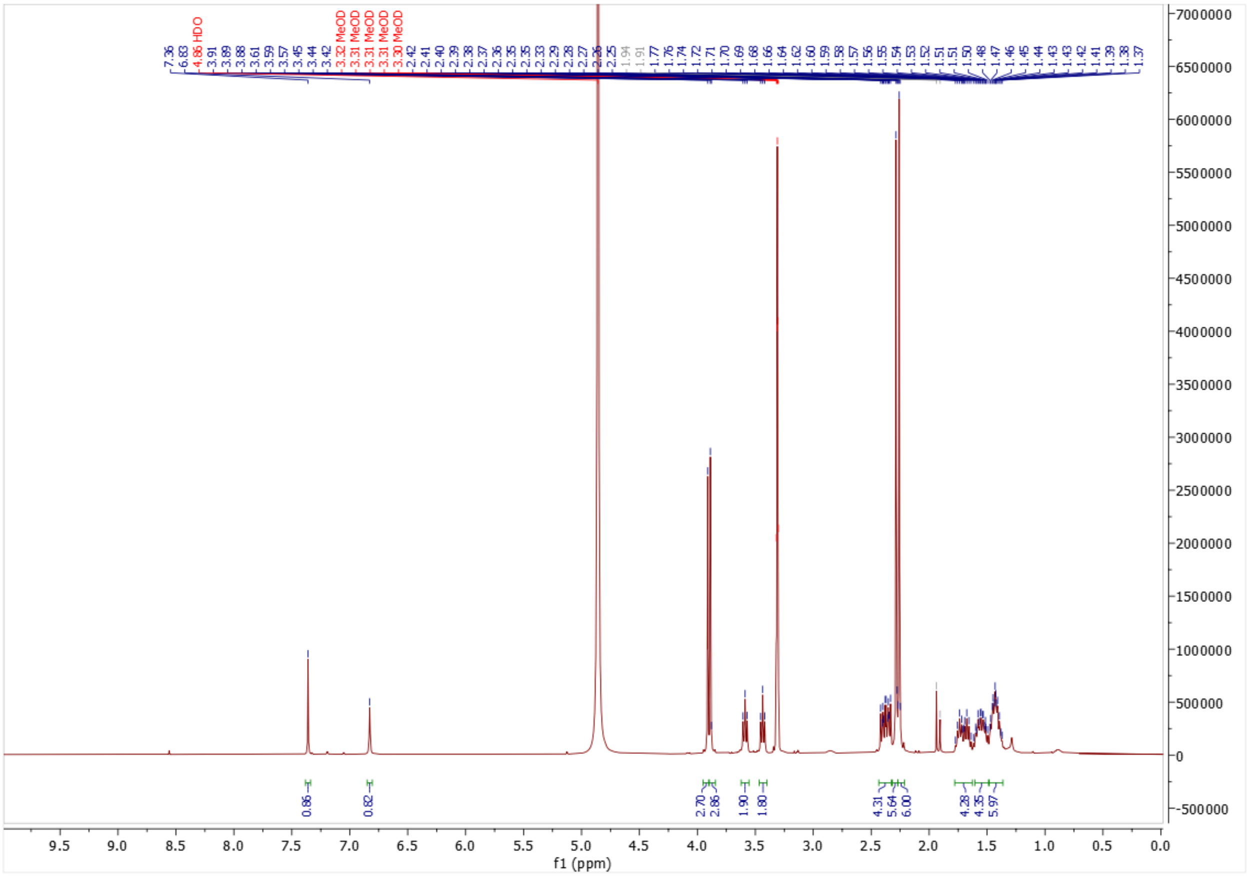

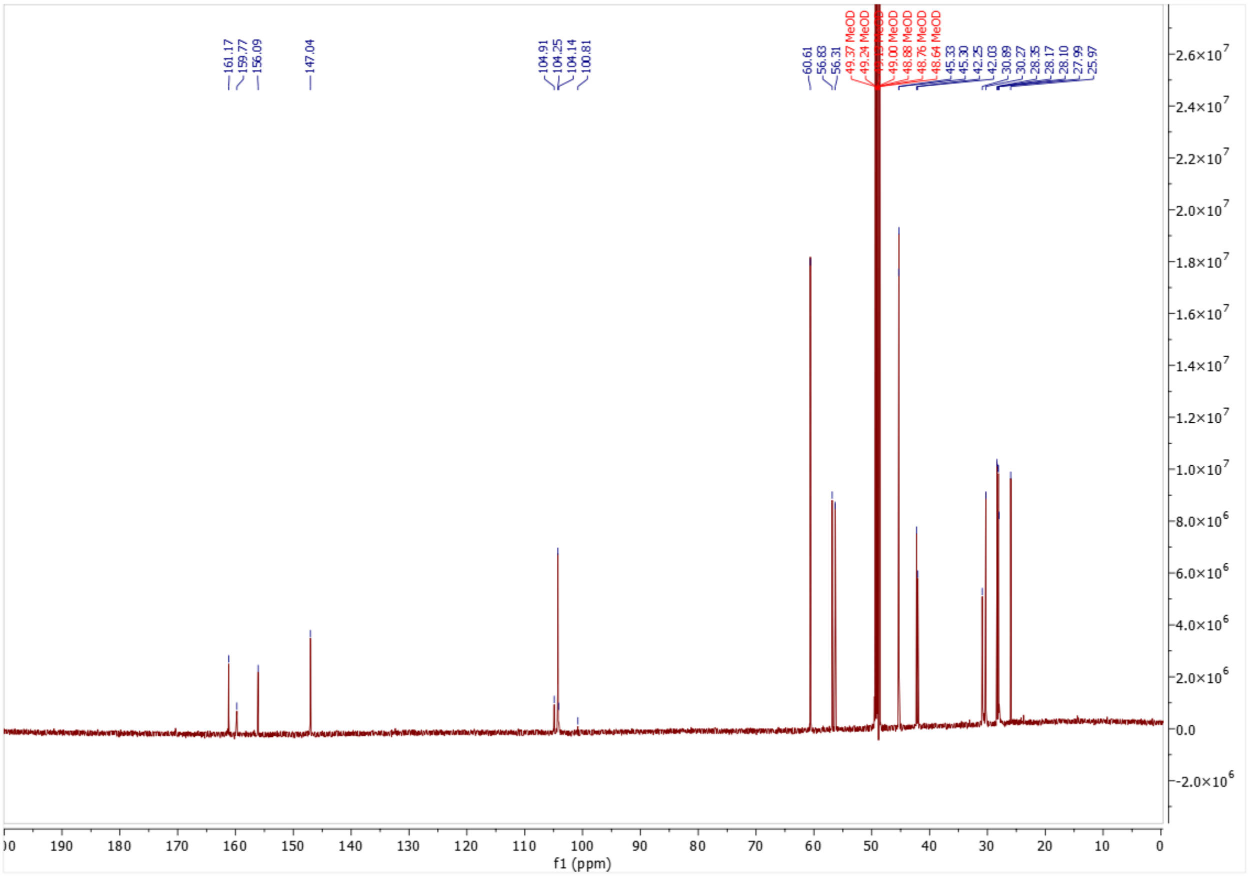
161.17
159.77
156.09
147.04
104.91
104.25
104.14
100.81
60.61
56.83
56.31
49.37 MeOD
49.24 MeOD
49.00 MeOD
48.88 MeOD
48.76 MeOD
48.64 MeOD
45.33
45.30
42.25
42.03
30.89
30.27
28.35
28.17
28.10
27.99
25.97
f1 (ppm)

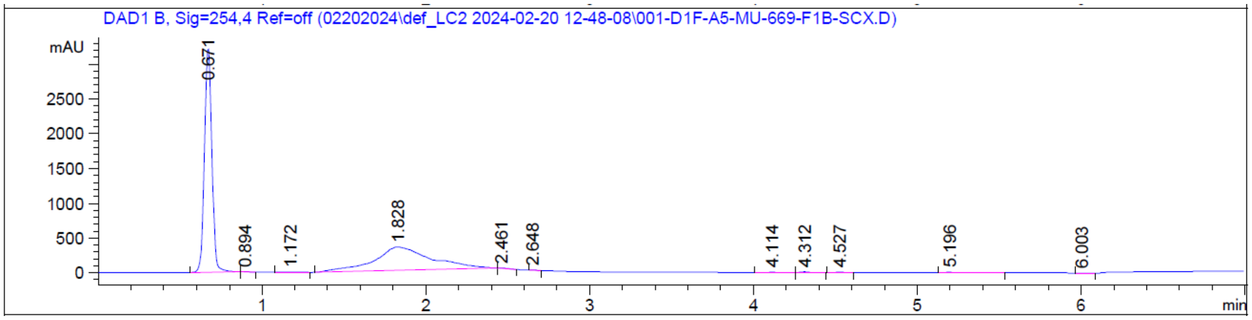
DAD1 B, Sig=254,4 Ref=off (02202024\def_LC2 2024-02-20 12-48-08\001-D1F-A5-MU-669-F1B-SCX.D)
mAU
0.671
0.894
1.172
1.828
2.461
2.648
4.114
4.312
4.527
5.196
6.003
min

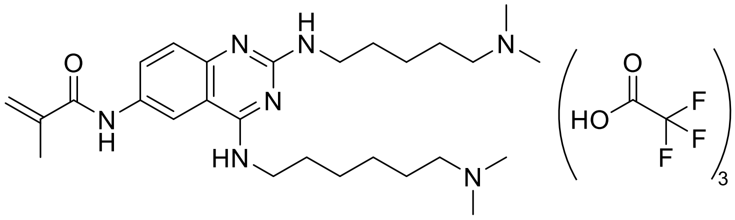

***N*-(4-((6-(dimethylamino)hexyl)amino)-2-((5-(dimethylamino)pentyl)amino)quinazolin-6-yl)methacrylamide • 3 TFA salts (UNC10014 (1))**

**UNC10014** was obtained by following General Procedure 1 using **Intermediate 4** and methacryloyl chloride. Flash chromatography (10 to 50% MeOH in 0.1% TFA in $H_2O$) yielded the desired product as a yellow oil (1 mg, 2%) as a TFA salt. $^1$H NMR (400 MHz, MeOD) δ 8.47 (d, *J* = 2.2 Hz, 1H), 7.77 (dd, *J* = 8.9, 2.2 Hz, 1H), 7.42 (s, 1H), 5.92 – 5.82 (m, 1H), 5.58 (td, *J* = 1.7, 0.9 Hz, 1H), 3.70 (t, *J* = 6.3 Hz, 2H), 3.58 (s, 2H), 3.17 – 3.06 (m, 4H), 2.88 (s, 6H), 2.87 (s, 6H), 2.06 – 2.04 (m, 2H), 1.85 – 1.69 (m, 8H), 1.55 – 1.41 (m, 6H). MS(ES+) m/z 484.4 [M + H]$^+$.

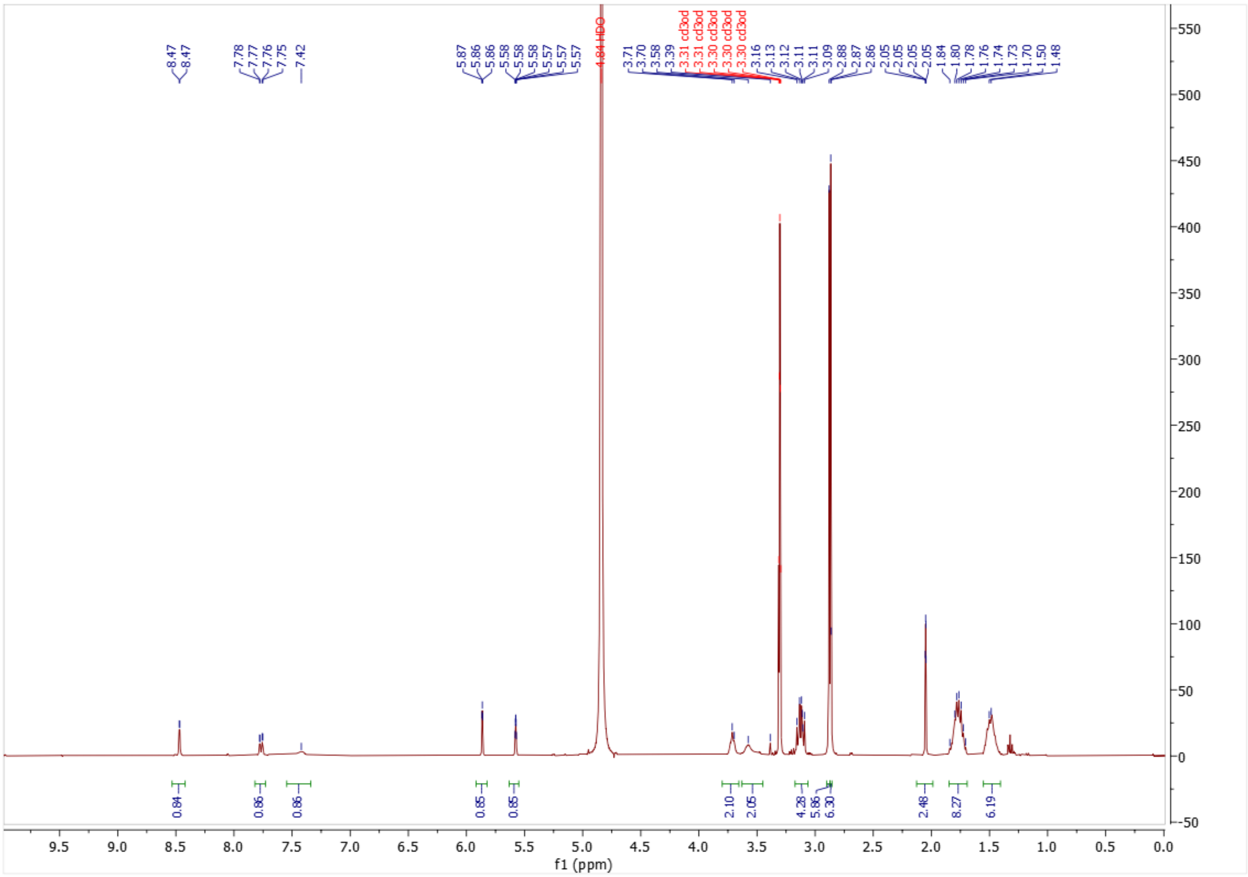

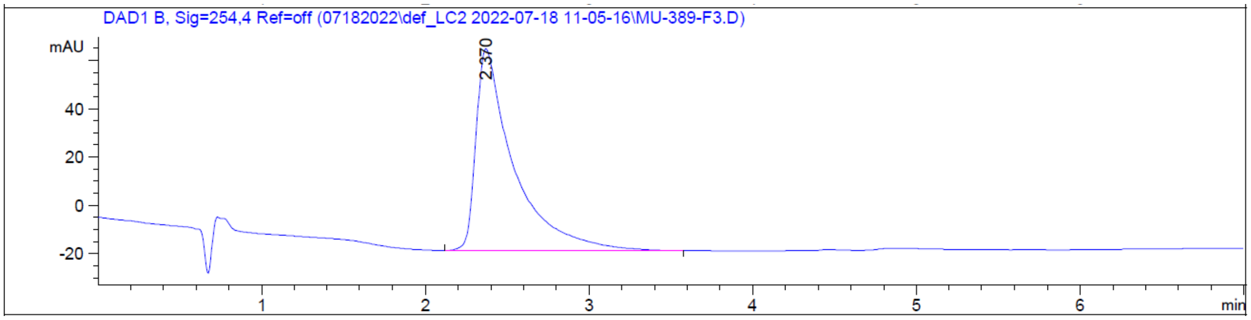

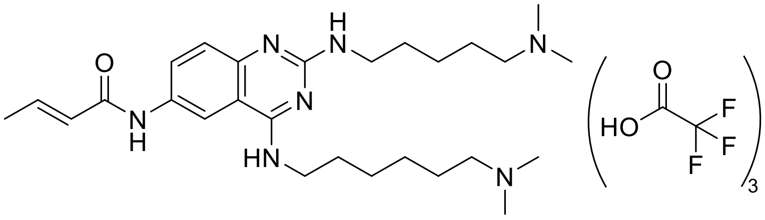

**(*E*)-*N*-(4-((6-(dimethylamino)hexyl)amino)-2-((5-(dimethylamino)pentyl)amino)quinazolin-6-yl)but-2-enamide • 3 TFA salts (UNC10016 (2))**

**UNC10016** was obtained by following General Procedure 1 using **Intermediate 4** and (*E*)-but-2-enoyl chloride. Flash chromatography (10 to 50% MeOH in 0.1% TFA in $H_2O$) followed by semi-prep HPLC purification (5% to 50% MeOH in 0.1% TFA in $H_2O$) yielded the desired product as a yellow solid (7.0 mg, 20%) as a TFA salt. $^1H$ NMR (400 MHz, MeOD) δ 8.53 (d, *J* = 2.2 Hz, 1H), 7.69 (dd, *J* = 9.0, 2.2 Hz, 1H), 7.42 (s, 1H), 7.08 – 6.95 (m, 1H), 6.15 (dq, *J* = 15.2, 1.5 Hz, 1H), 3.72 (t, *J* = 6.7 Hz, 2H), 3.58 (m, 2H), 3.19 – 3.08 (m, 4H), 2.88 (s, 6H), 2.87 (s, 6H), 1.95 (dd, *J* = 6.9, 1.7 Hz, 3H), 1.84 – 1.69 (m, 8H), 1.56 – 1.42 (m, 6H). $^{13}C$ NMR (176 MHz, MeOD) δ 166.70, 162.78 (q, TFA), 161.72, 154.51, 143.27, 136.99, 136.68, 129.35, 126.05, 120.53 (TFA), 118.86 (TFA), 118.41, 117.20 (TFA), 115.51 (TFA), 115.08, 111.15, 58.93, 58.85, 43.40, 43.38, 42.89, 41.96, 29.73, 29.47, 27.70, 27.29, 25.62, 25.33, 24.76, 18.00. MS(ES+) m/z 484.3 $[M + H]^+$.

Note: IPA in NMR

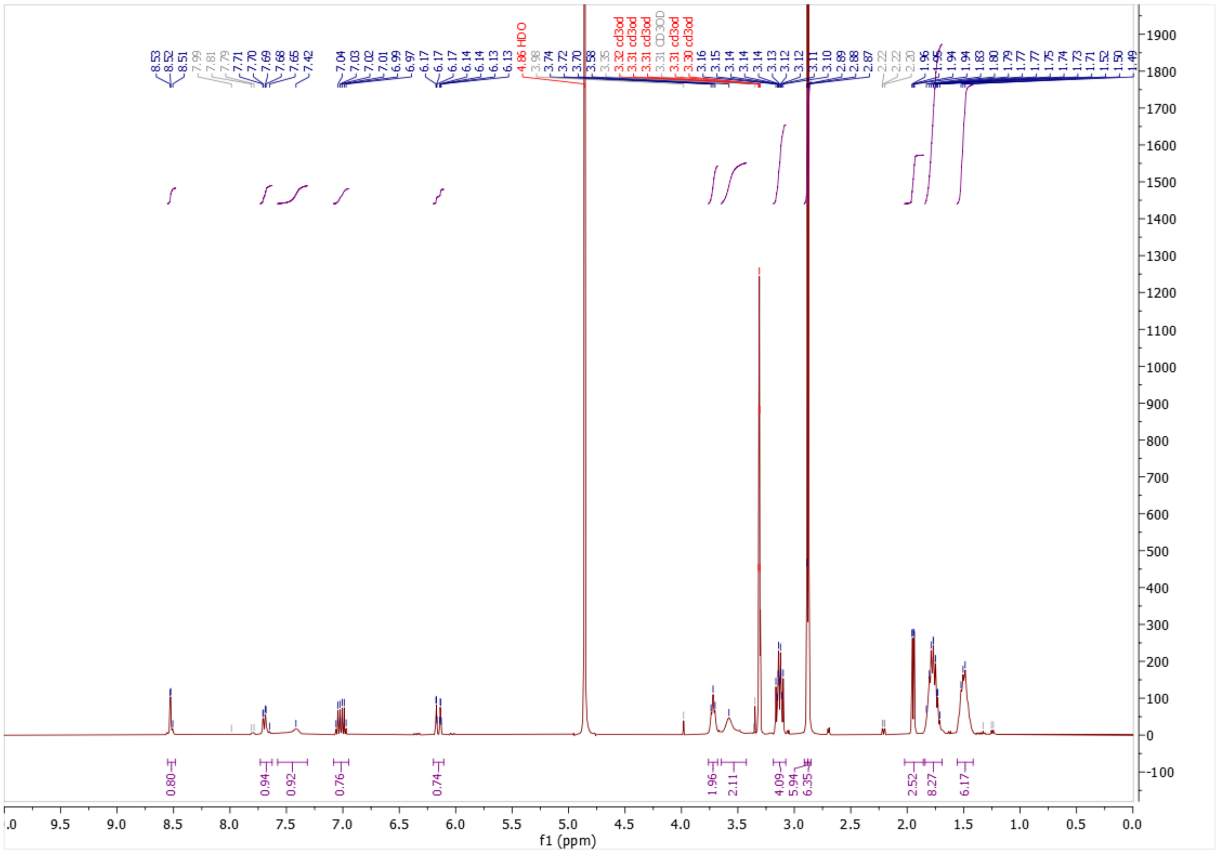
f1 (ppm)

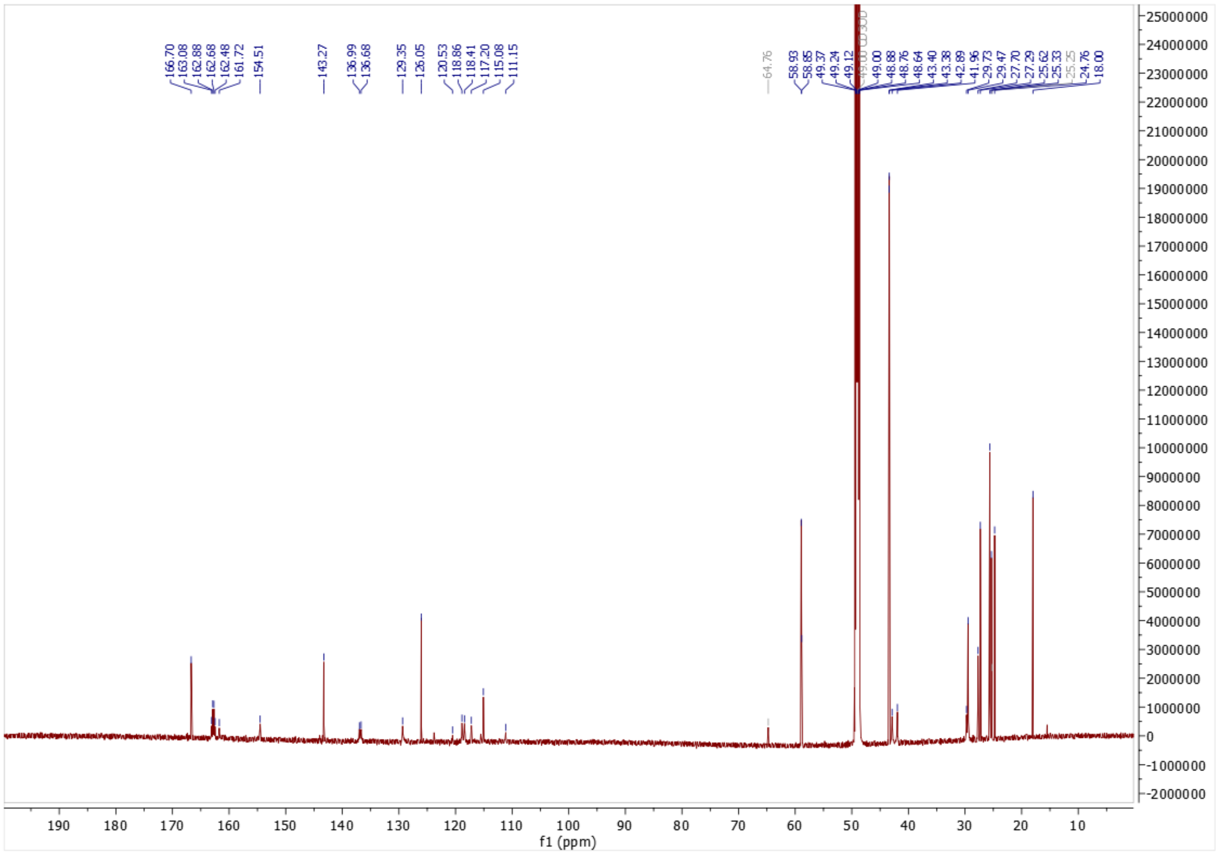

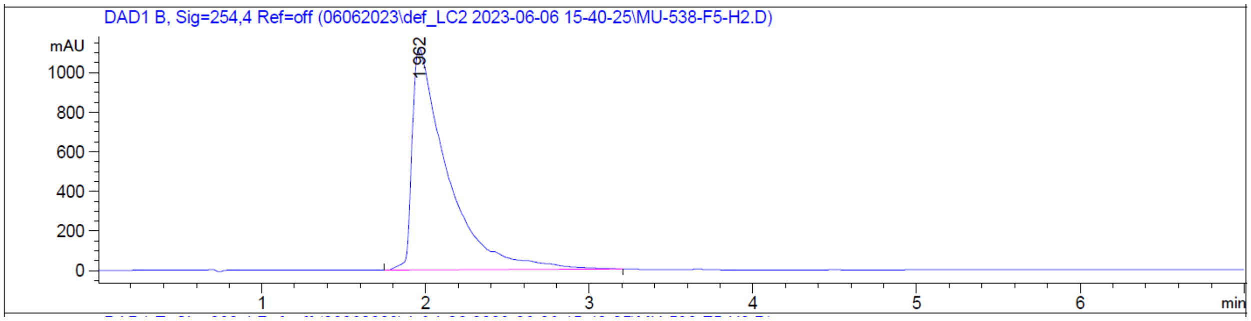
DAD1 B, Sig=254,4 Ref=off (06062023\def_LC2 2023-06-06 15-40-25\MU-538-F5-H2.D)

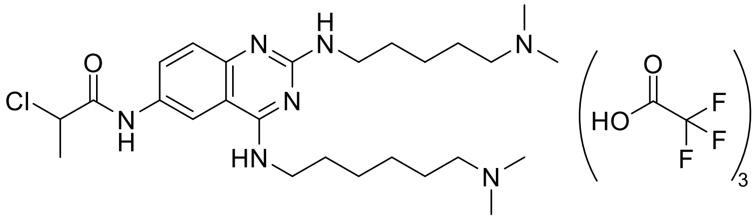

**2-chloro-*N*-(4-((6-(dimethylamino)hexyl)amino)-2-((5-(dimethylamino)pentyl)amino)quinazolin-6-yl)propenamide • 3 TFA salts (UNC9846 (3))**

**UNC9846** was obtained by following General Procedure 1 using **Intermediate 4** and 2-chloropropanoyl chloride. Flash chromatography (10 to 50% MeOH in 0.1% TFA in $H_2O$) yielded the desired product as a light yellow oil (9 mg, 24%) as a TFA salt. $^1$H NMR (400 MHz, MeOD) δ 8.47 (d, *J* = 2.2 Hz, 1H), 7.72 (dd, *J* = 8.9, 2.2 Hz, 1H), 7.43 (s, 1H), 3.71 (t, *J* = 6.6 Hz, 2H), 3.58 (s, 2H), 3.19 – 3.08 (m, 4H), 2.88 (s, 6H), 2.87 (s, 6H), 1.85 – 1.73 (m, 8H), 1.72 (d, *J* = 6.7 Hz, 3H), 1.57 – 1.40 (m, 6H). $^{13}$C NMR (176 MHz, MeOD) δ 170.52, 162.96 (q, TFA), 161.67, 154.59, 137.50, 135.89, 129.68, 118.97 (TFA), 118.50, 117.33 (TFA), 115.85, 111.18, 58.92, 58.84, 55.47, 43.40, 43.37, 42.89, 41.93, 29.70, 29.41, 27.67, 27.26, 25.60, 25.31, 24.74, 21.66. MS(ES+) m/z 506.3 [M + H]$^+$.

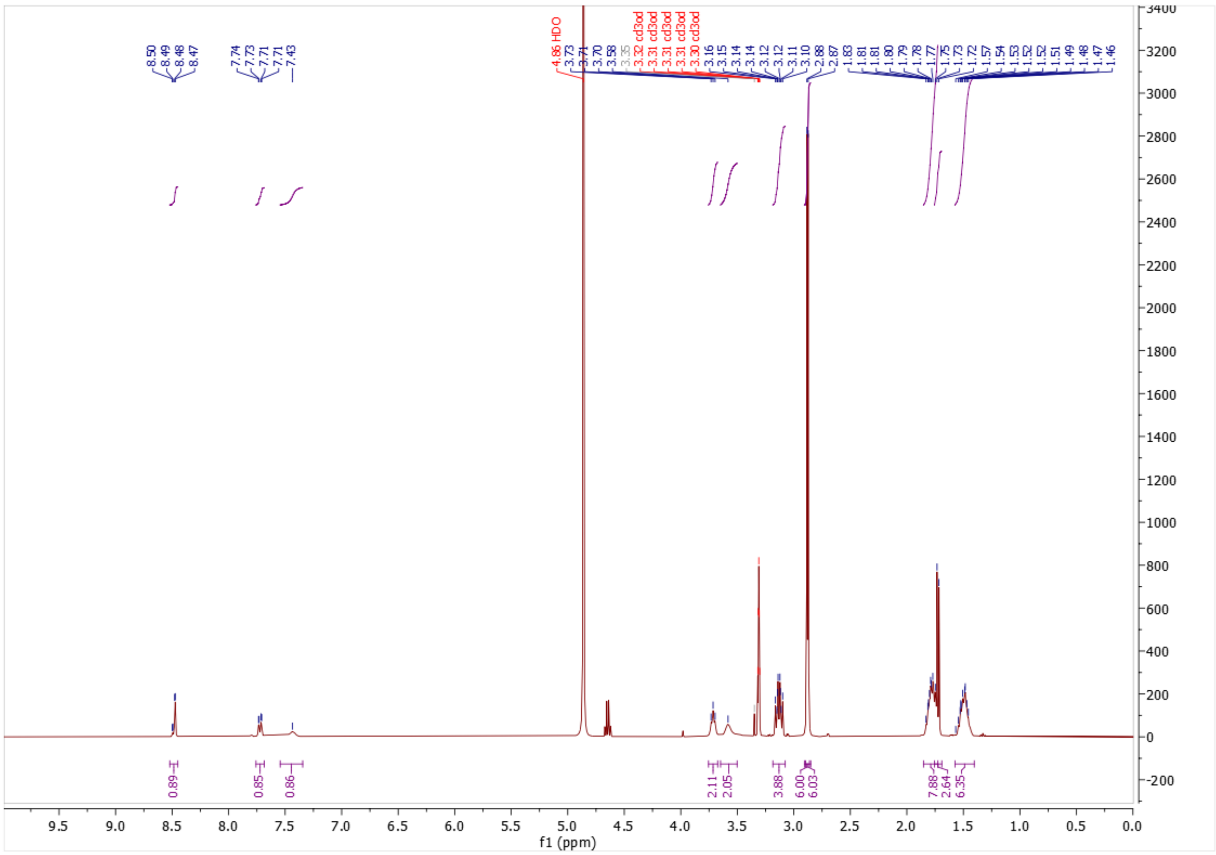

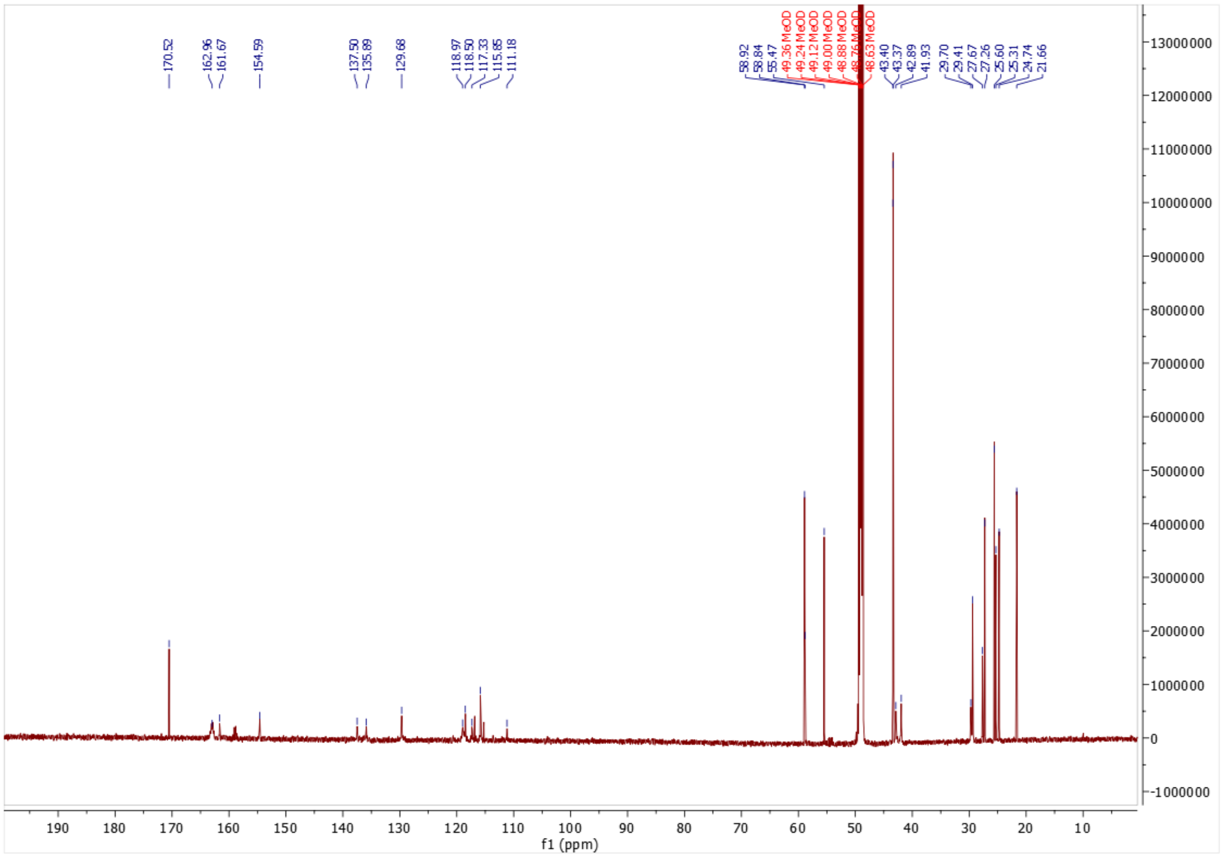
170.52
162.96
161.67
154.59
137.50
135.89
129.68
118.97
118.50
117.33
115.85
111.18
58.92
58.84
55.47
49.36 MeOD
49.24 MeOD
49.12 MeOD
49.00 MeOD
48.88 MeOD
48.76 MeOD
48.63 MeOD
43.40
43.37
42.89
41.93
29.70
29.41
27.67
27.26
25.60
25.31
24.74
21.66
f1 (ppm)

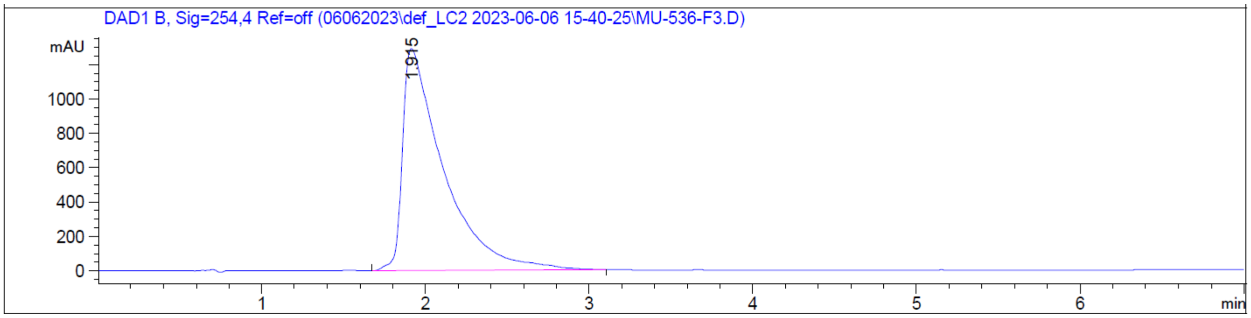
DAD1 B, Sig=254,4 Ref=off (06062023\def_LC2 2023-06-06 15-40-25\MU-536-F3.D)
mAU
1.915
min

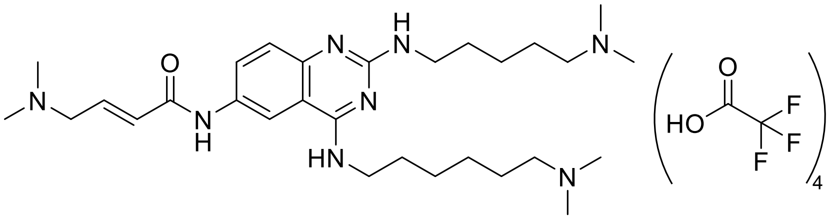

**(*E*)-4-(dimethylamino)-*N*-(4-((6-(dimethylamino)hexyl)amino)-2-((5-(dimethylamino)pentyl)amino)quinazolin-6-yl)but-2-enamide • 4 TFA salts (UNC10013 (4))**

**UNC10013** was obtained by following General Procedure 2 using **Intermediate 4** and (*E*)-4-(dimethylamino)but-2-enoic acid hydrochloride. Flash chromatography (10 to 50% MeOH in 0.1% TFA in $H_2O$) followed by semi-prep HPLC purification (5% to 50% MeOH in 0.1% TFA in $H_2O$) yielded the desired product as a brown oil (11 mg, 29%) as a TFA salt. $^1$H NMR (400 MHz, MeOD) δ 8.53 (d, *J* = 2.2 Hz, 1H), 7.82 (d, *J* = 8.8 Hz, 1H), 7.43 (s, 1H), 6.92 (dt, *J* = 15.0, 7.2 Hz, 1H), 6.60 (dd, *J* = 15.0, 1.3 Hz, 1H), 4.01 (dd, *J* = 7.2, 1.2 Hz, 2H), 3.72 (t, *J* = 7.0 Hz, 2H), 3.58 (s, 2H), 3.18 – 3.07 (m, 4H), 2.93 (s, 6H), 2.88 (s, 6H), 2.86 (s, 6H), 1.85 – 1.68 (m, 8H), 1.56 – 1.40 (m, 6H). $^{13}$C NMR (176 MHz, MeOD) δ 162.95, 160.98 (q, TFA), 160.28, 153.17, 136.01, 134.79, 132.35, 131.45, 127.98, 118.91 (TFA) 117.25, 117.11 (TFA), 115.59 (TFA), 113.97, 109.81, 57.53, 57.45, 57.39, 42.00, 41.97, 41.86, 41.53, 40.55, 28.32, 28.07, 26.29, 25.87, 24.21, 23.92, 23.35. MS(ES+) m/z 527.4 [M + H]$^+$.

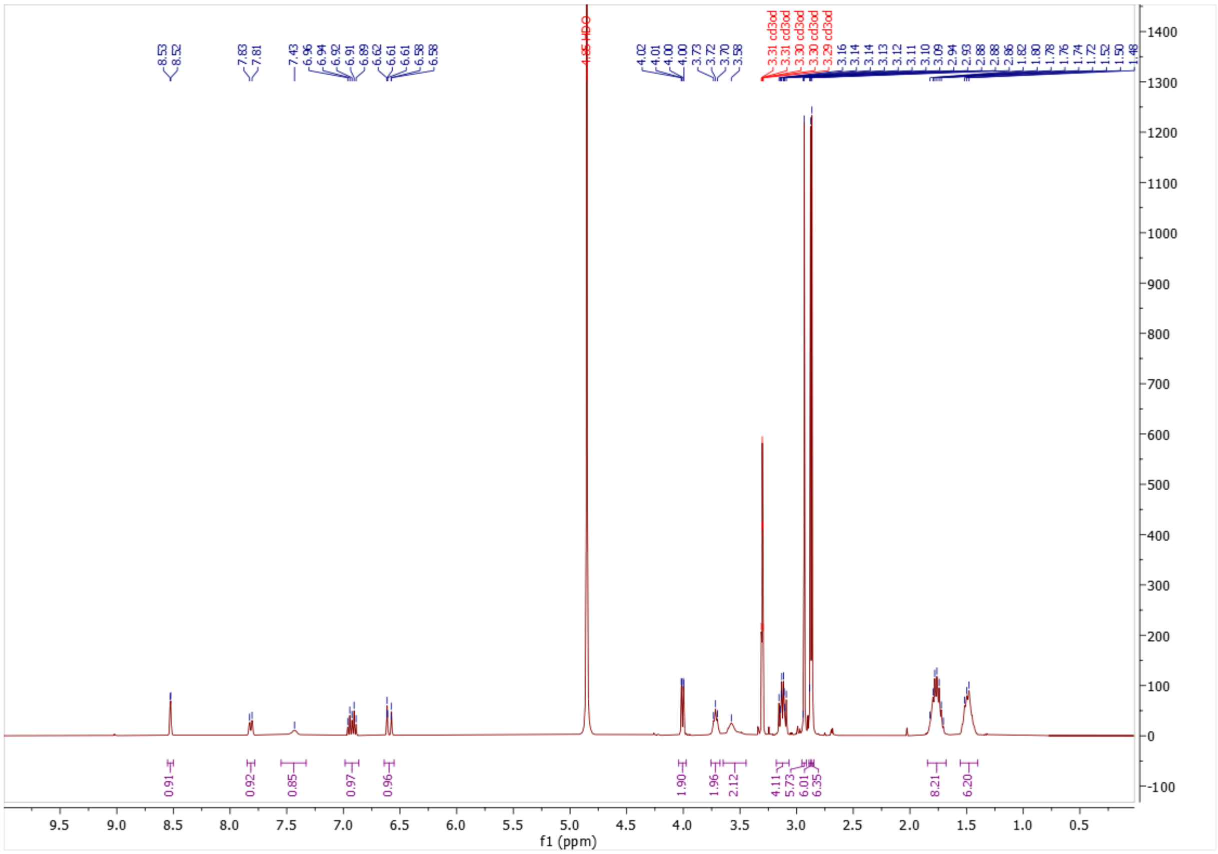
8.53
8.52
7.83
7.81
7.43
6.96
6.94
6.92
6.91
6.89
6.62
6.61
6.61
6.58
6.58
4.85 HDO
4.02
4.01
4.00
4.00
3.73
3.72
3.70
3.58
3.31 cd3od
3.31 cd3od
3.30 cd3od
3.30 cd3od
3.30 cd3od
3.29 cd3od
3.16
3.14
3.14
3.13
3.12
3.11
3.10
3.09
2.94
2.93
2.88
2.86
1.82
1.80
1.78
1.76
1.74
1.72
1.52
1.50
1.48
0.91
0.92
0.85
0.97
0.96
1.90
1.96
2.12
4.11
5.73
6.01
6.35
8.21
6.20
f1 (ppm)

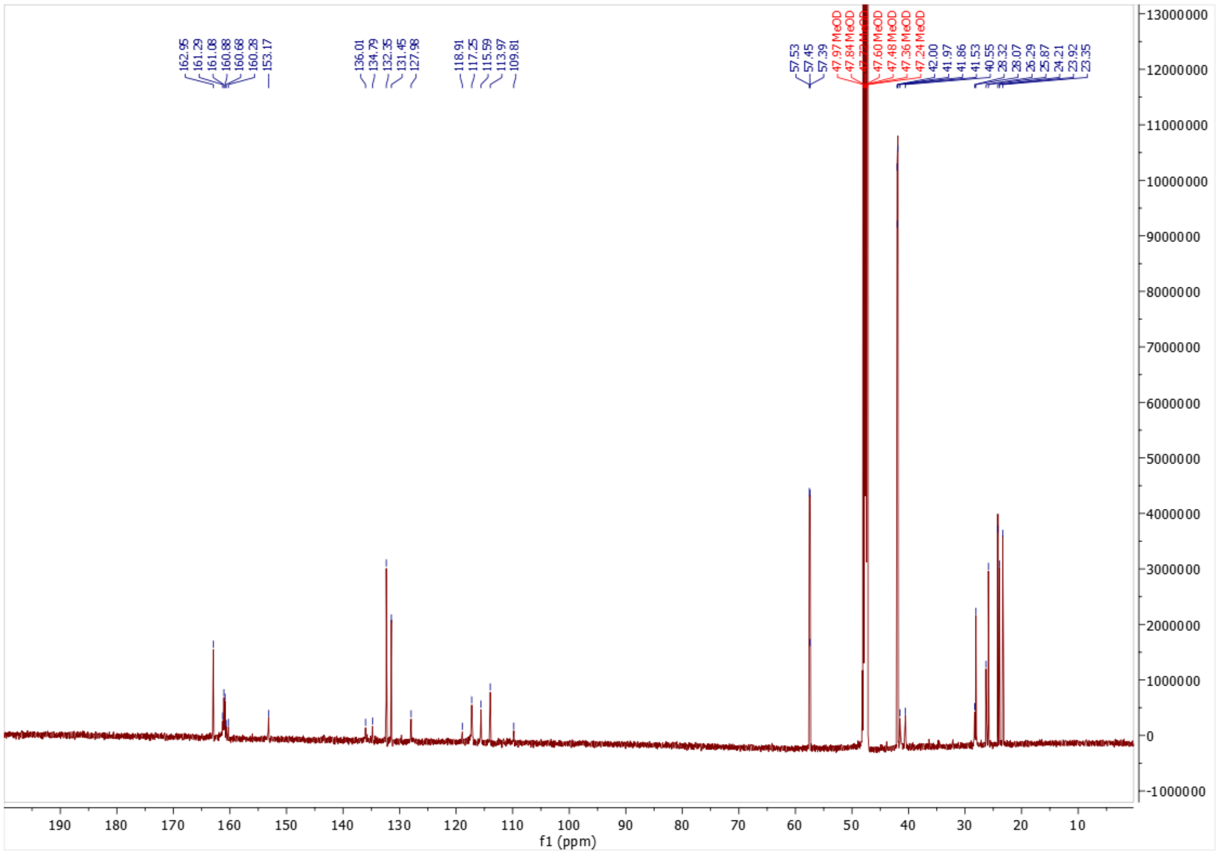
162.95
161.29
161.08
160.88
160.68
160.28
153.17
136.01
134.79
132.35
131.45
127.98
118.91
117.25
115.59
113.97
109.81
57.53
57.45
57.39
47.97 MeOD
47.84 MeOD
47.60 MeOD
47.48 MeOD
47.36 MeOD
47.24 MeOD
42.00
41.97
41.86
41.53
40.55
28.32
28.07
26.29
25.87
24.21
23.92
23.35
13000000
12000000
11000000
10000000
9000000
8000000
7000000
6000000
5000000
4000000
3000000
2000000
1000000
0
-1000000
190
180
170
160
150
140
130
120
110
100
90
80
70
60
50
40
30
20
10
f1 (ppm)

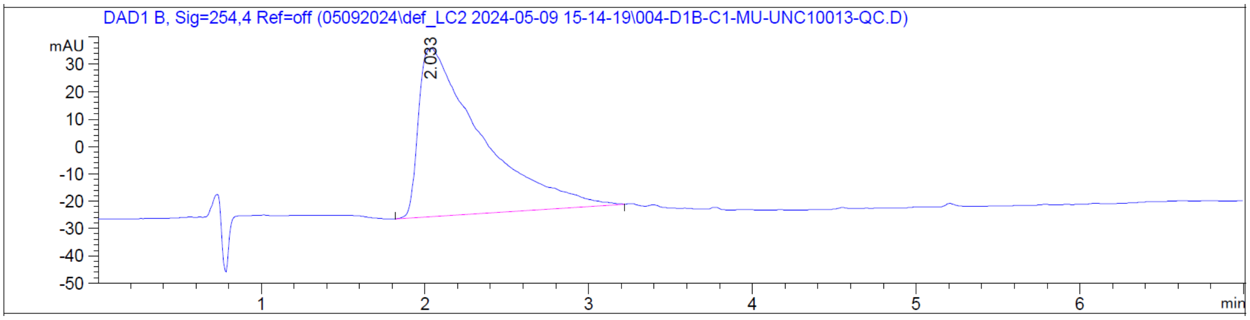
DAD1 B, Sig=254,4 Ref=off (05092024\def_LC2 2024-05-09 15-14-19\004-D1B-C1-MU-UNC10013-QC.D)
mAU
30
20
10
0
-10
-20
-30
-40
-50
2.033
1
2
3
4
5
6
min

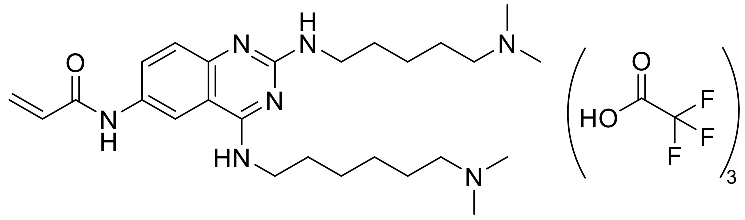

***N*-(4-((6-(dimethylamino)hexyl)amino)-2-((5-(dimethylamino)pentyl)amino)quinazolin-6-yl)acrylamide • 3 TFA salts (UNC10015 (5))**

**UNC10015** was obtained by following General Procedure 1 using **Intermediate 4** and acryloyl chloride. Flash chromatography (10 to 50% MeOH in 0.1% TFA in $H_2O$) followed by semi-prep HPLC purification (5% to 50% MeOH in 0.1% TFA in $H_2O$) yielded the desired product as a light brown oil (1.4 mg, 4%) as a TFA salt. $^1$H NMR (400 MHz, MeOD) δ 8.58 (d, *J* = 2.2 Hz, 1H), 7.71 (d, *J* = 8.7 Hz, 1H), 7.43 (s, 1H), 6.51 – 6.40 (m, 2H), 5.84 (dd, *J* = 8.1, 3.7 Hz, 1H), 3.73 (t, *J* = 6.5 Hz, 2H), 3.58 (m, 2H), 3.16 – 3.09 (m, 4H), 2.89 (s, 6H), 2.87 (s, 6H), 1.86 – 1.70 (m, 8H), 1.56 – 1.43 (m, 6H). MS(ES+) m/z 470.3 [M + H]$^+$.

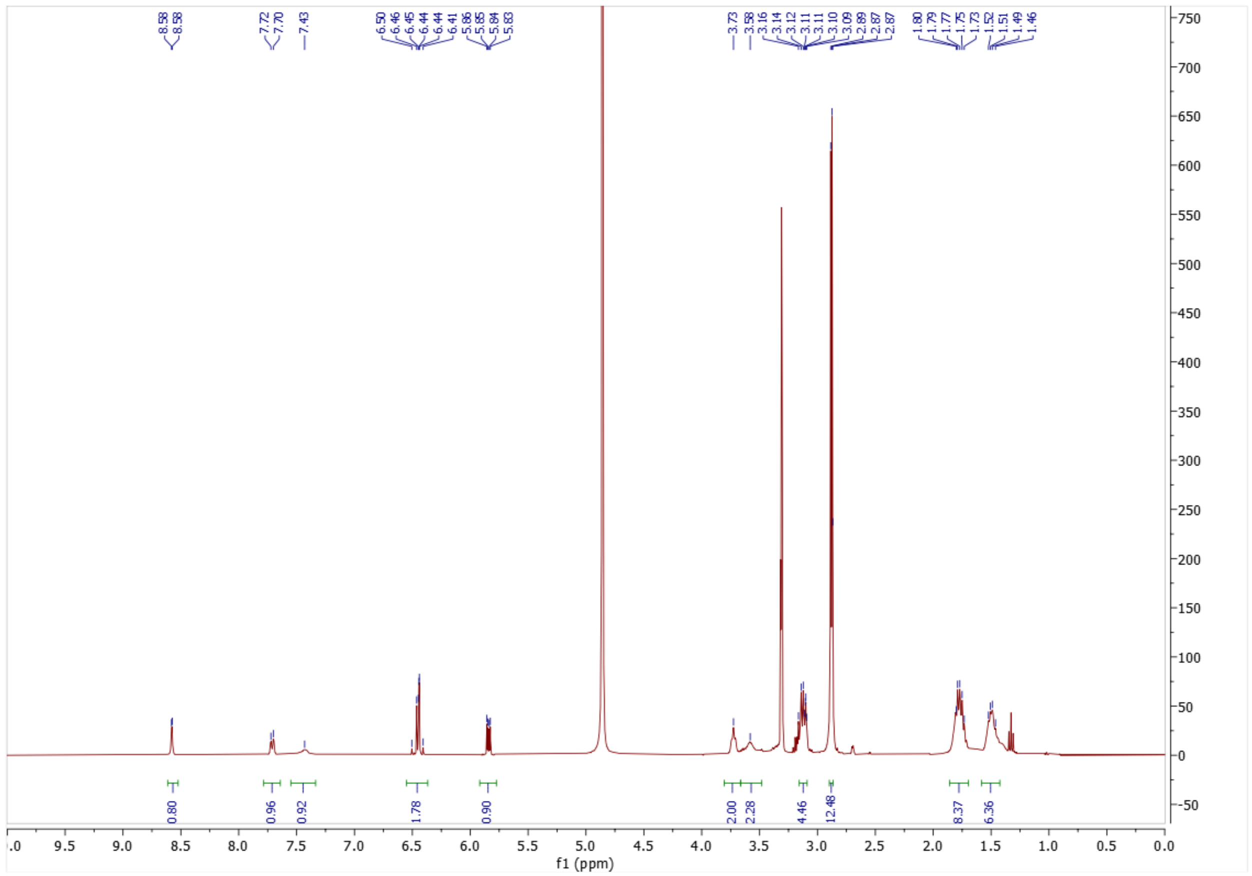

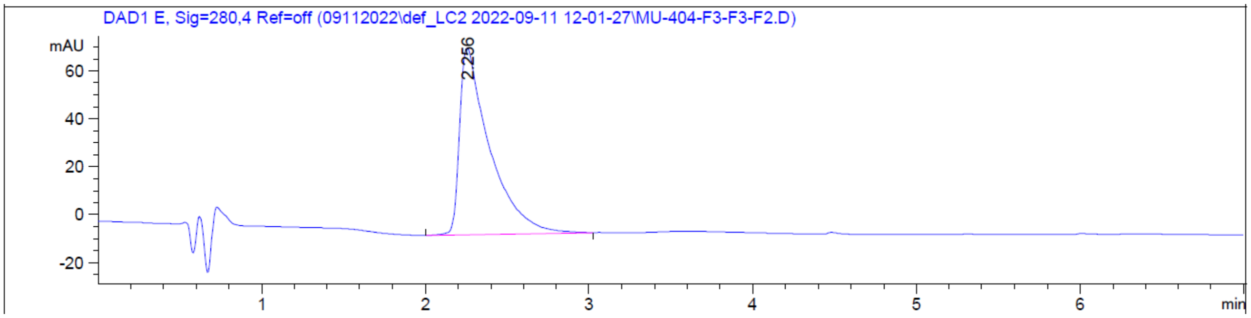

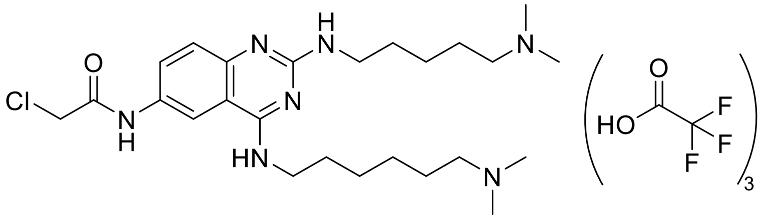

**2-chloro-*N*-(4-((6-(dimethylamino)hexyl)amino)-2-((5-(dimethylamino)pentyl)amino)quinazolin-6-yl)acetamide • 3 TFA salts (UNC9569 (6))**

**UNC9569** was obtained by following General Procedure 1 using **Intermediate 4** and 2-chloroacetyl chloride. Flash chromatography (10 to 50% MeOH in 0.1% TFA in $H_2O$) yielded the desired product as a brown oil (14 mg, 17%) as a TFA salt. $^1H$ NMR (400 MHz, MeOD) δ 8.46 (d, *J* = 2.2 Hz, 1H), 7.72 (d, *J* = 8.8 Hz, 1H), 7.52 – 7.33 (m, 1H), 4.25 (s, 2H), 3.71 (t, *J* = 7.0 Hz, 2H), 3.58 (s, 2H), 3.18 – 3.07 (m, 4H), 2.88 (s, 6H), 2.87 (s, 6H), 1.84 – 1.70 (m, 8H), 1.56 – 1.42 (m, 6H). MS(ES+) m/z 492.3 $[M + H]^+$.

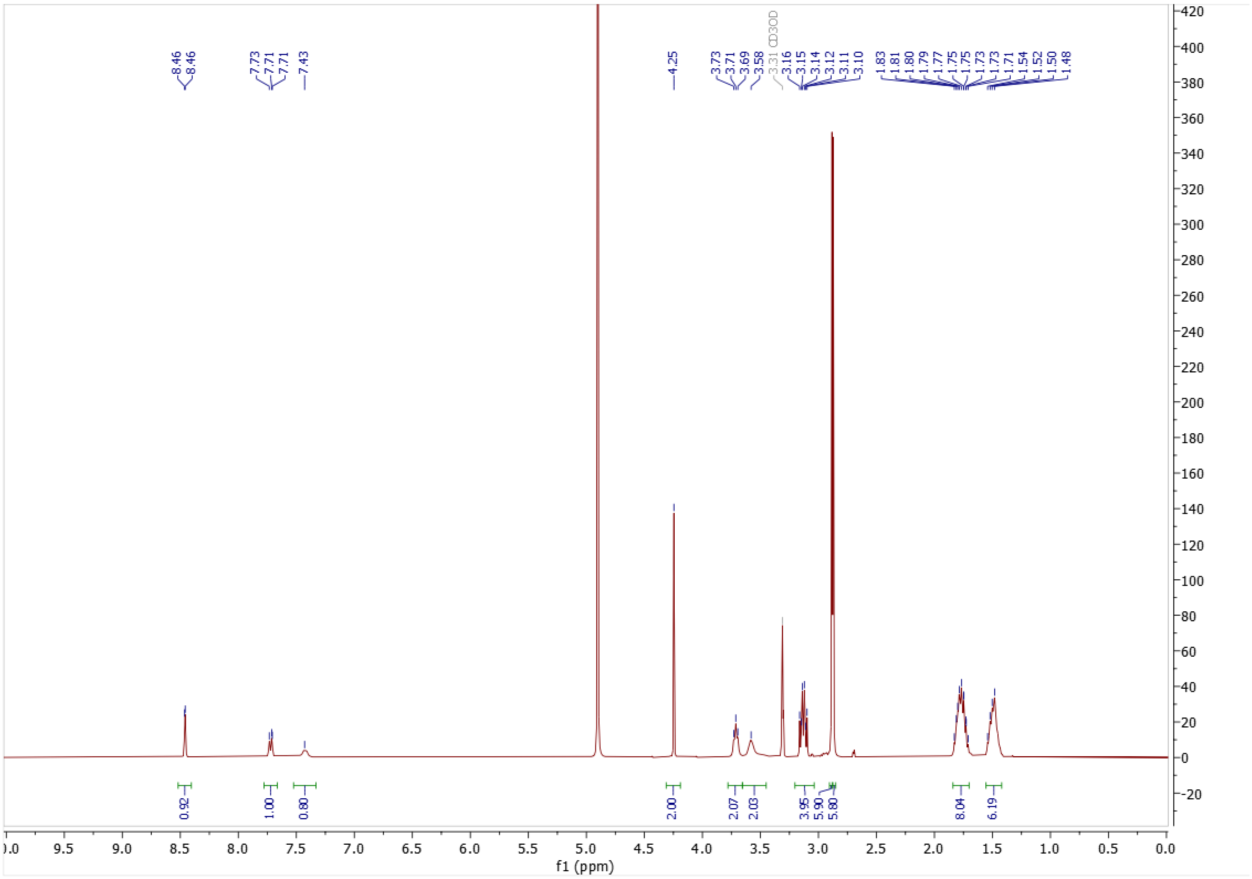

f1 (ppm)

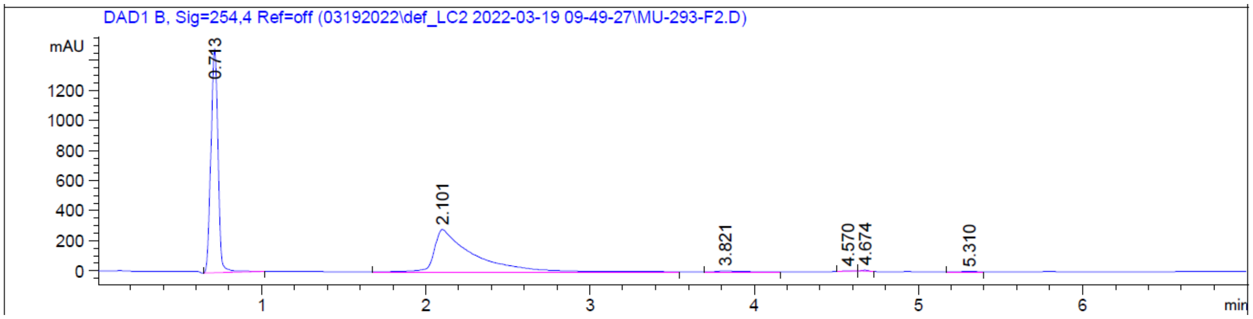

DAD1 B, Sig=254,4 Ref=off (03192022\def_LC2 2022-03-19 09-49-27\MU-293-F2.D)
mAU
min

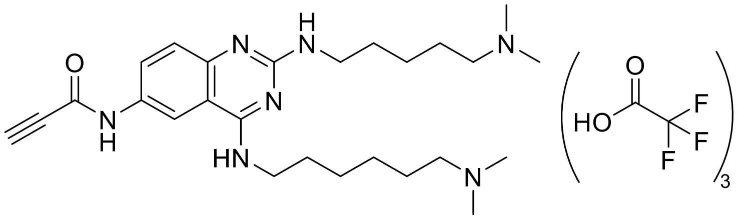

**N-(4-((6-(dimethylamino)hexyl)amino)-2-((5-(dimethylamino)pentyl)amino)quinazolin-6-yl)propiolamide • 3 TFA salts (UNC9970 (7))**

**UNC9970** was obtained by following General Procedure 2 using **Intermediate 4** and propiolic acid. Flash chromatography (10 to 50% MeOH in 0.1% TFA in $H_2O$) yielded the desired product as a yellow oil (2 mg, 5%) as a TFA salt. $^1H$ NMR (400 MHz, MeOD) δ 8.46 (d, *J* = 2.1 Hz, 1H), 7.70 (d, *J* = 8.4 Hz, 1H), 7.42 (s, 1H), 3.85 (s, 1H), 3.78 – 3.64 (m, 2H), 3.58 (s, 2H), 3.18 – 3.08 (m, 4H), 2.89 (s, 6H), 2.87 (s, 6H), 1.85 – 1.69 (m, 8H), 1.57 – 1.41 (m, 6H). MS(ES+) m/z 468.3 $[M + H]^+$.

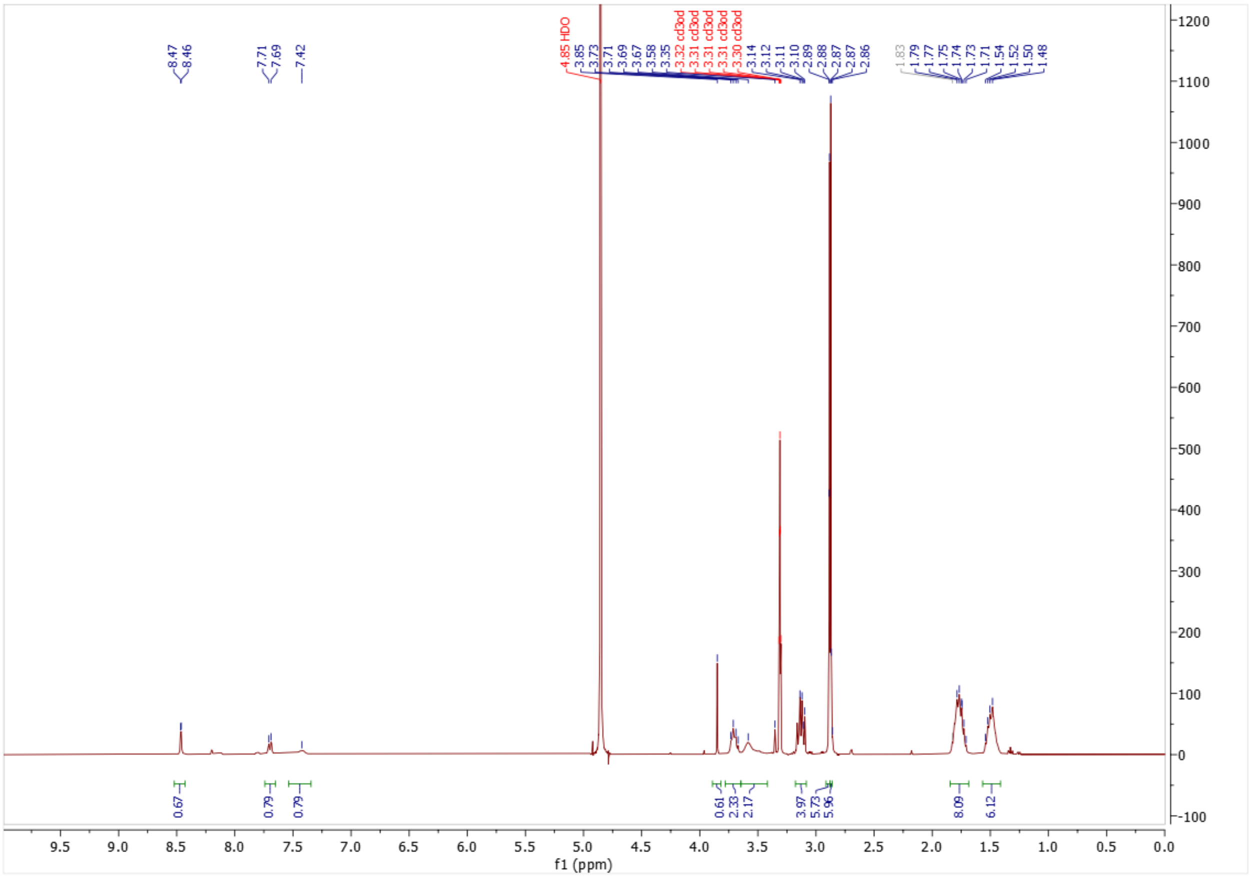

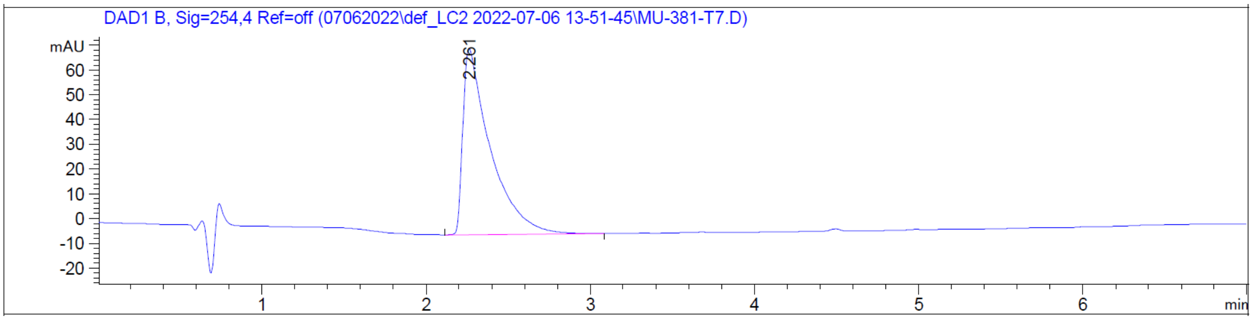

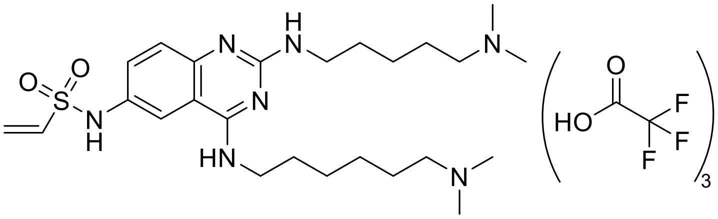

**_N_-(4-((6-(dimethylamino)hexyl)amino)-2-((5-(dimethylamino)pentyl)amino)quinazolin-6-yl)ethenesulfonamide • 3 TFA salts (UNC9773 (8))**

**UNC9773** was obtained by following General Procedure 1 using **Intermediate 4** and chloromethanesulfonyl chloride. Flash chromatography (10 to 50% MeOH in 0.1% TFA in $H_2O$) yielded the desired product as a light yellow oil (16 mg, 29%) as a TFA salt. $^1$H NMR

(400 MHz, MeOD) δ 7.88 (d, *J* = 2.1 Hz, 1H), 7.57 (dd, *J* = 8.8, 2.3 Hz, 1H), 7.40 (s, 1H), 6.72 (dd, *J* = 16.5, 10.0 Hz, 1H), 6.13 (d, *J* = 16.5 Hz, 1H), 5.97 (d, *J* = 10.0 Hz, 1H), 3.70 (t, *J* = 7.2 Hz, 2H), 3.57 (s, 2H), 3.17 – 3.10 (m, 4H), 2.88 (s, 6H), 2.87 (s, 6H), 1.88 – 1.68 (m, 8H), 1.57 – 1.40 (m, 6H). MS(ES+) m/z 506.3 $[M + H]^+$.

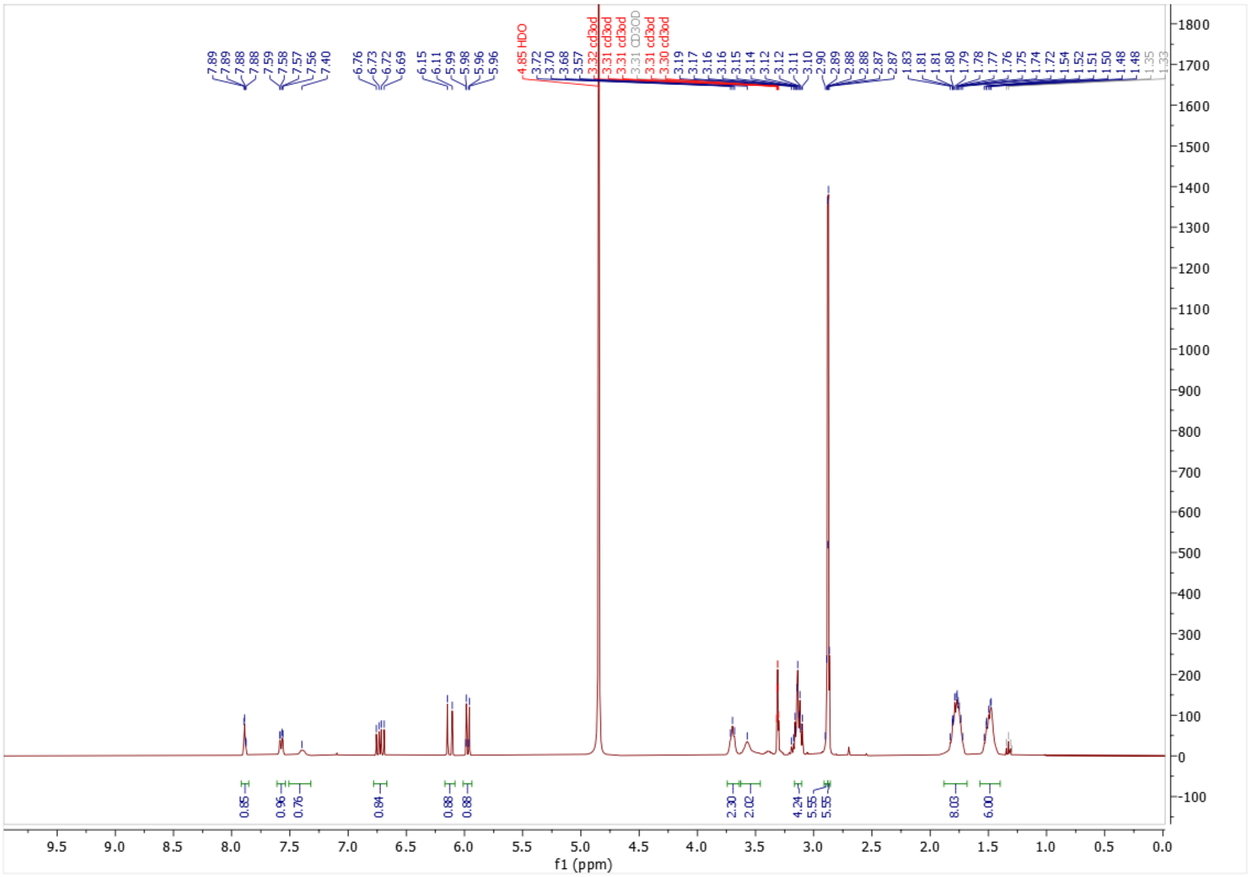

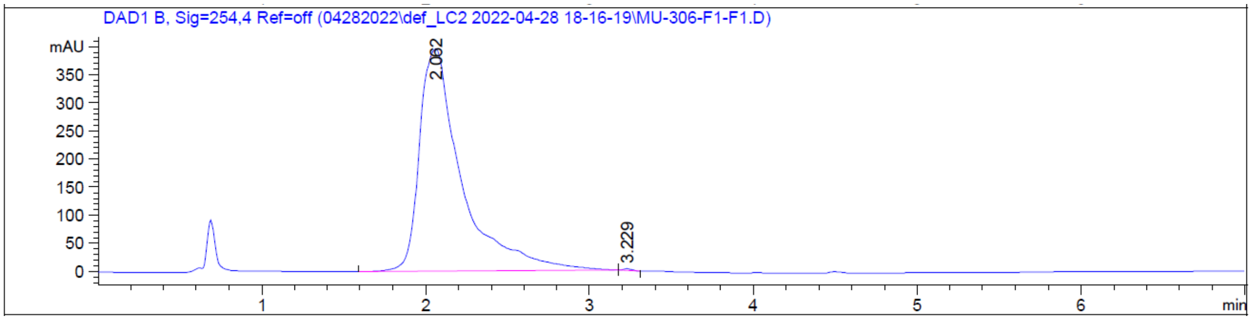

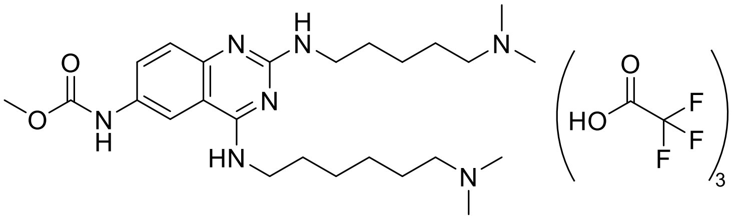

**Methyl (4-((6-(dimethylamino)hexyl)amino)-2-((5-(dimethylamino)pentyl)amino)quinazolin-6-yl)carbamate • 3 TFA salts (UNC9847 (9))**

**UNC9847** was obtained by following General Procedure 1 using **Intermediate 4** and methyl chloroformate. Flash chromatography (10 to 50% MeOH in 0.1% TFA in $H_2O$) yielded the desired product as a light yellow oil (10.2 mg, 17%) as a TFA salt. $^1H$ NMR (400 MHz, MeOD) δ 8.22 (s, 1H), 7.62 (dd, *J* = 8.9, 2.2 Hz, 1H), 7.37 (s, 1H), 3.77 (s, 3H), 3.72 – 3.65 (m, 2H), 3.55 (s, 2H), 3.14 – 3.07 (m, 4H), 2.86 (s, 6H), 2.85 (s, 6H), 1.84 – 1.67 (m, 8H), 1.54 – 1.40 (m, 6H). MS(ES+) m/z 474.4 [M + H]$^+$.

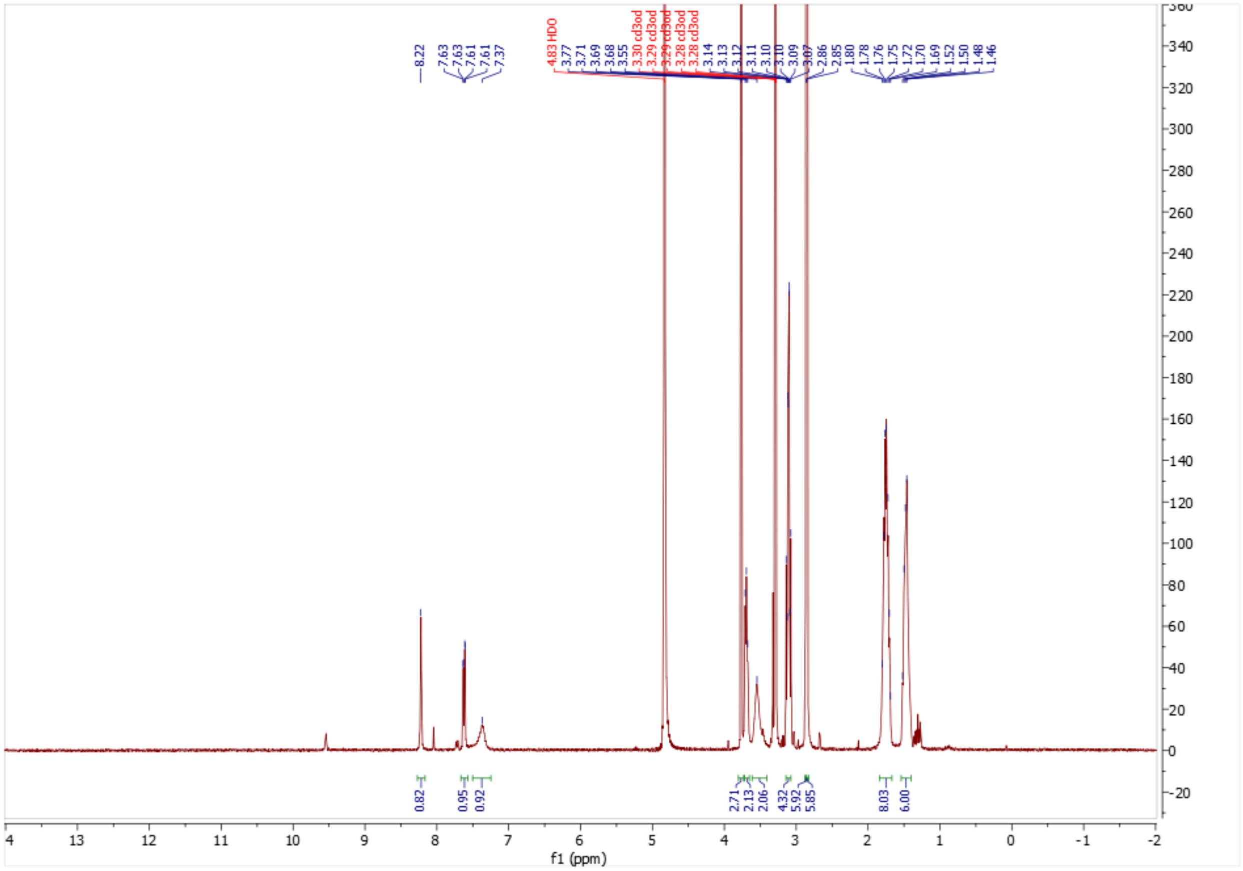

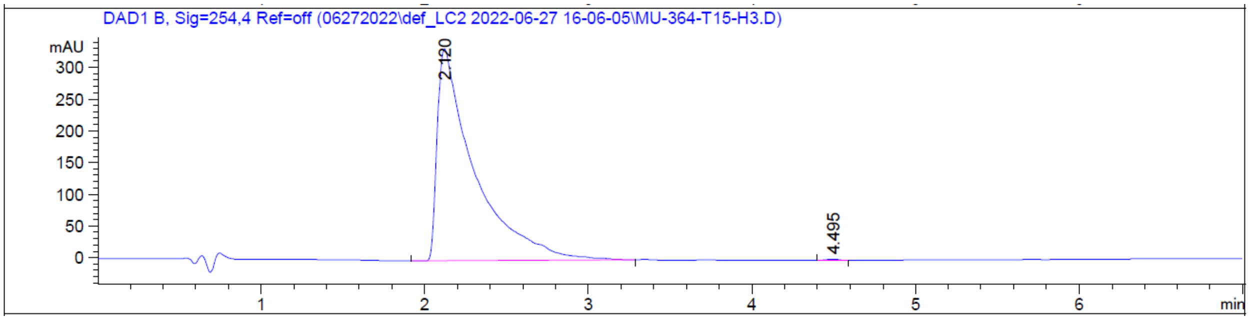

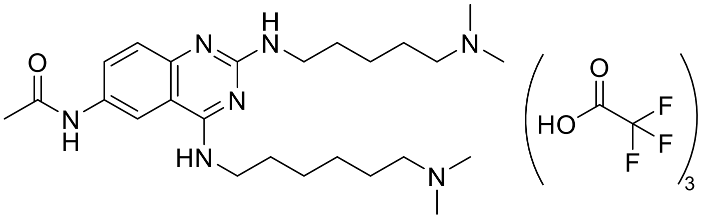

**N-(4-((6-(dimethylamino)hexyl)amino)-2-((5-(dimethylamino)pentyl)amino)quinazolin-6-yl)acetamide • 3 TFA salts (UNC9774 (10))**

**UNC9774** was obtained by following General Procedure 1 using **Intermediate 4** and acetyl chloride. Flash chromatography (10 to 50% MeOH in 0.1% TFA in $H_2O$) yielded the desired product as a yellow oil (31 mg, 30%) as a TFA salt. $^1$H NMR (400 MHz, MeOD) δ 8.39 (d, $J$ = 2.2 Hz, 1H), 7.68 (dd, $J$ = 8.9, 2.2 Hz, 1H), 7.39 (s, 1H), 3.70 (t, $J$ = 7.0 Hz, 2H), 3.57 (s, 2H), 3.19 – 3.08 (m, 4H), 2.88 (s, 6H), 2.87 (s, 6H), 2.18 (s, 3H), 1.86 – 1.68 (m, 8H), 1.57 – 1.39 (m, 6H). MS(ES+) m/z 458.3 $[M + H]^+$.

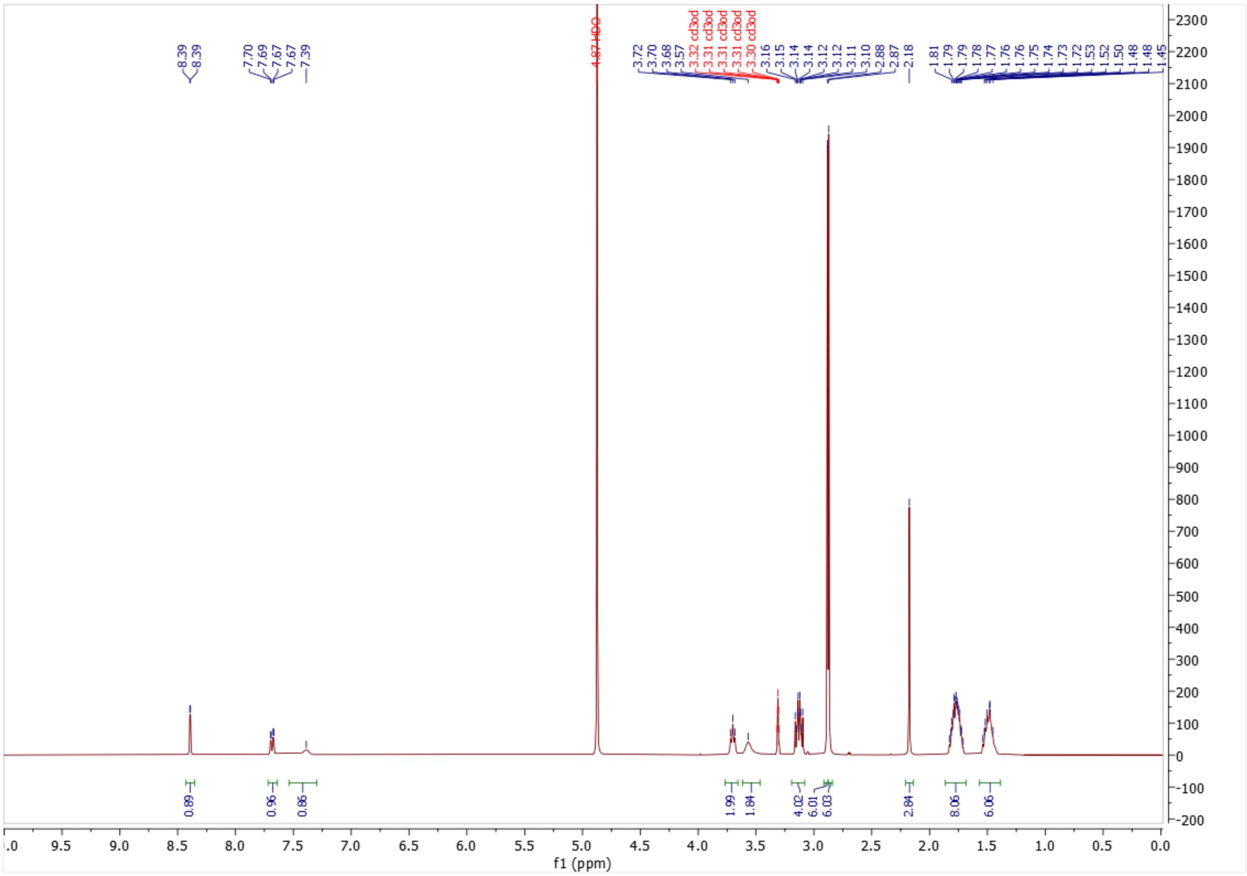

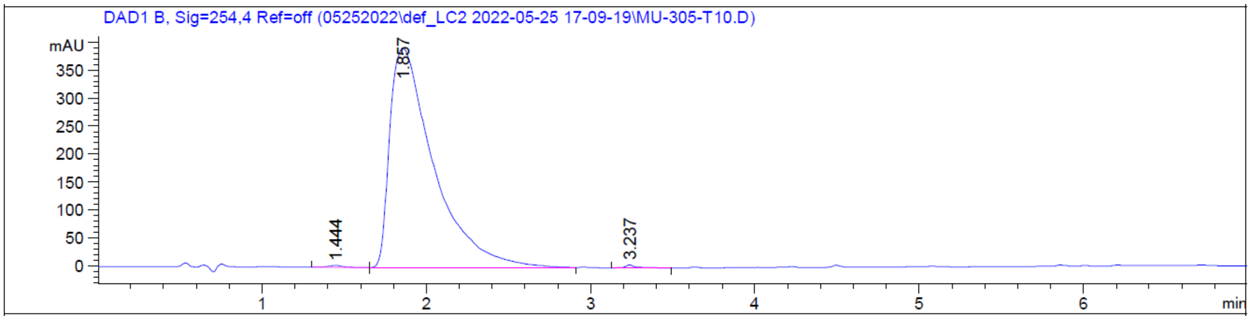

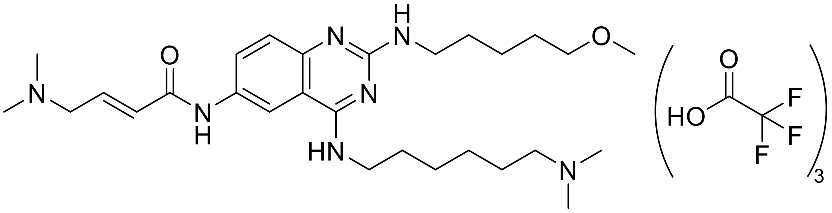

**(*E*)-4-(dimethylamino)-*N*-(4-((6-(dimethylamino)hexyl)amino)-2-((5-methoxypentyl)amino)quinazolin-6-yl)but-2-enamide • 3 TFA salts (UNC11277 (11))**

**UNC11277** was obtained by following General Procedure 2 using **Intermediate 9** and (*E*)-4-(dimethylamino)but-2-enoic acid hydrochloride. Flash chromatography (10 to 50% MeOH in 0.1% TFA in $H_2O$) followed by semi-prep HPLC purification (5% to 50% MeOH in 0.1% TFA in $H_2O$) yielded the desired product as a dark brown oil (8.2 mg, 12%) as a

TFA salt. $^1$H NMR (400 MHz, MeOD) δ 8.53 (d, *J* = 2.2 Hz, 1H), 7.82 (d, *J* = 8.9 Hz, 1H), 7.42 (s, 1H), 6.93 (dt, *J* = 14.8, 7.3 Hz, 1H), 6.60 (dt, *J* = 15.2, 1.3 Hz, 1H), 4.01 (dd, *J* = 7.3, 1.3 Hz, 2H), 3.76 – 3.67 (m, 2H), 3.55 (s, 2H), 3.42 (t, *J* = 6.4 Hz, 2H), 3.16 – 3.08 (m, 2H), 2.94 (s, 6H), 2.87 (s, 6H), 1.86 – 1.68 (m, 6H), 1.68 – 1.59 (m, 2H), 1.58 – 1.42 (m, 6H). MS(ES+) m/z 514.4 [M + H]$^+$.

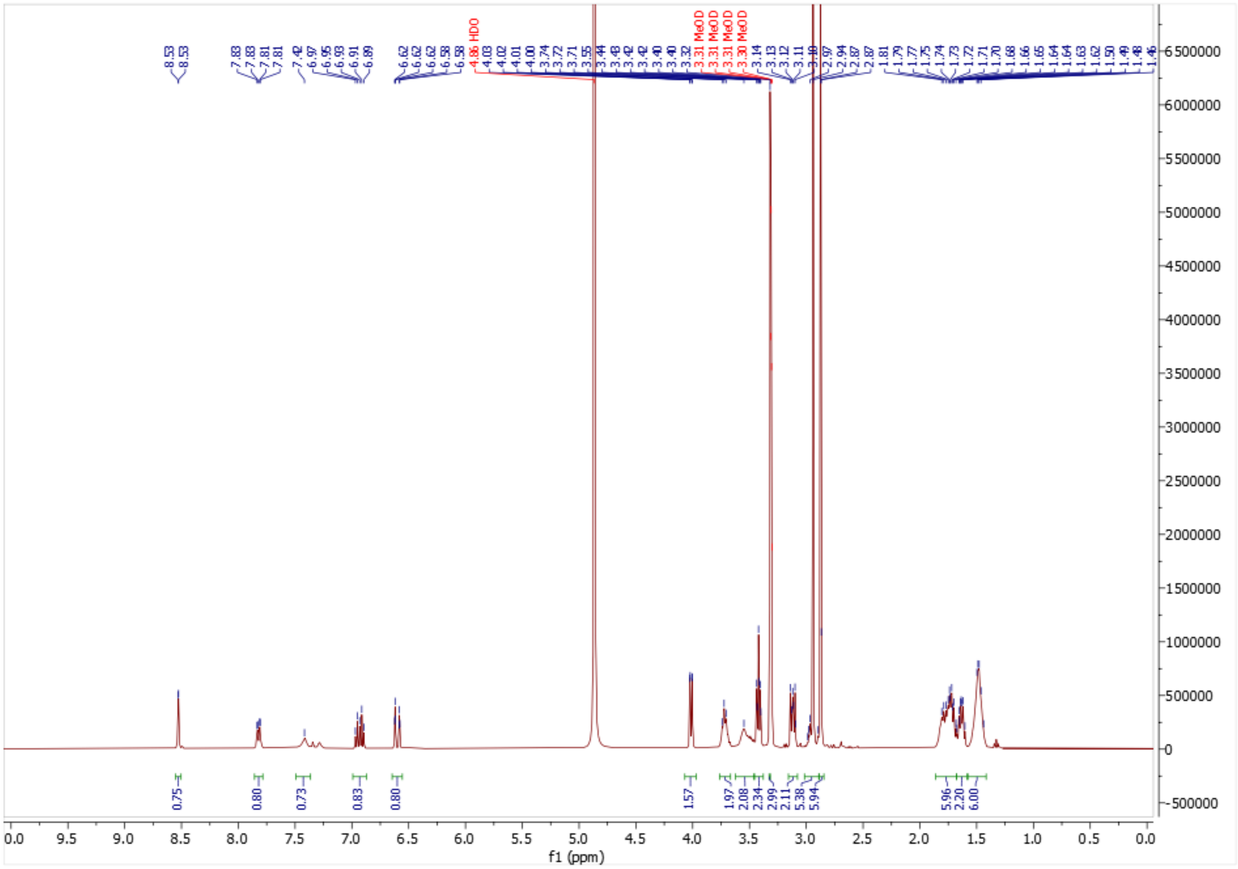

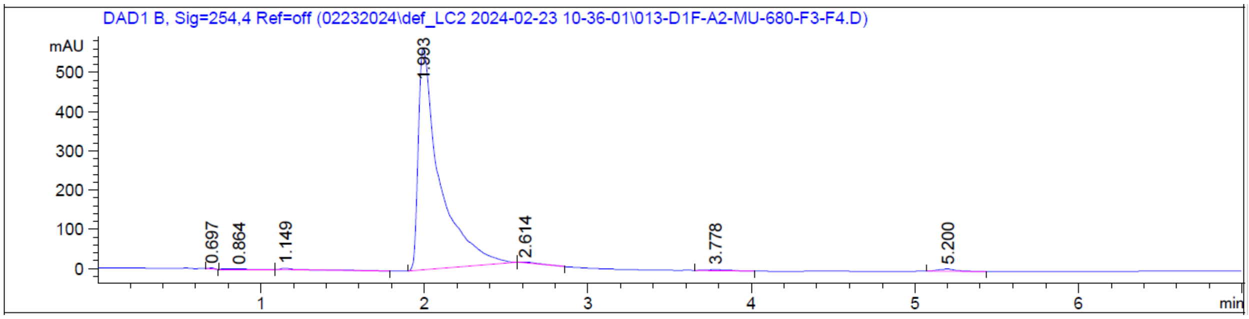

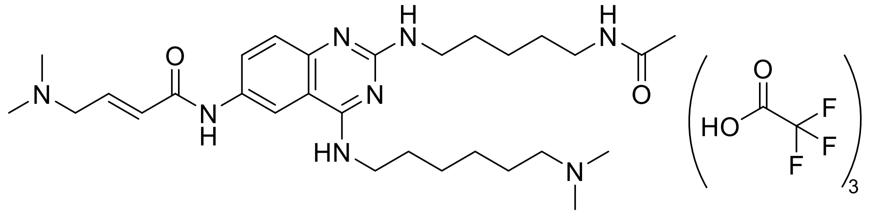

**(*E*)-*N*-(2-((5-acetamidopentyl)amino)-4-((6-(dimethylamino)hexyl)amino)quinazolin-6-yl)-4-(dimethylamino)but-2-enamide • 3 TFA salts (UNC11366 (12))**

**UNC11366** was obtained by following General Procedure 2 using **Intermediate 11** and (*E*)-4-(dimethylamino)but-2-enoic acid hydrochloride. Flash chromatography (10 to 80% MeOH in 0.1% TFA in $H_2O$) followed by semi-prep HPLC purification (5% to 80% MeOH in 0.05% TFA in $H_2O$) yielded the desired product as a dark brown oil (9.0 mg, 9%) as a TFA salt. $^1$H NMR (400 MHz, MeOD) δ 8.53 (d, *J* = 2.2 Hz, 1H), 7.82 (dd, *J* = 9.0, 2.2 Hz, 1H), 7.42 (s, 1H), 6.93 (dt, *J* = 14.8, 7.3 Hz, 1H), 6.60 (dt, *J* = 15.3, 1.3 Hz, 1H), 4.02 (dd, *J* = 7.3, 1.3 Hz, 2H), 3.72 (t, *J* = 7.4 Hz, 2H), 3.55 (s, 2H), 3.18 (t, *J* = 7.0 Hz, 2H), 3.15 – 3.08 (m, 2H), 2.94 (s, 6H), 2.88 (s, 6H), 1.92 (s, 3H), 1.83 – 1.66 (m, 6H), 1.63 – 1.51 (m, 3H), 1.51 – 1.42 (m, 5H). MS(ES+) m/z 541.4 [M + H]$^+$.

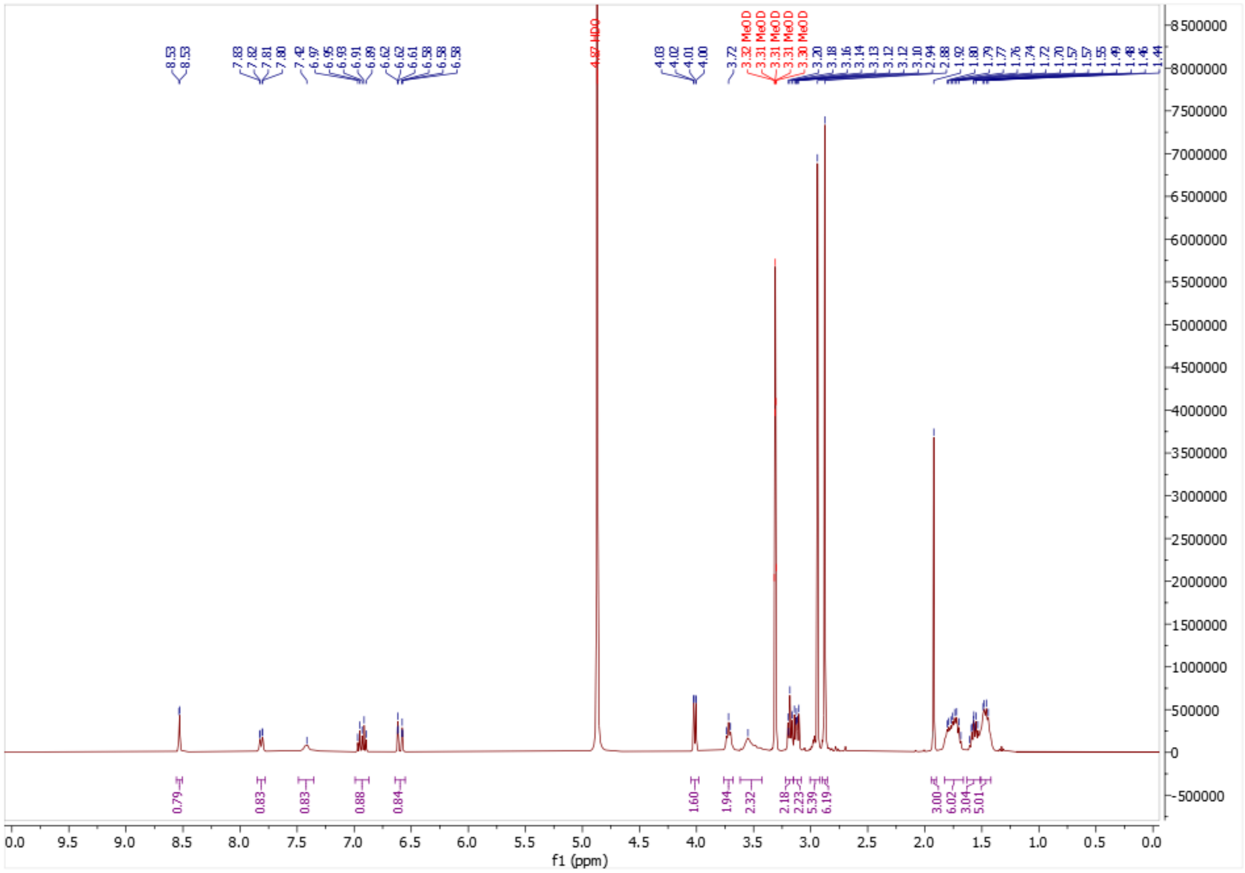

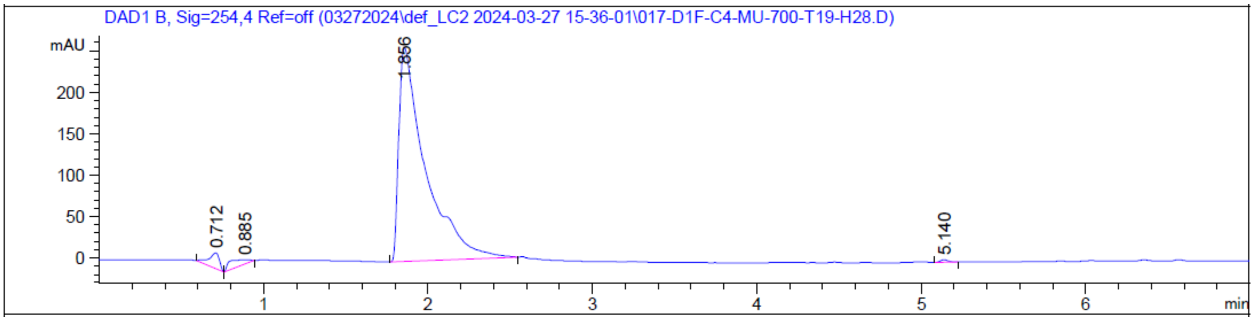

DAD1 B, Sig=254,4 Ref=off (03272024\def_LC2 2024-03-27 15-36-01\017-D1F-C4-MU-700-T19-H28.D)
mAU
200
150
100
50
0
0.712
0.885
1.856
5.140
1
2
3
4
5
6
min

#### 2. Crystallography

**Table 1. Data collection and refinement statistics.**

| | SETDB1-UNC10013 | SETDB1-UNC10016 | SETDB1-UNC6535 |
|---|---|---|---|
| PDB code | 9CUW | 9CUX | 8G5E |
| **Data collection** | | | |
| Space group | $P2_12_12_1$ | $P2_12_12_1$ | $P2_12_12_1$ |
| Cell dimensions | | | |
| *a*, *b*, *c* (Å) | 37.15, 55.13, 118.37 | 52.83,60.66,70.01 | 54.01,62.00,69.34 |
| a, b, g (°) | 90.0, 90.0, 90.0 | 90.0,90.0,90.0 | 90.0,90.0,90.0 |
| Resolution (Å) | 50.0-1.53 (1.56-1.53) * | 50.0-1.27 (1.29-1.27) * | 50.0-1.98 (2.01-1.98) * |
| $R_{sym}$ or $R_{merge}$ | 0.087 (0.938) | 0.052(0.916) | 0.102(0.677) |
| CC1/2 | 0.996(0.832) | 1.000(0.719) | 0.993(0.724) |
| I / sI | 30.3(2.0) | 43.4(2.0) | 16.0(1.8) |
| Completeness (%) | 100.0(100.0) | 100.0(99.9) | 97.2(90.0) |
| Redundancy | 12.2(11.2) | 11.7(8.5) | 5.0(3.9) |
| **Refinement** | | | |
| Resolution (Å) | 40.37-1.53 (1.57-1.53) | 45.89-1.27 (1.30-1.27) | 31.02-1.98 (2.03-1.98) |
| No. reflections | 36104 | 57040 | 15452 |
| $R_{work}$ / $R_{free}$(%) | 16.2/19.5 | 15.9/18.1 | 20.0/24.9 |
| No. atoms | 2058 | 2174 | 1874 |
| Protein | 1783 | 1831 | 1734 |
| Ligand/ion | 38 | 35 | 33 |
| Water | 237 | 308 | 100 |
| *B*-factors | 26.7 | 16.5 | 28.2 |
| Protein | 24.9 | 14.1 | 27.9 |
| Ligand/ion | 29.9 | 12.6 | 37.5 |
| Water | 38.7 | 29.4 | 29.7 |
| R.m.s. deviations | | | |
| Bond lengths (Å) | 0.005 | 0.005 | 0.010 |
| Bond angles (°) | 1.22 | 1.269 | 1.368 |

*Values in parentheses are for highest-resolution shell.

#### 3. Biological Evaluation

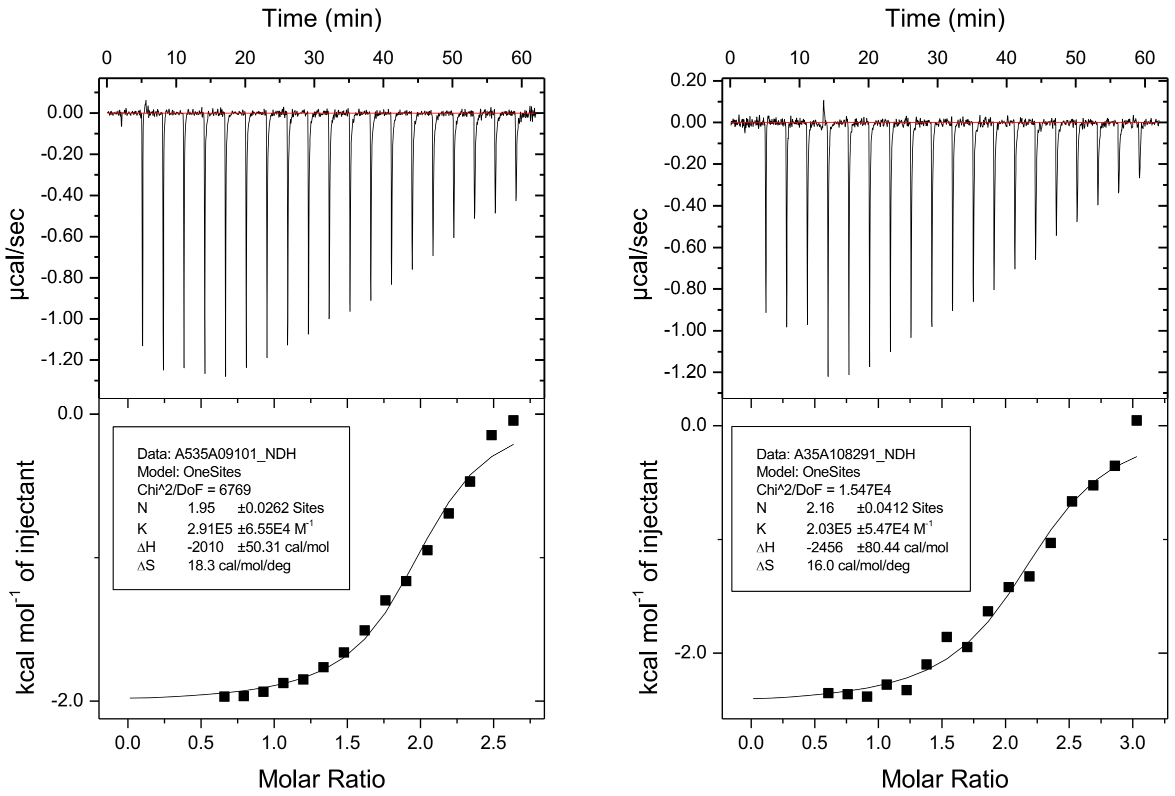

**Figure S1. Isothermal titration calorimetry results for UNC6535 and SETDB1 3TD.**

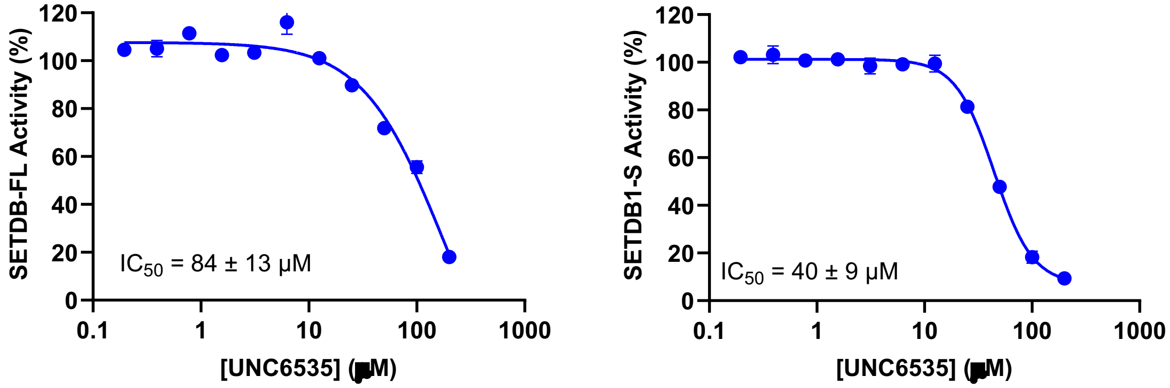

**Figure S2. Radiometric methyltransferase activity assay for UNC6535 using SETDB1-FL (aa 1-1291) or SETDB1-S (aa 570-1291).** Values are reported as the average of three independent experiments ± s.d.

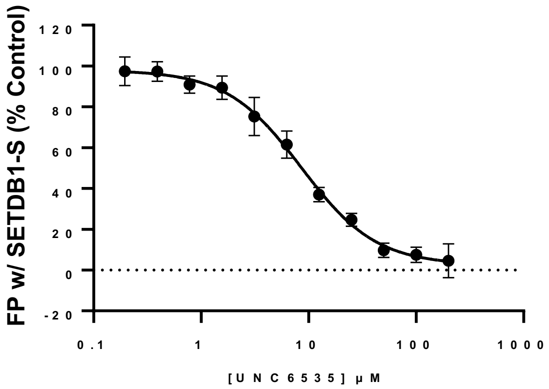

**Figure S3. Fluorescence polarization results for UNC6535 using SETDB1-S (aa 570-1291).** Values are reported as the average of three independent experiments ± s.d.

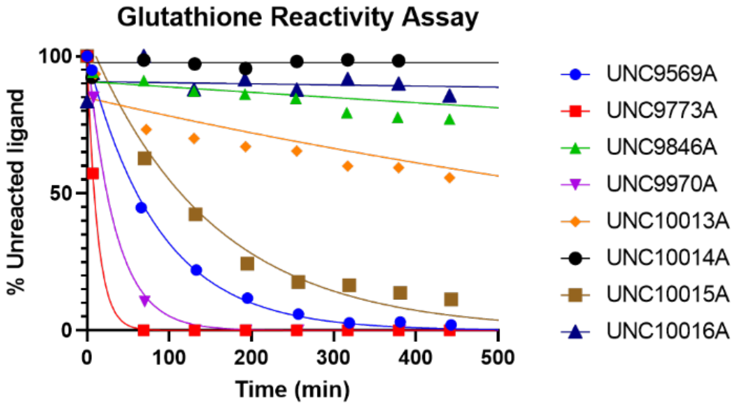

**Figure S4. Evaluation of the intrinsic reactivity of 1-8 towards glutathione using LC-MS quantification.**

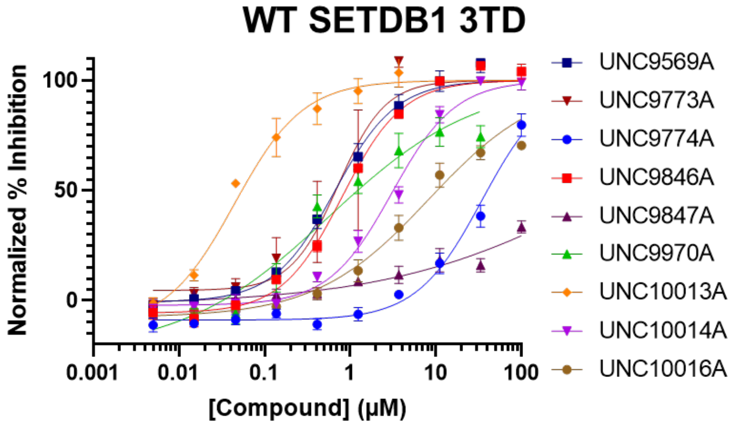

**Figure S5. TR-FRET displacement of H3K9Me2K14Ac for 1-8 using the WT SETDB1 3TD.**

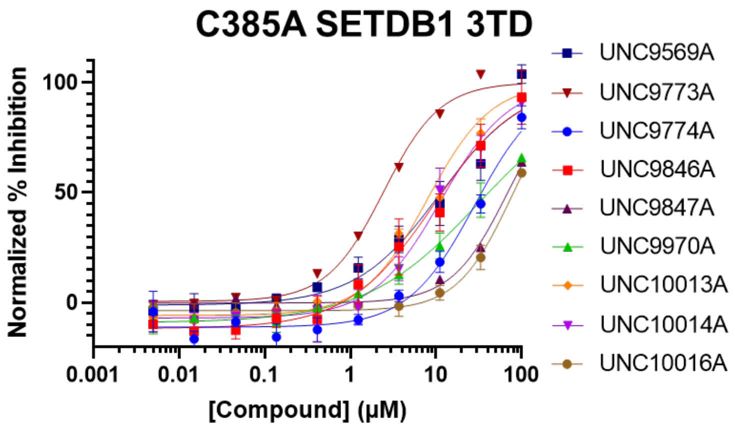

**Figure S6. TR-FRET displacement of H3K9Me2K14Ac for** 1-8 **using C385A SETDB1 3TD.**

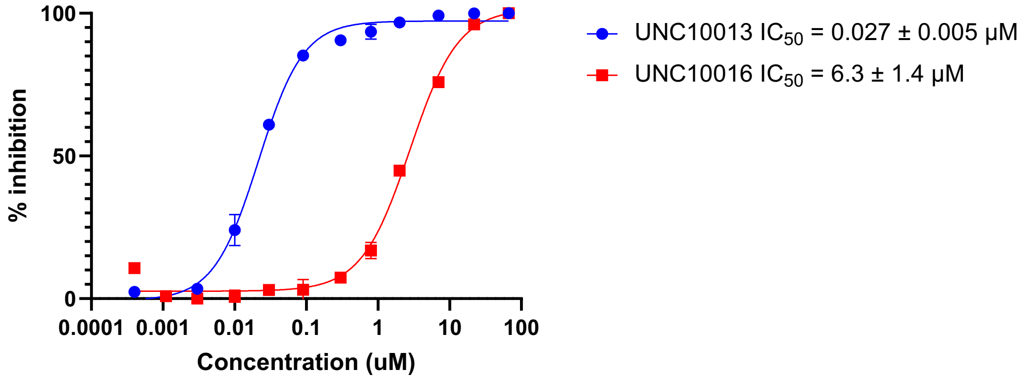

**Figure S7. TR-FRET displacement of H3K9Me2K14Ac-modified nucleosome for UNC10013 (4) and UNC10016 (2) using the WT SETDB1 3TD.** Values are reported as the average of three independent experiments ± s.e.m.

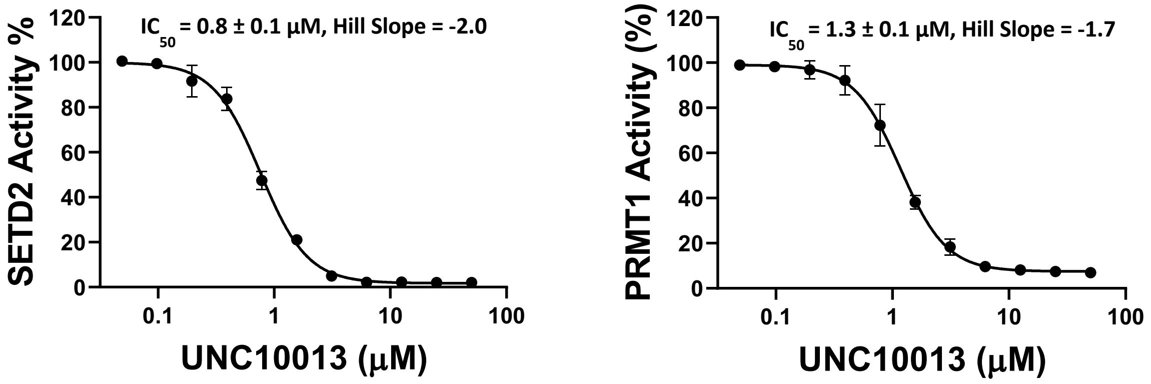

**Figure S8. Dose-response evaluation of UNC10013 (4) ability to inhibit the methyltransferase ability of of SETD2 (left) and PRMT1 (right).** Values are reported as the average of three independent experiments ± s.d.

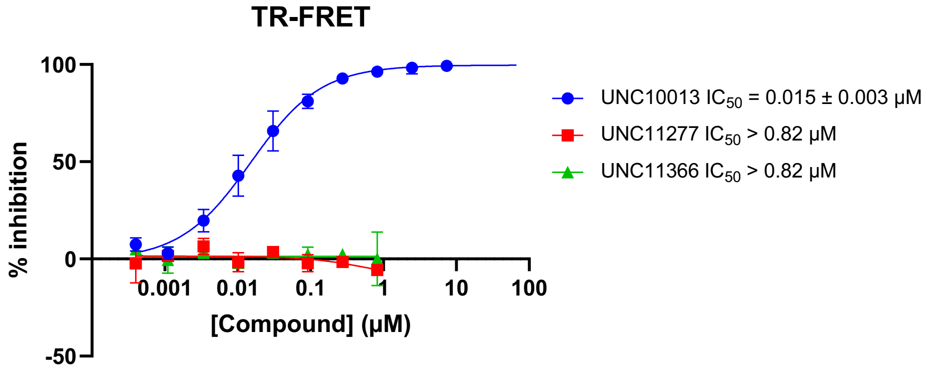

**Figure S9. TR-FRET displacement of H3K9Me2K14Ac peptide for UNC10013 (4), UNC11277 (11) and UNC11366 (12) using the WT SETDB1 3TD.** Values are reported as the average of three independent experiments ± s.e.m.

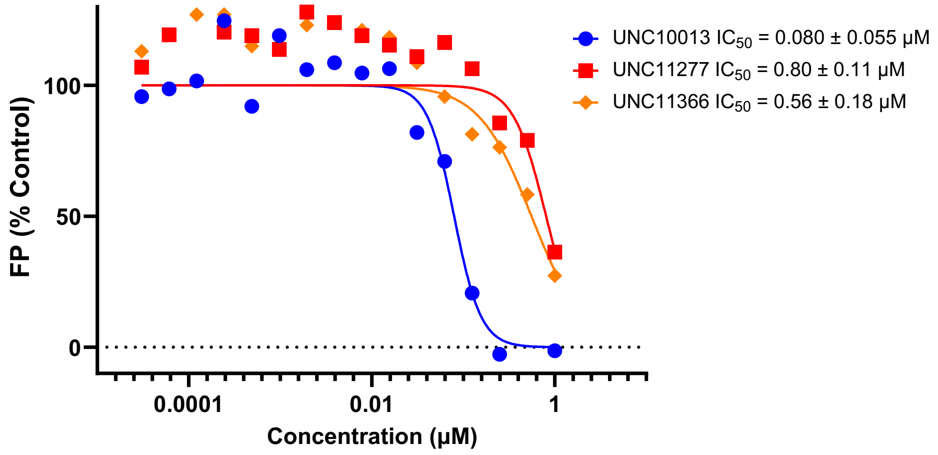

**Figure S10. FP displacement of FITC-H3K9Me2K14Ac peptide for UNC10013 (4), UNC11277 (11) and UNC11366 (12) using the WT SETDB1 3TD.** Values are reported as the average of three independent experiments ± s.d.

| MTase | % inhibition | | | |
|---|---|---|---|---|
| | UNC10013 (4) | | UNC10016 (2) | |
| | 10 µM | 50 µM | 10 µM | 50 µM |
| **G9A** | 63 | 88 | 25 | 55 |
| **GLP** | 57 | 84 | 20 | 53 |
| **SETDB1** | 2 | 62 | -4 | 15 |
| **SUV39H1** | 48 | 76 | 32 | 65 |
| **SUV39H2** | 5 | 19 | -4 | 1 |
| **SUV420H1** | 66 | 86 | 28 | 67 |
| **SUV420H2** | 42 | 79 | 20 | 57 |
| **SETD7** | 24 | 59 | 17 | 62 |
| **SETD8** | 0 | 3 | 1 | 5 |
| **MLL1** | 7 | 52 | 3 | 16 |
| **MLL3** | 1 | 35 | 12 | 28 |
| **PRDM9** | 61 | 84 | 47 | 80 |
| **SETD2** | 89 | 97 | 65 | 95 |
| **PRC2** | 52 | 84 | 35 | 72 |
| **SMYD2** | 48 | 77 | 32 | 63 |
| **SMYD3** | 2 | 31 | 1 | 2 |
| **PRMT1** | 93 | 97 | 65 | 93 |
| **PRMT3** | 79 | 92 | 34 | 86 |
| **PRMT4** | 52 | 79 | 57 | 78 |
| **PRMT5** | 8 | 11 | 3 | 3 |
| **PRMT7** | 46 | 85 | 30 | 69 |
| **PRMT6** | 80 | 89 | 55 | 82 |
| **PRMT8** | 51 | 87 | 37 | 73 |
| **PRMT9** | 57 | 81 | 42 | 70 |
| **DNMT1** | 3 | 2 | 3 | 1 |
| **BCDIN3D** | 76 | 91 | 53 | 71 |
| **DNMT3A/3L** | -69 | 6 | -80 | 6 |
| **DNMT3B/3L** | -30 | -10 | -26 | -21 |
| **NSD1** | -631 | -740 | -619 | -727 |
| **NSD2** | -485 | -1397 | -456 | -1497 |
| **NSD3** | -920 | -1559 | -935 | -1536 |
| **ASH1L** | -708 | -1923 | -600 | -2012 |
| **DOT1L** | -613 | -4797 | -645 | -2154 |

**Table S1.** Methyltransferase selectivity evaluation of UNC10013 (**4**) and UNC10016 (**2**). Compounds were tested at 10 and 50 µM. Light orange shows proteins with 60-80% inhibition, dark orange shows proteins with over 80% inhibition.

| Gene Symbol | Site Position | Protein ID | Description | Motif | Max Score | Redundancy | Sequence | Num Quant | Summed S/N, Column Normalized, Scaled to 100 per Site | | | | | | Ratio dmso/13a |
|---|---|---|---|---|---|---|---|---|---|---|---|---|---|---|---|
| | | | | | | | | | spike_dmso_1 | spike_dmso_2 | spike_dmso_3 | spike_13a_1 | spike_13a_2 | spike_13a_3 | |
| SETDB1 | 53 | Q15047 | Histone-lysine N-methyltransferase SETDB1 | ELEKMDCVQQRKK | 5000 | R | K.M*DC#VQQR.K | 17 | 9.876776339 | 9.610613557 | 9.955246641 | 10.35041232 | 10.26313246 | 10.37476095 | **0.950120888** |
| SETDB1 | 1207 | Q15047 | Histone-lysine N-methyltransferase SETDB1 | YDGEESCYIIDAK | 5000 | R | R.QFYDGEESC#YIIDAK.L | 13 | 9.077856367 | 9.556721149 | 10.28106705 | 10.46058615 | 10.17422095 | 10.26437786 | **0.935806061** |
| SETDB1 | 773 | Q15047 | Histone-lysine N-methyltransferase SETDB1 | YKRLEECLPTGVY | 361 | R | R.LEEC#LPTGVYECNKR.C | 12 | 8.772255103 | 8.961067856 | 9.605256119 | 10.47539963 | 10.34348602 | 10.33315177 | **0.877585588** |
| SETDB1 | 92 | Q15047 | Histone-lysine N-methyltransferase SETDB1 | SRAVTNCESLVKD | 5000 | R | R.AVTNC#ESLVK.D | 11 | 9.08056386 | 8.770898737 | 9.310676071 | 9.921693944 | 10.01348126 | 9.678118032 | **0.917227897** |
| SETDB1 | 968 | Q15047 | Histone-lysine N-methyltransferase SETDB1 | SIPVGGCNPPSSE | 5000 | R | K.DSHPPDLGPPHIPVPPSIPVGGC#NPPSSEETPK.N | 11 | 9.053456776 | 9.438923095 | 10.10998795 | 9.674170922 | 9.806716651 | 9.2431889 | **0.995762835** |
| SETDB1 | 1226 | Q15047 | Histone-lysine N-methyltransferase SETDB1 | RYLNHSCSPNLFV | 5000 | R | R.YLNHSC#SPNLFVQNVFVDTHDLR.F | 8 | 8.581681042 | 8.974633332 | 9.776665395 | 9.058062035 | 9.831560861 | 8.996653548 | **0.980158818** |
| SETDB1 | 987 | Q15047 | Histone-lysine N-methyltransferase SETDB1 | VASWLSCNSVSEG | 5000 | R | K.VASWLSC#NSVSEGGFADSDSHSSFK.T | 7 | 9.32039309 | 9.725080877 | 10.02624672 | 10.19300494 | 10.51052921 | 9.619849844 | **0.958722835** |
| SETDB1 | 1279 | Q15047 | Histone-lysine N-methyltransferase SETDB1 | EGKELLCCGAIE | 91 | R | K.ELLC#CCGAIECR.G | 7 | 7.25519752 | 7.481946682 | 7.622081283 | 8.267424393 | 7.968022422 | 7.774109227 | **0.931263595** |
| SETDB1 | 781 | Q15047 | Histone-lysine N-methyltransferase SETDB1 | PTGVYECNKRCKC | 527 | R | K.RLEECLPTGVYEC#NK.R | 6 | 8.36695891 | 8.094644555 | 9.614661456 | 9.446075842 | 9.890989754 | 9.565860357 | **0.902201561** |
| SETDB1 | 819 | Q15047 | Histone-lysine N-methyltransferase SETDB1 | KGWGIRCLDDIAK | 5000 | R | R.C#LDDIAK.G | 5 | 9.838606155 | 10.17719606 | 10.42702545 | 10.54869033 | 10.76845894 | 10.43433753 | **0.95878432** |
| SETDB1 | 385 | Q15047 | Histone-lysine N-methyltransferase SETDB1 | FLDDKRCEWIYRG | 5000 | R | K.RC#EWIYR.G | 4 | 21.40699098 | 20.5160461 | 20.8173267 | 2.958892641 | 2.864808523 | 2.410456234 | **7.61952447** |
| SETDB1 | 876 | Q15047 | Histone-lysine N-methyltransferase SETDB1 | YESDAPCSSDSSG | 5000 | R | K.EGYESDAPC#SSDSSGVDLK.D | 4 | 9.73294218 | 10.39063026 | 10.41475016 | 10.45205187 | 10.69661648 | 11.0104557 | **0.949600572** |
| SETDB1 | 753 | Q15047 | Histone-lysine N-methyltransferase SETDB1 | TIQATACTPGGQI | 434 | R | K.CACHQLTIQATAC#TPGGQINPNSGYQYK.R | 4 | 9.618806929 | 9.637234771 | 11.38156163 | 10.89822566 | 11.23782446 | 11.26230717 | **0.91733863** |
| SETDB1 | 634 | Q15047 | Histone-lysine N-methyltransferase SETDB1 | KTPCGLCLRTMQE | 5000 | R | K.TPC#GLC#LR.T | 3 | 8.346620676 | 8.235053483 | 9.565492267 | 8.734781537 | 8.825894746 | 8.249464245 | **1.013057887** |
| SETDB1 | 830 | Q15047 | Histone-lysine N-methyltransferase SETDB1 | AKGSFVCIYAGKI | 5000 | R | K.GSFVC#IYAGK.I | 3 | 9.166606848 | 8.581447717 | 9.799886743 | 9.741397416 | 10.47171642 | 9.883632825 | **0.915312928** |
| SETDB1 | 1281 | Q15047 | Histone-lysine N-methyltransferase SETDB1 | KELLCCCGAIECR | 76 | R | K.ELLCCC#GAIECR.G | 3 | 7.517963334 | 8.455586114 | 9.251699774 | 9.276339009 | 8.776229132 | 8.65602962 | **0.944461759** |
| SETDB1 | 631 | Q15047 | Histone-lysine N-methyltransferase SETDB1 | VIYKTPCGLCLRT | 5000 | R | K.TPC#GLC#LR.T | 2 | 9.517943481 | 9.632194546 | 9.951012462 | 11.14522332 | 10.81906525 | 10.74151494 | **0.889785523** |
| SETDB1 | 731 | Q15047 | Histone-lysine N-methyltransferase SETDB1 | FLVGCDCKDGCRD | 104 | R | K.GVFINTGPEFLVGCDC#K.D | 2 | 8.531805392 | 8.053124491 | 8.997989086 | 8.276180702 | 8.739713473 | 9.174464798 | **0.976806732** |
| SETDB1 | 582 | Q15047 | Histone-lysine N-methyltransferase SETDB1 | HVCSYTCLSRVRP | 78 | R | K.LFYLPHVCSYTC#LSR.V | 2 | 7.77146948 | 6.965090034 | 8.666341861 | 6.649896095 | 7.798315651 | 7.72274865 | **1.055565522** |
| SETDB1 | 1280 | Q15047 | Histone-lysine N-methyltransferase SETDB1 | GKELLCCCGAIEC | 32 | R | K.ELLCC#CGAIECR.G | 2 | 8.27917378 | 7.914782314 | 9.176674953 | 8.670520231 | 9.13973664 | 8.848223467 | **0.951690821** |
| SETDB1 | 329 | Q15047 | Histone-lysine N-methyltransferase SETDB1 | DIEDISCRDFIEE | 5000 | R | K.TWEDIEDISC#R.D | 1 | 8.583336772 | 8.859819255 | 8.484298271 | 7.544808044 | 6.861167279 | 6.764879847 | **1.224676759** |
| SETDB1 | 693 | Q15047 | Histone-lysine N-methyltransferase SETDB1 | EDVPLSCVNEIDT | 5000 | R | K.EDVPLSC#VNEIDTTPPPQVAYSK.E | 1 | 8.397421213 | 6.856039533 | 9.019210001 | 6.424059954 | 8.531596688 | 8.757404195 | **1.023599227** |
| SETDB1 | 578 | Q15047 | Histone-lysine N-methyltransferase SETDB1 | FYLPHVCSYTCLS | 79 | R | K.LFYLPHVC#SYTCLSR.V | 1 | 9.668985687 | 8.849204591 | 10.25095641 | 12.62462888 | 11.23321664 | 11.17117916 | **0.821294539** |

**Table S2.** Cysteine-targeted activity-based protein profiling data for the detected 23 cysteines of SETDB1.
